# The Nonsense-Mediated mRNA Decay pathway degrades dendritically-targeted mRNAs to regulate long-term potentiation and cognitive function

**DOI:** 10.1101/389585

**Authors:** Michael Notaras, Megan Allen, Francesco Longo, Nicole Volk, Miklos Toth, Noo Li Jeon, Eric Klann, Dilek Colak

## Abstract

Synaptic plasticity relies on new protein synthesis in dendrites that involves the selective translation of specific mRNAs. This requires a tight control of mRNA levels in dendrites. Consistently, RNA translation and degradation pathways have been recently linked to neurodevelopmental and neuropsychiatric diseases, suggesting a role for RNA regulation in synaptic plasticity and cognition. Despite being the only RNA regulatory pathway that is associated with multiple mental illnesses, the Nonsense-Mediated mRNA Decay (NMD) pathway presents an unexplored regulatory mechanism for synaptic function and plasticity. NMD is a highly conserved and selective RNA degradation pathway that exerts its function in a cell- and spatiotemporally-specific manner. Here, we show that neuron-specific disruption of NMD in adulthood attenuates learning, memory, hippocampal LTP, and potentiates perseverative/repetitive behavior. While it is known that local translation of specific mRNAs in dendrites enables synaptic plasticity, the tightly-controlled mechanisms that regulate local quantity of specific mRNAs remains poorly understood. We report that the NMD pathway operates within dendrites to regulate GluR1 surface levels. Specifically, NMD modulates the internalization of GluR1 and promotes its local synthesis in dendrites. We identified AMPK as a mechanistic substrate for NMD that contributes to the NMD-mediated regulation of GluR1 by limiting total GluR1 levels. These data establish that NMD regulates synaptic plasticity, cognition, and local protein synthesis in dendrites, providing fundamental insight into the neuron-specific function of NMD within the brain.

## Introduction

NMD is a process that regulates protein expression though degradation of specific mRNAs [1–3]. NMD was first identified as an mRNA surveillance pathway that degrades mRNAs undergoing premature termination of translation [4–6]. These mRNAs are detected during the first round of translation. In most mRNAs, all exon-junction complexes (EJCs), a group of proteins bound to mRNA after splicing, are upstream of the termination codon. When a ribosome encounters an EJC downstream of a termination codon, the termination codon is perceived as premature, and the mRNA is considered nonsense. This initiates NMD, recruiting core Up-Frameshift (UPF) proteins - namely UPF1, UPF2, and UPF3B - and ultimately leads to mRNA degradation [7, 8]. NMD thus prevents the production of truncated proteins with a dominant-negative function. Recently, it has been shown that NMD also targets physiological mRNAs that naturally have NMD-inducing features but code for functional proteins [9, 10]. A common event that leads to NMD is alternative splicing, since alternatively spliced exons or introns typically create in-frame termination codons followed by EJCs [11, 12]. Intra-axonal translation of such an mRNA coupled to NMD controls a switch in receptor expression and thereby regulates axon guidance [11]; suggesting that mRNA turnover is a key player in local protein synthesis. Intron retention in the 3’ UTRs of endogenously expressed transcripts also targets them to NMD [13]. For instance, the mRNA of the synaptic plasticity protein Arc is a known target of NMD due to intron retention in its 3’ UTR [13]. Arc is highly expressed in dendrites [14–17] and is required for synaptic plasticity and memory consolidation [18–21]. Similar to *Arc* mRNA, various mRNAs with intron retentions are localized to dendrites [22] and have been implicated in synaptic plasticity and memory [19, 23–26]. However, the extent to which NMD is physiologically relevant to synaptic plasticity is not clear.

Despite its putative link to many mental disorders including autism and schizophrenia [27–32], whether NMD is relevant to the synaptic pathology and behavioral deficits of these diseases remains unclear. A recent study showed that *UPF3B*-null mice exhibit deficits in fear-conditioned learning, supporting a role for NMD in regulating synaptic plasticity and cognition [33]. However, because NMD is ubiquitous and required for neuronal differentiation [34], disruption of this pathway during development hinders determining the direct role of NMD in these processes. Therefore, understanding the precise, cell-type specific role of NMD in developing and adult brain is likely to provide novel insights into key mechanisms critical for synaptic plasticity and behavior.

Synaptic plasticity underlies changes in neuronal network dynamics and is therefore considered to be the foundation of learning and memory. It takes several forms, including the modification of synaptic strength and spine structure. Synaptic transmission can be influenced by activity, becoming either enhanced, through long-term potentiation (LTP), or depressed, through long-term depression (LTD) [35]. Many higher cognitive disorders are associated with alterations in spine density and synaptic plasticity [36, 37]. Mechanisms that regulate synaptic plasticity are therefore fundamental to understanding higher brain functions, including learning and memory, and have been implicated in the etiology of neuropsychiatric diseases. Both LTP and LTD rely on local protein synthesis in dendrites [21, 38]. Locally synthesized proteins are thought to either strengthen a pre-existing synaptic connection, by inducing structural remodeling, or to promote the formation of new synaptic connections. Deficits in mRNA regulation can result in ectopic protein synthesis and lead to cognitive disabilities [39–45], having downstream effects on synaptic plasticity and higher-order cognition. For example, exaggerated mRNA translation leads to alterations in spine density, LTP and LTD [42, 45–47] suggesting that a tight control of protein synthesis is required at many levels within dendrites [40]. A great amount of prior research has addressed the pathways that regulate translational derepression in dendrites. However, the mechanisms that control mRNA quantity during synaptic function remain unknown. Importantly, it is not known whether NMD functions locally to regulate mRNA levels in dendrites, and if mRNA degradation contributes to the regulation of synaptic plasticity.

Here we show that both synaptic plasticity and cognitive function is physiologically regulated by neuronal NMD. Conditional deletion of the NMD component *UPF2* in adult glutamatergic neurons *in vivo* disrupts learning and memory and potentiates perseverative/repetitive behavior. Deletion of *UPF2* also results in altered LTP, but not LTD, in the adult hippocampus. This phenotype is accompanied by a significant reduction in total GluR1 levels in conditional knockout mice. We show that NMD regulates GluR1 surface levels through mechanisms that influences GluR1 internalization and nascent protein synthesis. Selective depletion of UPF2 from cultured hippocampal neurons reduces both total and surface expression of GluR1 without affecting *GluR1* mRNA levels. This phenotype is recapitulated when NMD is disrupted in dendrites via the targeted-inhibition of UP3B local synthesis; the only dendritically synthesized component of the NMD machinery. Disruption of NMD increases internalization of GluR1 in dendrites, which may account for the reduced surface levels of this receptor. This is consistent with previous findings that Arc mediates GluR1 internalization and thereby reduces surface levels of GluR1. However, elevated levels of Arc are, at most, only partially responsible for the reduced surface levels of GluR1 in UPF2-deficient dendrites. In addition to its altered internalization, the local synthesis of GluR1 is also reduced in UPF2-deficient dendrites due to increased levels of *AMPK* mRNA. Normalizing the levels of *AMPK* and *Arc* mRNAs, together, completely corrects GluR1 levels in UPF2-deficient dendrites. These data demonstrate a role for NMD in regulating cognitive function, the dendritic proteome, and synaptic plasticity, as well as potentially other essential dendritic functions.

## Results

### NMD is required for learning and memory and suppresses perseverative behavior

We first sought to determine the requirement for neuronal NMD in cognitive function and synaptic plasticity. To do this, we genetically disrupted NMD in postmitotic glutamatergic neurons using a conditional knockout mouse of *UPF2* (*UPF2*^fl/fl^) [48]. The function of UPF2 is exclusive to the NMD pathway, and this line has been successfully used to target NMD in various tissues [11, 48, 49]. Because NMD is ubiquitous and required for neuronal differentiation [34], deleting the *UPF2* gene during development would preclude determining an explicit role of NMD in adult plasticity and cognition. Therefore, we temporally controlled NMD disruption by crossing the *UPF2^fl/fl^* line with a mouse line that expresses tamoxifen-inducible Cre under the α*CaMKII* promoter (α*CaMKII*:*CreER^T2^*) [50]. In all behavioral experiments, we induced Cre in 2-month old conditional-knockout (*UPF2*^fl/fl^;α*CaMKII*:*CreER^T2^*; CKO) mice as well as in their control littermates (*UPF2*^wt/wt^;α*CaMKII*:*CreER^T2^*; CTRL) by tamoxifen. This experimental design ensures that phenotypes are not caused by an acute effect of tamoxifen, or a non-specific effect of Cre at cryptic LoxP sites. Loss of UPF2 protein in excitatory neurons was successfully achieved by 200mg/kg of tamoxifen administered every other day for 5 doses (Figure S1a). A 2-week washout period followed tamoxifen treatment before plasticity and behavioral analyses began. We first evaluated the general locomotor activity of conditional *UPF2* CKO mice in locomotor photocells [51]. No difference in locomotor activity was detected between *UPF2* CKO mice and controls, indicating that NMD does not induce hypo-or hyperactivity (Figure S1b). Likewise, no evidence of anxiety-related behavior was observed in *UPF2* CKO mice in the elevated-plus maze or open field assays (Figure S1c-d), nor was there a difference in sociability between groups (Figure S1e).

We next examined hippocampus-dependent learning and memory in *UPF2* CKO mice. To do this, we used multiple behavioral paradigms, including the Y-maze, contextual fear conditioning and the Morris water maze. On the Y-maze, *UPF2* CKO mice showed no preference for the novel arm relative to the other arms, and displayed significantly worse novel arm preference than the control group (Figure 1a), suggesting that *UPF2* CKO mice have disrupted short-term spatial memory. As an additional measure of hippocampal function, we tested *UPF2* CKO mice for contextual fear memory. In this test, *UPF2 CKO* mice froze for significantly less time than controls by the last tone-shock pairing during conditioning, and this conditioning deficit resulted in worse contextual fear memory in *UPF2* CKO mice relative to control mice 24 hr later (Figure 1b-c). To further disambiguate whether our observed phenotypes on the Y-maze and contextual fear conditioning represented an acquisition failure as opposed to a memory deficit, we subjected mice to the Morris water maze. In the Morris water maze, *UPF2* CKO mice displayed a robust attenuation of learning behavior across training and correspondingly decreased exploration of the target quadrant and increased latencies to enter the (now removed) hidden platform zone on the probe trial (Figure 1d-f). *UPF2* CKO mice failed in most trials to even attempt to locate the hidden platform despite being guided to its location after each trial. Indeed, the final task performance of these mice did not surpass the performance of control mice after just two days of training. These data collectively suggest that *UPF2* CKO mice have attenuated learning behavior, which produces worse performance on common assays of hippocampus-dependent memory function.

**Figure 1.**
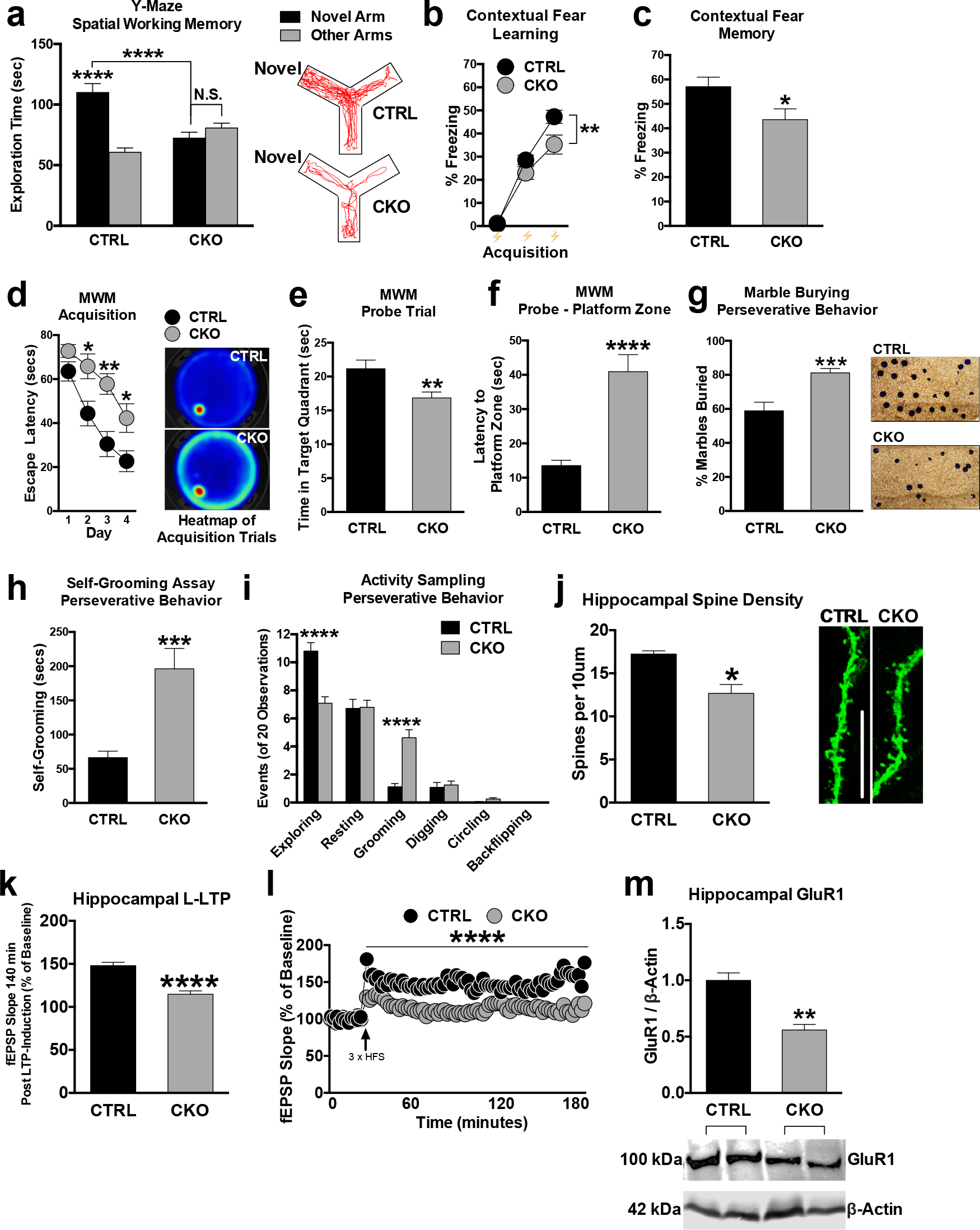
NMD is required for learning, memory, LTP, and suppresses perseverative behavior. To determine the role of neuronal NMD in cognitive function and synaptic plasticity, we targeted NMD in postmitotic neurons using a conditional knockout mouse of *UPF2* (*UPF2*^fl/fl^). We spatially and temporally controlled *UPF2* ablation by crossing *UPF2^fl/fl^* with a mouse line that expresses tamoxifen-inducible Cre under the α*CaMKII* promoter (*UPF2*^fl/fl^;α*CaMKII*:*CreER^T2^*; referred to as CKO). We treated CKO and CTRL (*UPF2*^wt/wt^;α*CaMKII*:*CreER^T2^*) mice with tamoxifen (200 mg/kg) every-other-day for 5 doses from 6 weeks of age. This tamoxifen regimen led to successful depletion of UPF2 protein from postmitotic hippocampal neurons (see Figure S1a). **a-c**, To assess hippocampus-dependent short-term spatial and associative memory, we subjected *UPF2* CKO and CTRL mice to the Y-maze and contextual fear conditioning respectively. In the Y-maze, mice were exposed to 2 of 3 arms of the maze for 10 min, subjected to a 1 hr delay, and then returned to explore all arms of the maze for 5 min. Preference for exploring the previously blocked ‘novel’ arm, relative to the other arms, was quantified as our index of short-term spatial memory. Compared to CTRL mice, which had an intact preference for exploring the novel arm relative to familiar arms of the Y-Maze (n=24), CKO mice displayed a complete lack of preference for the novel arm relative to the familiar arms of the Y-Maze (n=22, **a**). To examine learning (**b**) and hippocampus-dependent associative fear memory (**c**), we adapted a simple 2-day contextual fear memory protocol. Mice were conditioned using 3 tone-shock pairings on day 1, where increased freezing behavior to each successive tone-shock pairing was interpreted as learning. Analysis of freezing across tone-shock pairings revealed that CKO mice (n=22) froze less than CTRL mice (n=24) on the third tone-shock pairing at the end of conditioning, implicating that CKO mice displayed aberrant acquisition (**b**). On day 2, mice were returned to their conditioning context for 6 min. CKO mice (n=22) also displayed lower freezing levels than CTRL mice (n=24), implicating that CKO mice had worse contextual fear memory (**c**). **d-f**, To further confirm learning and memory phenotypes, we also employed the MWM. Mice were trained 4 times each day, for 4 consecutive days, to learn the location of a hidden submerged platform. Trials were averaged across each day of testing and plotted as a time-course, with gradually decreasing escape latencies being interpreted as spatial learning. When comparing the escape latencies to CTRL mice, CKO mice showed delayed escape latencies on days 2, 3, and 4 (n=18 for each group), implicating attenuated spatial learning upon disruption of UPF2 (**d**). Following training in the MWM, mice were subjected to a probe test to examine spatial memory of the hidden platform location. The hidden platform was removed from the MWM, and time spent exploring the quadrant that previously contained the hidden platform was quantified. Compared to CTRL mice, CKO mice spent significantly less time exploring the target quadrant that previously contained the hidden platform (n=18 per group), implicating a spatial-memory deficit (**e**). During the probe trial of the MWM, we also measured latency to enter the previously hidden platform zone as an additional measure of spatial memory. Consistent with quadrant search times, CKO mice took significantly longer to enter the hidden platform zone compared to CTRL mice supporting a spatial memory deficit in CKO mice (**f**). Together, these data suggest that disruption of UPF2 in hippocampal excitatory neurons leads to attenuated learning, and altered hippocampus-dependent memory. **g-i**, We also tested the *UPF2* CKO mice for repetitive and compulsive-like behavior. In the widely-adapted marble-burying test, we quantified the number of marbles buried as a % of total after a 20 min exposure period. CKO mice buried significantly more marbles than CTRL mice (58.86% versus 81.14% of marbles; n=25 per group), which in lieu of an anxiety-like phenotype in CKO mice (see Figure S1), is suggestive of an induced perseverative behavioral phenotype in *UPF2* CKO mice (**g**). As an additional measure, we also monitored self-grooming behavior over a 20 min observation period. CKO mice (n=24) groomed for significantly longer than CTRL mice (n=25), supporting the repetitive behavior seen in the marble burying test (**h**). During the self-grooming assay, we also monitored if CKO mice displayed any additional behavioral traits reminiscent of compulsive-like behavior. Apart from an increase in the number of grooming bouts, CKO mice showed no further compulsive-like traits when compared to CTRL mice (**i**). These data suggest that NMD in excitatory neurons is required to suppress perseverative behavior. **j**-**l**, NMD regulates spine density and L-LTP in hippocampus. Synaptic plasticity is tightly linked to spine density. We therefore sought to examine whether disruption of UPF2 alters spine density in hippocampus. To visualize dendrites and spines in CTRL and CKO mice, we stereotaxically injected AAV-CaMKIIa-eGFP virus bilaterally into the CA1 subfields of the hippocampus. To perform spine quantification, we cut CTRL and CKO brains at 200 μm and acquired images by creating maximum intensity projections from Z-stacks using up to ~30 optical sections (~0.5 μm, Olympus Fluoview). CKO mice had significantly fewer spines on apical dendrites of hippocampal CA1 neurons compared to CTRL mice (n=3 mice per group; 4-10 independent apical dendrite segments, acquired from up to 5 distinct fields per mouse) (**j**). To measure LTP, hippocampal slices (400 μm) were prepared at approximately 10 weeks of age. To induce LTP, we used either one (E-LTP) or three (L-LTP) 1 sec 100-Hz high-frequency stimulation trains, with an intertrain interval of 60 sec for the L-LTP experiments. Although E-LTP was not altered in CKO mice, when fEPSPs were compared to CTRL mice, we found that L-LTP was disrupted in CKO mice (n=12 hippocampal slices from 7 CTRL mice, n=14 slices from 8 CKO mice) (**k**and **l**). Similar to E-LTP, LTD was unaltered in the CKO hippocampus (Figure S1f-g). Together, these data indicate that NMD is required for proper synaptic plasticity and spine density in the adult hippocampus. **m**, GluR1 protein levels are decreased in *UPF2* CKO hippocampus. Disruption of EJC proteins, which are instrumental in directing selected mRNAs towards NMD, has been shown to result in increased GluR1 levels in dendrites. While this suggests that NMD may regulate GluR1 levels, it is also reported that EJC proteins are not specific to the NMD pathway. We therefore measured endogenous GluR1 levels in the hippocampus of *UPF2* CKO mice via western blot. The *UPF2* CKO mice had substantially decreased GluR1 expression relative to CTRL mice. This data indicates that the disruption of UPF2 results in a significant down-regulation of GluR1 levels. Data are represented as mean ± SEM; *p < 0.05, **p < 0.01, ***p <0.001 and ****p < 0.0001. N.S. represents “Not Significant”. Scale bar: 5 μm.

Because the NMD pathway has been previously linked to autism and schizophrenia [27–32], where stereotypies are common to both disorders, we also tested for perseverative/repetitive behaviors [45, 52–55]. This primarily included an assessment of marble burying and self-grooming. In our marble burying assay, *UPF2* CKO mice buried significantly more marbles than control mice (Figure 1g; 81.14% vs 58.86% of marbles). In an activity-sampling test session, where grooming behavior was continuously monitored for 20 min, CKO mice groomed for longer and displayed a greater frequency of grooming bouts compared to the control group (Figure 1h-i). Other stereotypies such as digging, circling, and backflipping were not significantly different between the genotypes (Figure 1i). Together, these data suggest that NMD is required for proper learning and memory, and its disruption potentiates perseverative/repetitive behaviors.

### NMD ensures proper spine density and L-LTP in hippocampus

We next examined spine density in *UPF2* CKO mice. During LTP, activation of glutamate receptors increases connectivity of specific neurons through the enlargement of pre-existing spines or the formation of new spines, which supports learning and memory. To determine whether NMD is required for proper spine density, we quantified spine density in hippocampal dendrites of both *UPF2* CKO and control brains cut at 200 μm. To visualize dendrites and spines, we injected AAV-αCaMKII-eGFP virus into the dorsal CA1 field 7d prior to perfusion. We observed significantly reduced CA1 spine density in *UPF2* CKO hippocampal dendrites compared to control dendrites *in vivo* (Figure 1j). We next sought to investigate the role of NMD in well-studied forms of synaptic plasticity in the adult brain by assessing both LTD and LTP in these mice. To do this, we prepared hippocampal slices from 3-month old *UPF2* CKO mice and their control littermates following induction of Cre expression by tamoxifen at 2-month of age. We incubated 400 μm slices for at least 3 hr in artificial cerebral spinal fluid (ACSF) before measuring LTP. LTP can be divided into two phases: early phase LTP (E-LTP) and late phase LTP (L-LTP). E-LTP does not require new gene transcription or mRNA translation, whereas L-LTP requires new gene transcription and mRNA translation and results in an increase in the number of AMPA receptors accompanied by an increase in the size of the synaptic connection. Briefly, we recorded field excitatory postsynaptic potentials (fEPSPs) from the stratum radiatum and activated Schaffer collateral/commissural afferents at 0.05 Hz. We used a single 100 Hz high-frequency stimulation (HFS) to induce E-LTP and three trains of HFS with a 60 sec intertrain interval to induce L-LTP. We found that L-LTP, but not E-LTP, was significantly decreased in UPF2-deficient slices compared to control slices (Figure 1k-l). On the other hand, LTD, which was induced by application of 100 μM DHPG for 10 min, was not different in UPF2-deficient slices compared to control slices (Figure S1f-g). These data suggest that NMD is required for proper LTP and spine density but not for LTD in hippocampus.

### NMD positively regulates GluR1 surface expression

We next addressed the potential mechanisms through which NMD regulates synaptic plasticity and cognitive function. It was previously shown that disruption of the EJC, an essential component for targeting mRNAs to the NMD pathway, causes an increase in the levels of AMPA receptor GluR1 in hippocampal dendrites *in vitro* [13]. Functionally, AMPA receptors mediate the vast majority of excitatory synaptic transmission by ensuring rapid responses to glutamate, the principal neurotransmitter in the mammalian central nervous system. Modulation of AMPA-receptor integrity plays a crucial role in synaptic strength. Strengthening of synapses by LTP or weakening of synaptic strength by LTD can be represented by the synaptic insertion or removal of AMPA receptors, respectively [56]. For example, low surface expression of AMPA receptor GluR1 increases the number of dendritic spines [57]. Similarly, reductions in GluR1 expression cause working-memory deficits in mice [58]. If EJC function were specific to the NMD pathway, the EJC disruption study would suggest that NMD negatively regulates GluR1 levels; but EJC proteins have multiple functions, including transportation and translation of mRNAs [59]. Intriguingly, contrary to what is observed in studies of disrupted EJC proteins, we found that the loss of *UPF2* led to a decrease in GluR1 levels *in vivo* (Figure 1m). To explore this phenotype *in vitro*, we infected hippocampal neurons with *UPF2*-shRNA:GFP lentivirus to interrupt endogenous NMD activity at 7 days *in vitro* (DIV7) and examined GluR1 expression on DIV21. We tested 2 *UPF2*-shRNA lentiviruses. Infection of hippocampal neurons with either *UPF2*-shRNA 1 or *UPF2*-shRNA 2 led to a nearly complete loss of UPF2 protein in these neurons (Figure S2). We measured the total levels of GluR1 using an anti-Cterminus-GluR1 antibody in permeabilized cells and found that total levels of GluR1 were decreased both in cell bodies and dendrites of hippocampal neurons upon knockdown of *UPF2* (Figure S3a-c). Thus, the reduction in GluR1 observed in *UPF2* CKO mouse hippocampus was recapitulated in neurons and their dendrites *in vitro*.

We next asked whether NMD regulates GluR1 surface expression in neurons. An essential component of GluR1 signaling involves the dynamic insertion and removal of the GluR1 receptor to the synaptic surface [60]. To label surface GluR1, we incubated fixed, but not permeabilized, neurons with a monoclonal anti-Nterminus-GluR1 antibody at 4°C as described previously [61, 62]. We found that, similar to the total levels, the loss of UPF2 led to a decrease in the surface levels of GluR1 (Figure 2; see also Figure S3c and S4 for *UPF2* shRNA 2 knockdown and surface GluR1/PSD-95 quantifications, respectively). This suggests that NMD positively regulates GluR1 surface expression. *GluR1* mRNA is not a canonical target of NMD. However, it is known that some mRNAs that do not carry any NMD-inducing features can also be degraded by NMD [63]. In addition, it has been suggested that GluR1 transcription is regulated by Arc [64]. Consistent with *Arc* mRNA being an established target of NMD [11, 13], the levels of both *Arc* mRNA and Arc protein are increased in hippocampal neurons upon knockdown of *UPF2* (Figure S5). We, therefore, examined cytoplasmic *GluR1* mRNA levels in UPF2-deficient cell bodies. Overall *GluR1* mRNA levels were unaltered following knockdown of *UPF2* in hippocampal neurons, indicating that the reduction in total and surface GluR1 levels is not due to low levels of its mRNA (see Figure S7 for viral infection efficiency in hippocampal cultures). These data suggest that NMD modulates GluR1 signaling by restraining a mechanism that negatively regulates the synthesis, surface localization or surface turnover of GluR1 in dendrites.

**Figure 2.**
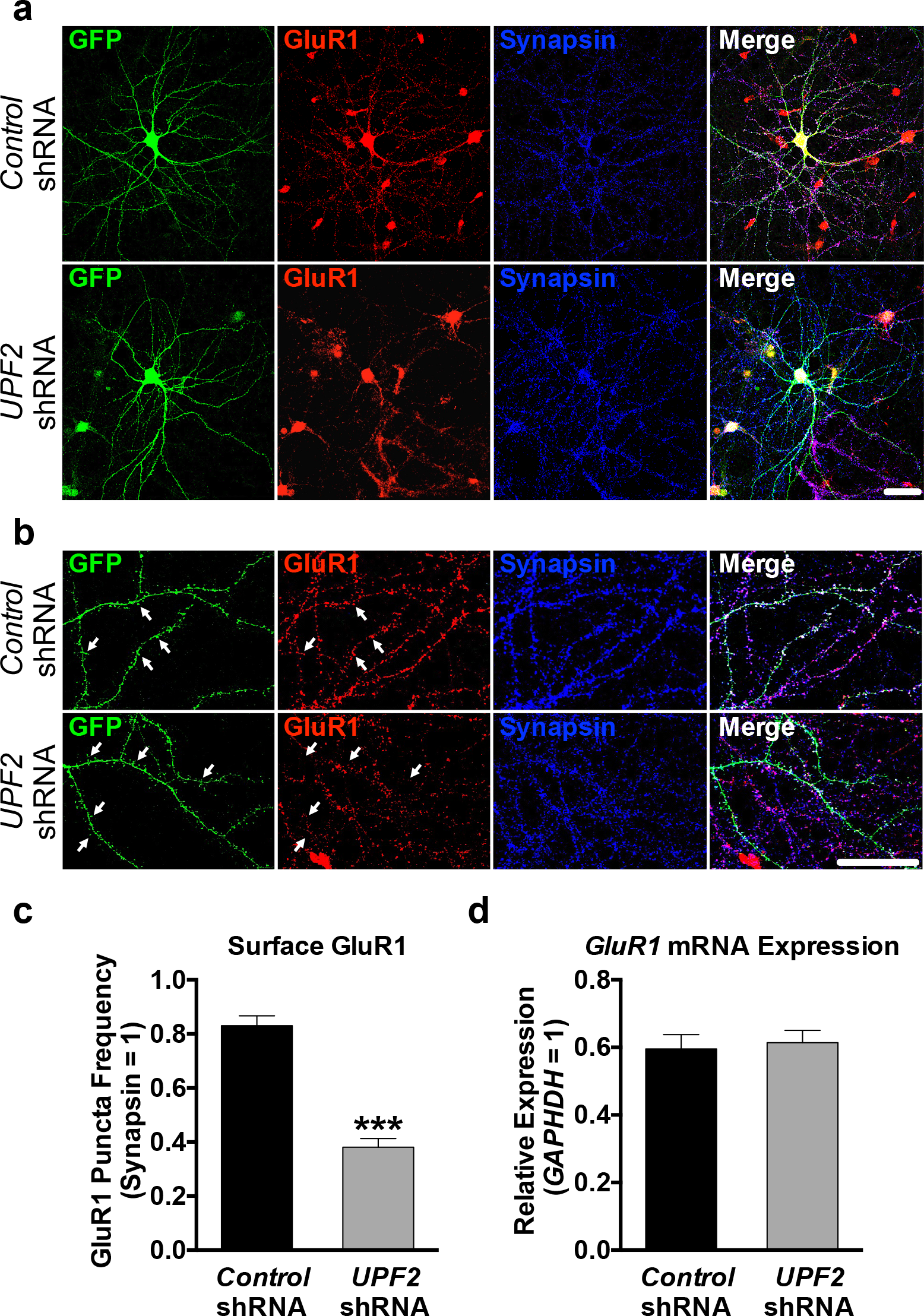
NMD is required for proper surface expression of GluR1. To study how NMD regulates GluR1 expression, we cultured mouse hippocampal neurons at E16 and infected with a *control*- or *UPF2*-shRNA lentivirus at DIV7, which led to a robust and global depletion of UPF2 in these neurons (2 independent shRNA constructs; see Figure S2). The reduction in GluR1 observed in *UPF2* CKO mouse hippocampus was recapitulated in neuron cell bodies and dendrites upon knockdown of *UPF2 in vitro* (Figure S3a-b). Dynamic insertion and removal of the GluR1 receptor to the synaptic surface plays an important role in GluR1 signaling and as a result in synaptic strength (49). We next asked whether NMD regulates GluR1 surface expression in neurons. **a-b**, Hippocampal neurons infected with the *control*- or *UPF2-shRNA* 1 lentivirus (referred to as *UPF2-shRNA* throughout the text) at DIV7 were fixed at DIV21. To detect surface GluR1 levels, fixed but not permeabilized neurons were stained with an anti-N-terminus-GluR1 antibody. Following this, we permeabilized neurons and costained for the synaptic vesicle marker, synapsin. The cells were visualized with GFP endogenously expressed in infected neurons by the lentiviruses applied. Representative examples of low and high magnification of surface GluR1 staining in GFP-positive neurons shown in (**a**) and (**b**), respectively. Arrows show examples of positive and negative GluR1 signal at spines. **c**, Quantifications of surface GluR1 puncta revealed that neurons with disrupted NMD had approximately 50% lower surface GluR1 frequency in dendrites, compared to neurons infected with control virus (n=3 biological replicates per group; 10 neurons and 24 dendrites per group; see also Figure S3c for the *UPF2* shRNA 2 data). Knockdown of *UPF2* did not change synapsin density indicating that decreased GluR1 levels do not arise from an alteration in the synaptic potential of UPF2-deficient neurons. Additionally, this phenotype was recapitulated when PSD-95 was utilized as an alternative synaptic marker (Figure S4). **d**, To determine whether a reduction in *GluR1* mRNA levels accounts for the decrease in surface GluR1 protein, we prepared cDNA from control and UPF2-deficient neurons at DIV21 and performed qRT-PCR for *GluR1* mRNA. qRT-PCR data revealed no significant difference in *GluR1* mRNA levels between *control*- and *UPF2*-shRNA infected neurons (n=3 replicates per group; with more than 90% of infection efficiency in each replicate. See Figure S7 for the high efficiency of neuronal infection by *UPF2* shRNA virus). This data suggests that alteration in *GluR1* transcription or degradation is not responsible for the loss of surface GluR1 protein in UPF2-deficient dendrites Data are represented as mean ± Standard Error of the Mean (SEM); **p < 0.01. Scale bar: 20 μm.

### NMD negatively regulates GluR1 internalization in dendrites

We next sought to determine how NMD regulates surface expression of GluR1 in dendrites. It is possible that the reduction in total GluR1 levels (Figure 1m and Figure S3a-b) might account for decreased surface levels of this receptor. However, locally synthesized Arc is also known to cause GluR1 endocytosis, leading to a decrease in the surface levels of this receptor [21, 65]. This suggests that increased levels of Arc might also contribute to downregulated surface-GluR1 levels upon loss of UPF2. To determine if GluR1 endocytosis is altered after disruption of NMD, we measured the basal internalization rate of GluR1 upon knockdown of *UPF2* in hippocampal dendrites using a previously established protocol [66]. We observed significantly increased GluR1 basal internalization in UPF2-deficient hippocampal dendrites (Figure 3a-b). This is consistent with the increased levels of Arc in these neurons, namely because Arc-mediated internalization of GluR1 only influences surface but not total levels of GluR1 [67]. This suggests that UPF2 regulates total GluR1 levels via yet defined NMD targets.

**Figure 3.**
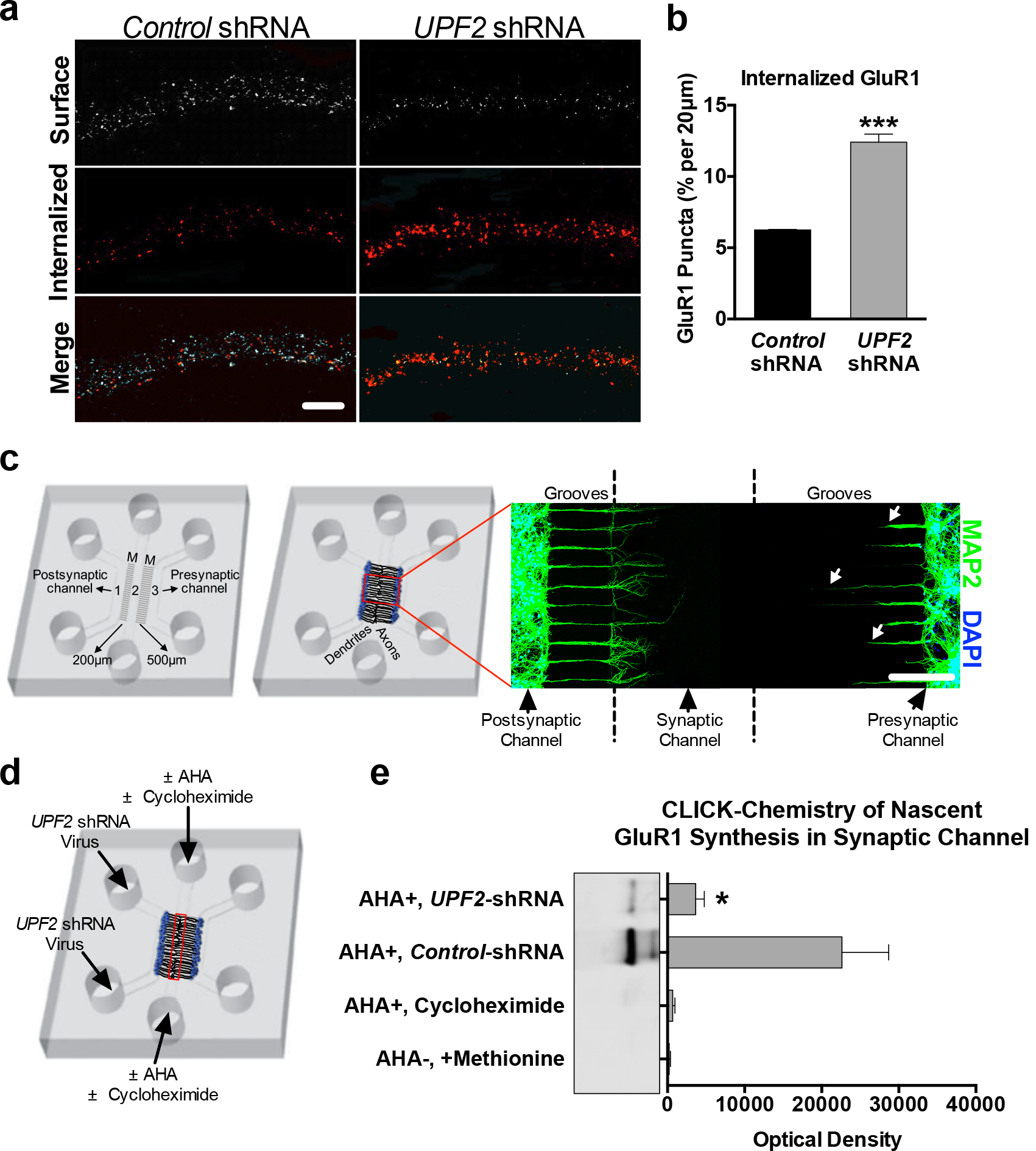
NMD regulates GluR1 internalization and local dendritic synthesis. **a**, It is possible that the reduction in total GluR1 levels (Figure 1m and Figure S3a-b) might account for decreased surface levels of this receptor. However, locally synthesized Arc is also known to cause GluR1 endocytosis, leading to a decrease in the surface levels of this receptor [21, 61]. The levels of both *Arc* mRNA and Arc protein are increased in hippocampal dendrites upon knockdown of *UPF2* (Figure S5). Because Arc drives the removal of surface GluR1 from synapses [21, 61], we examined the possibility that GluR1 internalization may occur at an increased rate in UPF2-deficient dendrites due to increased Arc levels. To do this, we measured basal internalization rates of GluR1 receptor at DIV 21. We pre-incubated live cells with anti-N-terminus GluR1 antibody for 5 min, fixed, and labeled with Alexa-546 secondary to allow surface labeling of GluR1 (see Methods). Percent internalization showed that neurons with disrupted UPF2 internalized approximately 50% more GluR1 compared to neurons infected with *control*-shRNA lentivirus (6.25% vs. 12.41% internalized GluR1; n=3 biological replicates per *control*-shRNA [13 neurons, 33 dendrites] and per *UPF2*-shRNA [20 neurons, 53 dendrites]). **b**, Local regulation of GluR1 degradation and synthesis play a prominent role in GluR1 levels and signaling (51, 52). Therefore, in addition to the reduction in global GluR1 protein levels, an alteration in the homeostatic balance of local synthesis and local degradation of GluR1 protein is likely to be responsible for the reduction in local GluR1 levels observed upon knockdown of *UPF2*. To study the local regulation of GluR1 protein levels, we used a custom microfluidic device (see Methods) that allows selective access to synaptic compartments *in vitro*. The presented schematic details the microfluidic device that contains three channels separated by microgrooves (M). The separation of channels by microgrooves creates both fluidic and physical isolation between channels. When neurons are cultured in channels 1 and 3, channel 2 (i.e., the middle “synaptic” channel) becomes enriched with dendrites and axons. As the microgrooves that separate channels 1 and 3 from the middle channel are of different lengths, dendrites of neurons cultured only in channel 1 (postsynaptic) can extend to the synaptic channel located in the middle of the device. At DIV8, immunostaining for the dendrite marker MAP2 (see axonal projection in Figure S6) showed that the synaptic channel is exclusively populated with dendrites emerging from the postsynaptic channel. Arrows show that MAP2-positive dendrites from channel 3, or presynaptic channel, do not reach the synaptic channel. **c**, To examine local synthesis of GluR1 protein in UPF2-deficient dendrites, we cultured E16 hippocampal neurons in tripartite chambers and applied *control*- or *UPF2*-shRNA virus to postsynaptic cell body channels at DIV7. This resulted in loss of UPF2 protein only in postsynaptic cell bodies and dendrites but not in presynaptic cells (Figure S7). To measure the dendritic synthesis of GluR1, we used a CLICK-chemistry approach that labeled only locally synthesized proteins at synapses. At DIV21, we exchanged the growth medium selectively within synaptic channels to methionine-free medium containing the methionine analog azidohomoalanine (AHA, 25 μM). After harvesting material from the synaptic channels (red box), we incubated lysates with DBCO-biotin (30 μM) for 1 hr at ambient temperature. We purified biotin-containing proteins over streptavidin columns, separated by SDS-PAGE, and detected nascent GluR1 with an anti-C-terminus antibody. The dendrites of UPF2-deficient neurons exhibited a significant reduction in newly synthesized local GluR1 levels compared to control cultures. This suggests that the decreased rate of locally synthesized GluR1in dendrites contributes to the overall reduction in total, as well as local, GluR1 levels upon loss of UPF2 protein. Data are represented as mean ± SEM; **p < 0.01, and ***p < 0.001. Scale bar: **a** 20 μm, **b** 200 μm.

### NMD positively regulates local synthesis of GluR1 in dendrites

An overall reduction in GluR1 protein levels likely contributes to the decreased dendritic GluR1 levels upon loss of UPF2 in hippocampal neurons (Figure S3). However, the total levels of dendritic GluR1 are determined by the amount of GluR1 protein trafficked to dendrites from cell bodies as well as a balance of local protein synthesis [68] and local protein degradation [69, 70] in dendrites. Thus, we also investigated the possibility of enhanced GluR1 proteolysis or decreased GluR1 synthesis in UPF2-deficient dendrites. To do this, we used a custom-fabricated tripartite microfluidic device, which, unlike traditional microfluidic chambers with two channels [71], contains three channels (Figure 3c and Figure S6-7). As each channel is fluidically isolated, tripartite chambers permit the selective treatment or harvesting of discrete neuronal populations or compartments. In order to access a high number of dendritic synapses, a ‘synaptic’ channel was placed 200 μm from the edge of one channel and 500 μm away from the other. Because the average length of dendrites extending into the microgrooves is less than 300 μm at DIV21, the closer “postsynaptic cell channel” allows a large number of dendrites to project from this channel to the “synaptic” channel (Figure 3c). This enabled us to selectively isolate, treat and harvest dendrites in subsequent experiments.

When AMPA receptors are internalized, endocytosed receptors undergo lysosome- or proteasome-mediated degradation [70, 72–75]. To study local degradation of GluR1, we cultured E16 hippocampal neurons in tripartite chambers and applied control- or *UPF2*-shRNA viruses only to postsynaptic cell channels at DIV7. This resulted in disruption of NMD only in postsynaptic cell bodies and dendrites but not in presynaptic cells (Figure S7). At DIV21, we inhibited both proteosomal and lysosomal degradation by treating the synaptic channels with MG132 and leupeptin. Following 6 hr of treatment, we measured the surface levels of GluR1 in control and UPF2-deficient dendrites in the absence and presence of degradation inhibitors. Co-inhibition of lysosome- and proteasome-mediated degradation did not restore GluR1 levels in UPF2-deficient dendrites (Figure S8). Thus, degradation of GluR1 receptors at the protein-level does not contribute to the reduction in surface GluR1 observed in UPF2-deficient dendrites.

We next asked whether local synthesis of the GluR1 receptor is altered in NMD-deficient dendrites. To measure the dendritic synthesis of GluR1, we used a CLICK-chemistry based approach and labeled locally synthesized nascent proteins at synapses (Figure 3c-e). We cultured E16 hippocampal neurons in tripartite chambers and targeted NMD only in postsynaptic cells by applying the *UPF2*-shRNA virus into the postsynaptic cell channel at DIV7. At DIV21, we exchanged the growth medium in the synaptic channels to methionine-free medium containing the methionine analog azidohomoalanine (AHA) [76] to label all newly synthesized proteins at synapses (Figure 3d). After harvesting the material from the synaptic channels, we incubated lysates with DBCO-biotin [77] to label newly synthesized proteins with biotin. We purified biotin-containing proteins over streptavidin columns, separated by SDS-PAGE, and detected protein by immunoblotting with an anti-Cterminus-GluR1 antibody. The amount of GluR1 protein that bound the streptavidin in the UPF2-deficient synapses was significantly lower compared to the amount of GluR1 that bound the streptavidin in the control synapses (Figure 3e). This data indicates that dendritic synthesis of GluR1 is reduced upon disruption of UPF2.

### Local degradation of *Arc* and *AMPK* mRNAs by NMD in dendrites

We next sought to determine whether local NMD contributes to the regulation of GluR1 protein synthesis in dendrites. Similar to *Arc* mRNA, the mRNA of AMP-activated protein kinase (AMPK) also contains an intron retention in its *3’UTR* [13], making this mRNA a potential substrate for NMD. Activated AMPK interrupts translation by either inhibiting the mammalian target of rapamycin (mTOR) kinase or the translation elongation factor 2 [78–83]. Interestingly, AMPK has been suggested to negatively regulate GluR1 protein levels in dendrites [25]. The *AMPK* mRNA that carries NMD-inducing feature is transcribed from the *PRKAG3* gene and encodes the non-catalytic subunit of the AMPK heterodimer protein. Intriguingly, both *Arc* and *AMPK* mRNAs are localized to dendrites, suggesting that they might be degraded by NMD in these compartments.

To explore this possibility, we first asked whether NMD functions locally in dendrites. NMD occurs in axonal growth cones and influences the local proteome by regulating mRNA dynamics [11]. Similar to growth cones, we found that the core proteins of the NMD machinery are localized to dendrites (Figure 4a), suggesting that local NMD might contribute to regulation of GluR1 levels in these compartments. We first examined *Arc* and *AMPK* mRNA levels in UPF2-deficient synaptic regions. To do this, at DIV7, we selectively infected postsynaptic neurons with *UPF2*-shRNA lentivirus and isolated RNA from synaptic channels at DIV21. qRT-PCR showed an increase in the levels of both *Arc* and *AMPK* mRNAs in UPF2-deficient synaptic channels compared to control synaptic channels, suggesting that these targets are subjected to NMD in dendrites (Figure 4b).

**Figure 4.**
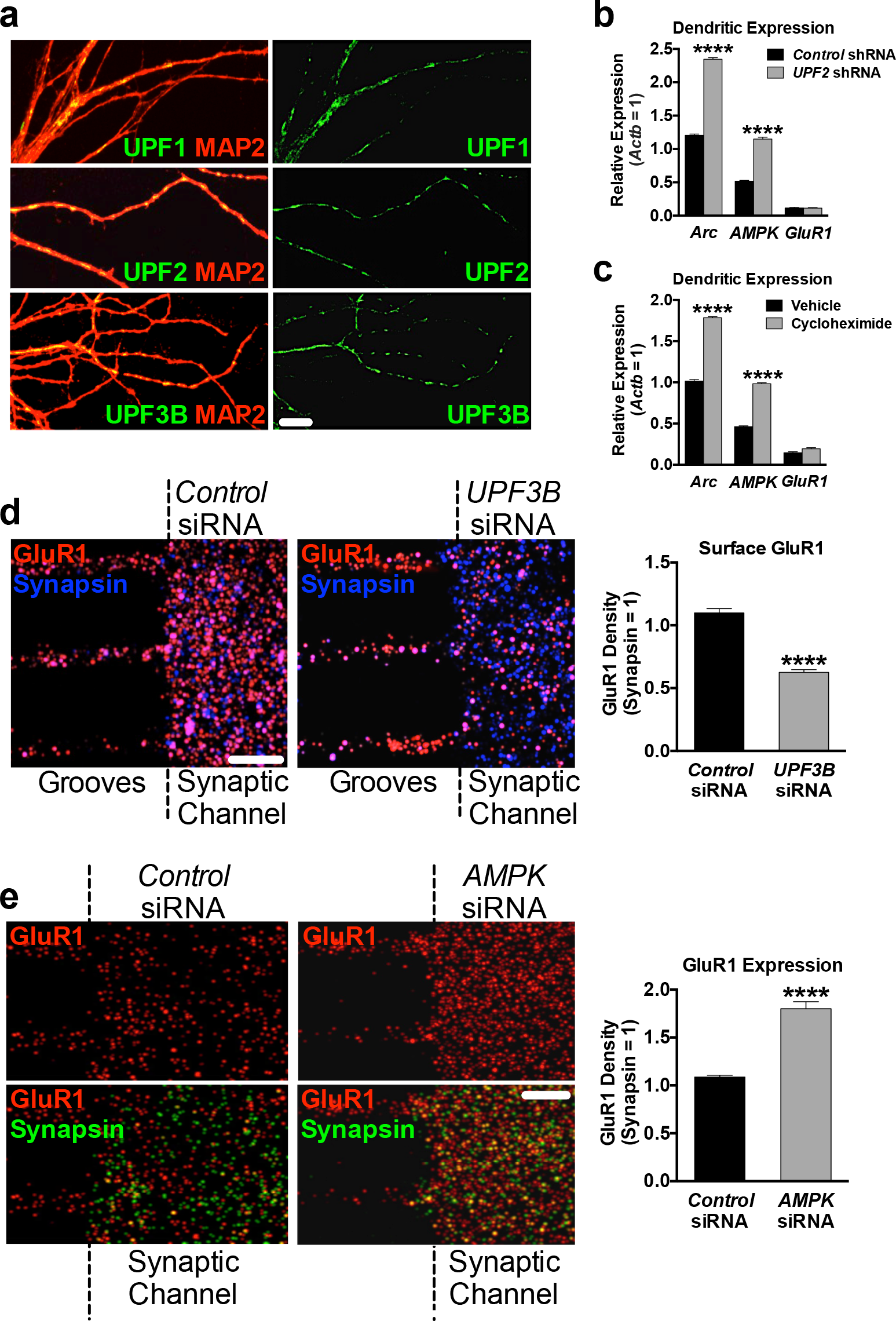
Local regulation of GluR1 levels by NMD. Dendritic GluR1 phenotypes seen upon knockdown of *UPF2* in postsynaptic cells (Figure 3) may arise from NMD targets that are affected in the cell body but act locally within dendrites. However, it is also possible that NMD might occur locally within dendrites to degrade its targets, which in turn regulates the local dendritic proteome and thus GluR1 expression. **a**, The NMD machinery is localized to dendrites. We first examined whether the major proteins of the NMD machinery localize to dendrites. We cultured E16 hippocampal neurons in the cell-body channels of tripartite microfluidic devices, and at DIV21, we fixed and stained synaptic channels for UPF1, UPF2 and UPF3B. All three proteins were readily detected in dendrites. MAP2 was used to visualize dendrites. **b-c**, *Arc* and *AMPK* mRNAs are subjected to NMD in dendrites. Arc is known to influence surface levels of GluR1 by regulating GluR1 internalization [21, 61]. Intriguingly, AMPK, another protein whose mRNA contains intron retention in the 3’*UTR*, has also been reported to regulate protein levels by suppressing protein synthesis [111]. Because both internalization and local synthesis of GluR1 are altered in UPF2-deficient dendrites, we asked whether *Arc* and *AMPK* mRNAs are subjected to NMD in the synaptic channels of tripartite microfluidic devices. We first examined mRNA levels upon knockdown of *UPF2*. We cultured E16 neurons in tripartite chambers and infected postsynaptic neurons with *control*- or *UPF2*-shRNA lentivirus at DIV7. We harvested material from synaptic channels for qRT-PCR at DIV21. Compared to the synaptic material harvested from cultures infected with *control*-shRNA virus, both *Arc* and *AMPK* mRNAs were elevated in the synaptic regions of UPF2-deficient postsynaptic neurons (n=3 biological replicates) (**b**). NMD is translation dependent; specifically, NMD recognizes its targets only upon their translation. Therefore, inhibition of translation causes an increase in the levels of mRNAs normally subjected to NMD, thus identifying accumulated mRNAs as NMD targets. To determine whether *Arc* and *AMPK* are accumulated in dendrites upon inhibition of translation, we selectively treated synaptic channels with the translation inhibitor CHX (10 μM) at DIV21 for 3 hr. This resulted in the significant accumulation of both *Arc* and *AMPK* mRNAs (n=3 biological replicates) Together, these data suggest that NMD degrades both *Arc* and *AMPK* locally within dendrites. **d**, NMD locally regulates surface expression of GluR1 in dendrites. In the previous experiments, *UPF2* was knocked-down in postsynaptic neurons, which did not directly test a local role of NMD in regulating GluR1 signaling specifically in dendrites. To disrupt NMD locally within dendrites, we interfered with the local synthesis of UPF3B by applying a siRNA cocktail against the *UPF3B* mRNA selectively to the synaptic channels of tripartite chambers. UPF3B is specific to the NMD pathway, and its local synthesis in dendrites is the only source for this protein in these compartments (see Figure S9a-b) allowing the study of local NMD. We cultured E16 neurons in cell-body channels and, starting at DIV14, we repeatedly treated synaptic channels with non-overlapping *UPF3B*-siRNAs (10 nM) for 7 days, which led to complete loss of UPF3B exclusively in these channels (see Figure S9d). Exclusive loss of UPF3B in synaptic channels resulted in a greater than 40% decrease in the surface levels of GluR1 compared to control cultures (n=10; 10 neurons and 10 dendrites per group). This suggests that the NMD-mediated regulation of GluR1 also occurs locally in dendrites. **e**, Locally synthesized AMPK negatively regulates GluR1 levels in dendrites. We next asked whether locally synthesized AMPK, whose mRNA is locally targeted by NMD, modulates dendritic GluR1 levels. To do this, we inhibited AMPK synthesis in the synaptic channels of tripartite microfluidic devices and examined total GluR1 levels. Application of 10 nM of two non-overlapping *AMPK*-siRNAs to synaptic channels for 7 days, starting at DIV14, led to selective depletion of AMPK in dendrites in synaptic channels but not in dendrites in microgrooves (Figure S9e). Local ablation of AMPK increased the total levels of GluR1 by ~80% in synaptic channels (n=3 biological replicates per *control*-siRNA [6 neurons, 9 dendrites] and per *AMPK*-siRNA [8 neurons, 9 dendrites] (**b**). This suggests that elevated levels of dendritic AMPK contribute to the reduction in total GluR1 levels in UPF2-deficient dendrites (see Figure S3a). Data are represented as mean ± SEM; ****p < 0.0001. Scale bar: 30 μm.

To determine whether NMD-mediated mRNA degradation occurs in dendrites, we examined the *AMPK* and *Arc* mRNAs upon selective inhibition of translation in these compartments. NMD can only occur when ribosomes recognize NMD-inducing features in mRNAs during the first round of translation [84, 85]. Inhibition of translation hinders NMD and leads to the accumulation of NMD-target mRNAs [7]. Therefore, a widely used approach to determine whether an mRNA is subjected to NMD is to determine if it accumulates after the treatment of cells with cycloheximide (CHX), a translation inhibitor. To confirm whether these targets undergo translation-dependent degradation in synaptic regions, we cultured E16 neurons in tripartite chambers and, at DIV21, inhibited translation selectively in synaptic channels. qRT-PCR on material harvested from the synaptic channels showed that both *AMPK* and *Arc* mRNAs accumulated in these compartments upon inhibition of translation (Figure 4c).

We next asked whether NMD contributes to the regulation of GluR1 surface levels locally in dendrites. To address this, we targeted NMD selectively in synaptic channels by inhibiting the local synthesis of UPF3B, a protein specific to the NMD pathway [86, 87]. From the NMD machinery, only the *UPF3B* mRNA is localized to dendrites [22] (Figure S9a). We confirmed that the *UPF3B* mRNA is locally translated in dendrites, and that its local translation is the only source for UPF3B protein in synaptic channels (Figure S9b). To disrupt local NMD in dendrites, we targeted the *UPF3B* mRNA by siRNA transfection of synaptic channels. The siRNA-mediated depletion of targets has been successfully used in microfluidic devices previously, allowing exclusive knockdown in neurites without influencing expression in cell bodies or even microgrooves [88–93]. We cultured E16 neurons in cell-body channels and, starting at DIV14, repeatedly treated synaptic channels with siRNAs against *UPF3B* mRNA for seven days. This resulted in complete loss of UPF3B protein in synaptic channels (Figure S9d). To determine whether the expression of GluR1 is affected upon a strictly local disruption of NMD in synaptic channels, we measured the frequency of surface GluR1 upon knockdown of local *UPF3B* in these channels. Similar to the knockdown of *UPF2* in postsynaptic cells, the surface frequency of GluR1 was reduced upon selective loss of UPF3B protein in synaptic channels (Figure 4d).

Taken together, these experiments establish the local requirement for NMD in synaptic regions and suggest that the local regulation of GluR1 by this pathway might involve *AMPK* degradation in addition to *Arc*.

### AMPK negatively regulates GluR1 expression in dendrites

We next examined whether AMPK locally influences GluR1 expression in synaptic channels. To do this, we cultured E16 neurons in cell-body channels and, starting at DIV14, repeatedly treated synaptic channels with siRNAs against the *AMPK* mRNA for seven days. This resulted in an almost complete loss of AMPK protein in synaptic channels (Figure S9e). We then evaluated the total levels of GluR1 in synaptic channels upon loss of AMPK. Selective inhibition of AMPK in synaptic channels significantly increased the total levels of GluR1 (Figure 4e).

Because the synaptic channels of tripartite chambers contain both dendrites and axons, it is possible that GluR1 phenotypes observed after knockdown of local *UPF3B* or local *AMPK* may arise also from axonal changes and therefore not selectively from dendrites. To clarify this, we examined the proteins of the NMD machinery and targets of NMD in mature axons. Neither the NMD-specific components of the NMD machinery, nor *AMPK* or *Arc* mRNAs, were present in isolated hippocampal axons at DIV21 (Figure S10). In fact, the axonal transcriptome significantly shrinks as axons mature [94] and does not include any known NMD targets [95, 96]. These data indicate that alterations in GluR1 expression upon selective disruption of either UPF3B or AMPK proteins in synaptic channels solely originate from a local effect within dendrites.

### Normalization of *AMPK* and *Arc* mRNA levels rescues GluR1 surface expression in UPF2-deficient dendrites

Lastly, we functionally tested a mechanistic role of locally synthesized AMPK and Arc in the NMD-mediated regulation of surface GluR1 expression. To do this, we systematically modulated the elevated levels of *Arc* and *AMPK* mRNAs in UPF2-disrupted dendrites by siRNA transfection of synaptic channels. As siRNA treatment affects mRNA expression in a concentration-dependent manner [97–100], we first determined the amount of siRNA cocktail (see Methods) required to correct levels of *Arc* and *AMPK* mRNAs in UPF2-deficient dendrites. First, we applied the *UPF2*-shRNA virus to postsynaptic cell channels at DIV7. Starting at DIV14, we treated the synaptic channels of UPF2-deficient dendrites with non-overlapping siRNA cocktails at different concentrations for each target mRNA (Figure S11). After determining the amount of siRNA needed to normalize the exaggerated levels of *Arc* and *AMPK* mRNAs, we modulated the levels of these targets in synaptic channels containing UPF2-deficient dendrites (Figure 5a). After a week of siRNA treatment, we evaluated the surface expression of GluR1. Compared to synaptic channels with intact NMD (i.e., those treated with *control*-shRNA lentivirus), cultures infected with *UPF2*-shRNA lentivirus had significantly lower GluR1 density as expected. Restoring expression levels of *Arc* or *AMPK* mRNAs in synaptic channels led to a significant but incomplete recovery of surface GluR1 in these compartments for each target (Figure 5b-c). We next considered whether the surface GluR1 phenotype in UPF2-deficient dendrites might be a combinatorial effect of both elevated Arc and AMPK, with altered GluR1 surface levels arising from an amalgamation of both increased internalization and reduced protein synthesis. Normalization of both *Arc* and *AMPK* mRNAs together led to a complete recovery of GluR1 density in UPF2-deficient dendrites. This data indicates that local NMD contributes to the regulation of GluR1 surface expression through physiological degradation of both *Arc* and *AMPK* mRNAs in dendrites.

**Figure 5.**
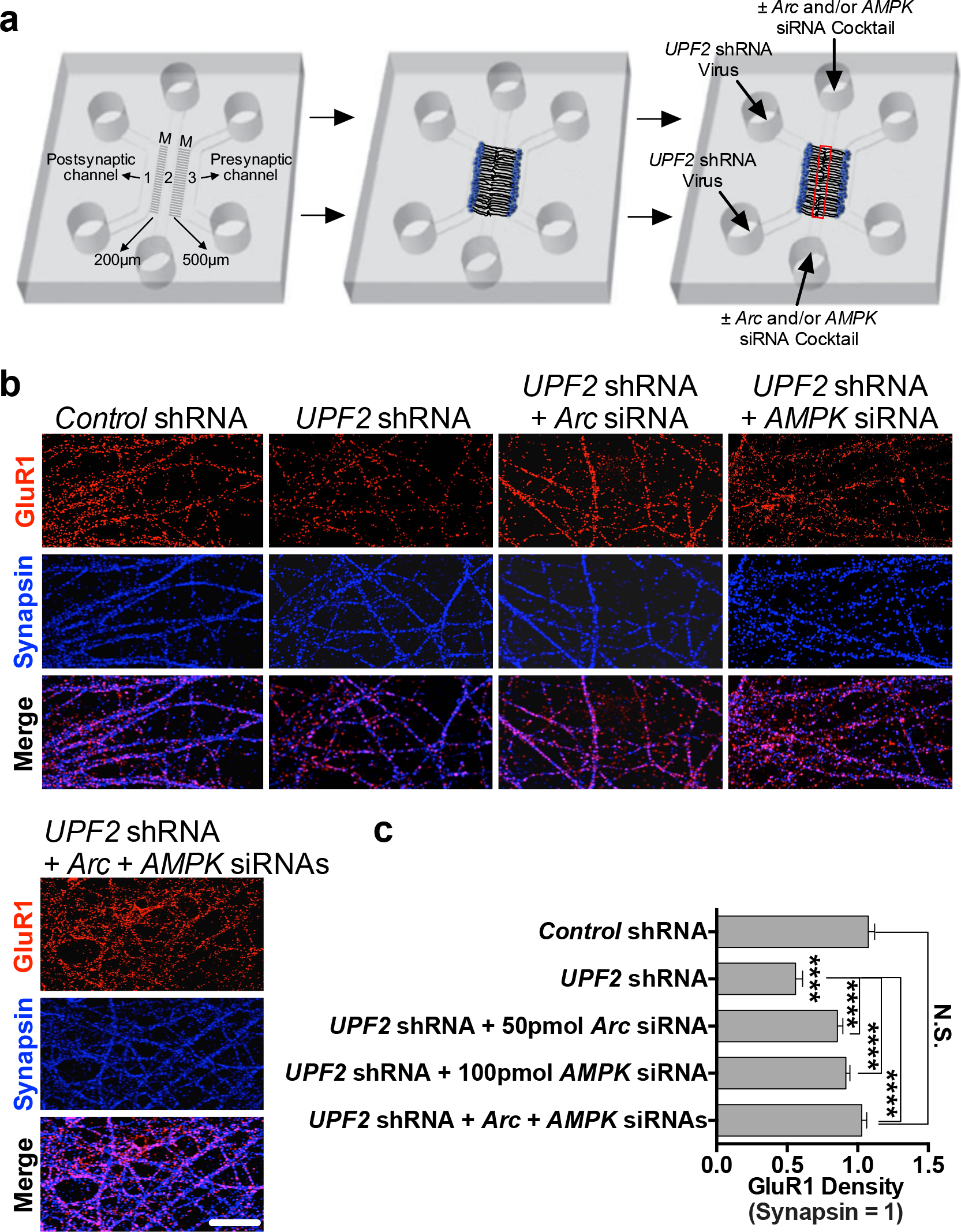
NMD regulates surface GluR1 levels via degradation of *Arc* and *AMPK* in dendrites. **a-c**, We next sought to functionally test the role Arc and AMPK in NMD-mediated GluR1 regulation. To do this, we targeted NMD only in postsynaptic neurons by applying *UPF2*-shRNA virus in the postsynaptic cell channels of tripartite chambers at DIV7, and then systematically normalized the elevated levels of NMD targets in dendrites by siRNA transfection of synaptic channels at DIV14 for a week (**a**). First, we determined the amount of siRNA required to correct the levels of each target in UPF2-deficient dendrites by treating the synaptic channels of tripartite chambers with non-overlapping siRNA cocktails at different concentrations (10 pm - 1 nm) for each target mRNA (see Figure S11). Immunostainings and quantifications were performed within synaptic channels (red box). Compared to cultures with intact NMD in postsynaptic cells (i.e., those treated with *control*-shRNA lentivirus), cultures infected with *UPF2*-shRNA lentivirus had significantly lower surface GluR1 as expected. Restoring expression levels of *Arc* (50 pmol of siRNA) or *AMPK* (100 pmol of siRNA) in UPF2-deficient dendrites led to a significant but incomplete recovery of GluR1 density for each target (n=3 biological replicates per *control*-siRNA [10 neurons, 10 dendrites], *Arc*-siRNA [10 neurons, 10 dendrites] and per *AMPK*-siRNA [10 neurons, 10 dendrites]). We next considered whether reduced GluR1 in UPF2-deficient dendrites might be a combinatorial effect of both Arc and AMPK, due to an amalgamation of both increased internalization and decreased translation from each respective molecule. Co-normalization of both *Arc* and *AMPK* levels led to a complete recovery of GluR1 surface density in NMD-deficient dendrites, resulting in an expression profile indistinguishable from control cultures. These data indicate that increased *Arc* and *AMPK* upon loss of NMD together account for the low surface expression of GluR1 in dendrites. Data are represented as mean ± SEM; ****p < 0.0001, N.S. represents “Not Significant”. Scale bar: 75 μm.

## Discussion

Our study identifies neuronal NMD as a mechanism that contributes to the regulation of synaptic plasticity and cognitive function. Specifically, we report that NMD influences synaptic plasticity as well as learning and memory. In addition to the enrichment of NMD-machinery components in dendrites, which suggests that NMD functions locally in these compartments, our data indicates that local NMD participates in GluR1 regulation by limiting the amount of Arc and AMPK proteins in dendrites.

### Role of NMD in regulating synaptic plasticity and cognitive function

Our data indicate that proper synaptic plasticity and cognitive function requires NMD. Our data show that *UPF2*-deficient hippocampal slices show decreased L-LTP. In addition, spine density in the hippocampus of conditional *UPF2* mutant mice is reduced relative to control mice. Mice also show attenuated learning and memory upon conditional ablation of *UPF2* in excitatory neurons. Memory deficits were readily detectable in *UPF2*-deficient mice on three independent hippocampus-dependent behavioral assays. In addition to attenuated learning and memory, loss of *UPF2* in excitatory neurons resulted in perseverative behavior in mice, including increased marble burying and self-grooming behavior. Together, these data implicate a role for NMD in both synaptic plasticity and behavioral endophenotypes common to a spectrum of neurodevelopmental diseases including autism and schizophrenia.

Our data also provide comprehensive insight into the neuron-specific and spatiotemporal requirement for NMD in plasticity and behavior. Recently, a *UPF3B* null mutant mouse line was shown to have fear-conditioning defects but, unlike our observations in *UPF2* conditional knockout mice, no spatial learning deficit [33]. There may be several reasons for this. For instance, NMD is ubiquitous and required for neuronal differentiation [34, 101]. Consistent with this, global interruption of NMD by ablation of *UPF3B* during brain development alters neuronal maturation and is thereby likely to contribute to the behavioral phenotypes observed in the *UPF3B* null mouse line. In addition, while UPF2 and UPF3 proteins are known to be specific to the NMD machinery, it has been suggested that NMD might have different branches in which NMD exerts its function independent of UPF2 or UPF3B [87, 102, 103]. Therefore, studies of conditional *UPF2* deletion provide a platform to dissect not only the temporal and cell-type specific functions of the NMD machinery, but also branch-specific functions of NMD in brain development and behavior.

### NMD-mediated GluR1 regulation and its contribution to behavior

Our data also demonstrate that NMD positively regulates GluR1 signaling in hippocampus. The dynamic insertion and removal of GluR1 receptors to the synaptic surface directly modulates GluR1 signaling [49]. We report that loss of UPF2 in adult hippocampal neurons caused a concomitant decrease in total and surface levels of this receptor. *GluR1* mRNA expression was not significantly changed in UPF2-deficient neurons, indicating that *GluR1* mRNA is not degraded upon disruption of UPF2. This suggests that synthesis or degradation of the GluR1 receptor must be regulated via proteins affected by NMD-mediated degradation. Our CLICK-chemistry experiments suggest that NMD positively regulates GluR1 protein synthesis. GluR1 synthesis is repressed in the absence of UPF2 protein, causing a drop in the total levels of GluR1 protein. In addition, loss of UPF2 results in a higher GluR1 internalization rate. These mechanisms together account for the surface GluR1 phenotype caused by the disruption of NMD. Therefore, NMD-dependent GluR1 regulation involves restraining synthesis and internalization of the GluR1 receptor.

Modulation of AMPA-receptor integrity plays a crucial role in synaptic strength. Strengthening of synapses by LTP or weakening of synaptic strength by LTD can be represented by the synaptic insertion or removal of AMPA receptors, respectively [56]. Consistent with this, alterations in GluR1 levels are linked to impairments in LTP as well as abnormalities in memory [52–54]. Our data show that hippocampal L-LTP, which is thought to underlie memory consolidation, is decreased in conditional *UPF2* mutant mice. Activation of glutamate receptors facilitates LTP either by through the enlargement of pre-existing spines or promoting the formation of new spines. Therefore, a reduction in spine density is consistent with a decrease in L-LTP in conditional *UPF2* mutants. This phenotype is accompanied by behavioral defects, including a lack of short-term spatial memory in the Y-maze, defective associative fear learning and memory, and attenuated spatial learning and memory on the Morris water maze. Consistent with these data, reductions in GluR1 expression were shown to cause spatial memory dissociations in mice [58]. Similarly, contextual conditioning requires increased levels and surface localization of GluR1 [52, 104, 105]. We show that UPF2-deficient neurons display reduced GluR1 synthesis, which results in an overall decrease in GluR1 levels. NMD-dependent control of GluR1 signaling is therefore likely to contribute to the plasticity and behavioral phenotypes conditional *UPF2* mutant mice exhibit. However, we cannot exclude the possibility that NMD regulates other receptors or intracellular signaling pathways that might also contribute to LTP, learning and memory. Although the ionotropic glutamate receptors, N-methyl-D-aspartate receptors (NMDARs), are more static than AMPA receptors in mature neurons, subunits of NMDARs are also locally translated in dendrites [106], suggesting a potential function for NMD in regulation of NMDAR dynamics.

### Role of local AMPK in GluR1 regulation and synaptic plasticity

Our data also show that the NMD-target AMPK, whose levels are elevated in UPF2-deficient dendrites, negatively regulates GluR1 levels. The AMP kinase consists of one α, one β, and one γ subunit. We found that the mRNA of the γ subunit, which is required for AMPK activation, is a substrate of NMD (Figure 4b-c). Our data shows that local disruption of AMPK within dendrites led to an increase in GluR1 levels, suggesting that local AMPK functions to negatively regulate GluR1 levels. Activated AMPK is known to interrupt translation by either inhibiting the mammalian target of rapamycin (mTOR) kinase or the translation elongation factor 2 [78–83]. Our data indicates that the local synthesis of GluR1 protein in dendrites is reduced upon disruption of UPF2, which coincides with an elevation of AMPK levels in UPF2-deficient dendrites. We also show that modulation of AMPK levels within these dendrites is required to restore GluR1 surface levels (Figure 5b-c). These suggest that while AMPK represses GluR1 synthesis, NMD limits the amount of AMPK thereby serving as a balance in regulating total levels of the GluR1 receptor. Whether AMPK regulates the synthesis of other synaptic proteins and how this specificity is achieved is yet to be determined. Intriguingly, inhibition of AMPK was shown to alleviate impairments in hippocampal synaptic plasticity in a mouse model of Alzheimer’s disease [107]. Inhibition of AMPK corrected the Amyloid β-induced inhibition of LTP suggesting that AMPK has negative effects on LTP induction. This finding is consistent with our data that L-LTP was decreased in UPF2-deficient hippocampal slices.

### Activity-induced local NMD as a regulator of the dendritic proteome

We found that NMD can occur locally in dendrites. *Arc* mRNA was shown to be locally degraded in hippocampus [108]. However, the involvement of NMD in this, as well as more broadly hippocampal function, has not been demonstrated. Our data show that the NMD machinery is localized to dendrites, and the levels of the NMD targets *Arc* and *AMPK* are elevated upon selective inhibition of translation in these compartments (Figure 4c). The mRNA of UPF3B is trafficked to [22] and locally translated in dendrites (Figure S9). Local inhibition of dendritic UPF3B synthesis results in dysregulated surface localization of GluR1, indicating that NMD locally influences GluR1 levels within not just neuronal cell bodies but also their dendrites. This is consistent with the dynamic modulation of GluR1 surface levels at synapses.

Our data demonstrate that local translation is regulated by mechanisms that control mRNA degradation in dendrites. Synaptic plasticity relies on local protein synthesis at synaptic sites [21, 38]. Locally synthesized proteins are thought to either strengthen a pre-existing synaptic connection, by inducing structural remodeling, or to promote the formation of new synaptic connections. NMD is likely to have more targets than *Arc* and *AMPK* in dendrites. In addition to its canonical targets, NMD can also degrade mRNAs that do not carry any NMD-inducing features [63]. The mechanisms by which NMD recognizes its atypical targets remain unclear. This raises the possibility that NMD might be regulating various physiological mRNAs in dendrites that cannot be predicted based on sequence information. Similarly, RNA sequencing cannot distinguish between 1) true NMD targets, 2) secondary effects mediated by NMD-targets or 3) compensatory mechanisms that may occur upon disruption of NMD. Therefore, we cannot exclude the possibility that atypical targets of NMD may also contribute to the NMD-mediated GluR1 regulation at synapses as well as other synaptic events.

Our data also suggest that NMD is positively regulated by synaptic activity. mRNA degradation is regulated during development of the nervous system. microRNAs negatively regulate NMD activity in neural progenitors and impact the expression of a broad range of NMD targets that are important for appropriate differentiation [109]. *UPF3B* mRNA is locally translated in dendrites, where local translation of specific mRNAs is induced by synaptic stimuli [110]. This suggests that NMD activity might also be regulated in the context of synaptic function. Our data show that dendritic UPF3B protein is upregulated when synaptic channels are selectively treated with the group I mGluR agonist DHPG, a well-known synaptic stimulus (Figure S12). Activation of mGluRs induces the local translation of *Arc* [21], directing *Arc* mRNA to NMD in dendrites. NMD induction is thus likely to modulate the impact of synaptic activity on GluR1 internalization by limiting the amount of Arc protein. It is not known whether synaptic activity induces local synthesis of *AMPK* to suppress GluR1 synthesis in the long term. If that were the case, NMD would also limit the amount of AMPK to positively regulate total GluR1 levels upon synaptic activity.

STAU1-mediated mRNA decay (SMD) has been suggested as a competitive pathway to NMD in mammalian cells [111]. When bound to the *3’UTR*, STAU1 can elicit translation-dependent mRNA degradation. STAU1 is known to associate with dendritic mRNA transport granules. Like NMD, numerous neuronal mRNAs may be natural SMD targets. Thus, mRNA decay in dendrites might be a broader phenomenon that serves as a natural brake to influence local translation, which is in turn required for synaptic plasticity.

## Conclusions

Synaptic plasticity requires the selective translation of specific mRNAs targeted to dendrites. If this process is disrupted neurodevelopmental disorders may arise. Our experiments provide fundamental support for the idea that NMD-mediated mRNA decay modulates dendritic mRNA availability and therefore synaptic plasticity, learning, and memory. Through a number of locally-confined experiments in microfluidic devices and behavioral/LTP assays in neuron-restricted *UPF2* conditional knockout mice, we define a specific role for the NMD pathway in dendrites and synaptic function (see schematic model in Figure S13). Therefore, our paper provides novel insights into basic mechanisms that regulate synaptic function that are also likely to play a role in NMD-implicated diseases such as autism and schizophrenia.

## Acknowledgements

We thank J. Lykke-Andersen for generously providing the anti-UPF1 and anti-UPF2 antibodies, M.E. Ross for helpful comments and suggestions and M. Toth for helpful suggestions regarding behavioral assays. We also thank Nicole Volk, the research assistant of Colak lab, for technical help. This work was supported by a NHMRC CJ Martin Biomedical Fellowship awarded to M.N., KoreaNRF-2015R1A2A1A09005662 to N.L.J., NIH grants NS034007 and NS047384 to E.K, and NIH R01 MH114888 and Leon Levy Foundation Grants to D.C.

## Author contributions

M.N. performed and/or analyzed most of the experiments including behavioral assays, and manuscript preparation/writing. M.A. prepared all neuronal cultures. F.L. performed electrophysiology experiments. N.V. provided technical assistance. M.T. provided infrastructure support. N.L.J. designed and manufactured microfluidic devices. E.K. supervised electrophysiology experiments. D.C. initiated and conceived the project; designed, analyzed, and supervised experiments; and wrote the manuscript. All authors contributed to the experimental design and interpretation and commented on the manuscript.

## Methods

### Mice and constructs

Hippocampal neuronal cultures were prepared from C57BL/6J mouse embryos (Charles River Laboratories). *UPF2*-shRNA 1 virus is on a piLenti-shRNA-GFP backbone carrying one shRNA against the *UPF2* mRNA (AGGCGTATTCTGCACTCTAAAGGCGAGCT). *UPF2*-shRNA 2 virus is on a piLenti-shRNA-GFP backbone carrying four shRNAs against the *UPF2* mRNA (TGAAAGACTATGTTATTTGTTGTATGATA). In experiments with *UPF2* deletion in excitatory neurons, a *UPF2* conditional knockout mouse line, which carries flanked loxP sites in the second exon of this gene, [1] was crossed with α*CaMKII*:*CreER^T2^* line [2], and *UPF2*^wt/wt^;α*CaMKII*:*CreER^T2^* (CTRL) or *UPF2*^fl/fl^;α*CaMKII*:*CreER^T2^* (CKO) adults were used. To induce Cre, mice were treated with 200 mg tamoxifen/kg body weight every other day for five days. To outline dendrites and spines, an AAV-*CaMKIIa*-*eGFP* virus (Addgene; 50469-AAV5) was injected into the adult hippocampus of CTRL and CKO mice.

### Hippocampal neuronal cultures

For culturing hippocampal neurons, we followed a previously described protocol with some modifications [3]. We typically isolated hippocampal neurons from embryos at embryonic day 16 (E16) and plated 1×10^5^ neurons per coverslip (diameter: 12 mm) pre-coated with poly D-lysine (PDL) in Neurobasal medium containing 2% B27, 1 mM sodium pyruvate, 2 mM Glutamax, 30% D-glucose and penicillin (100 units/ml) /streptomycin (100 μg/ml).

### Compartmentalized culture in microfluidic devices

Tri-partite microfluidic device was designed and manufactured by the Jeon lab. Microfluidic chambers were prepared in house as described previously described [4]. To isolate synaptic regions or to selectively treat the synaptic regions as well as post- or presynaptic cells with either pharmacological agents or sh/siRNAs, hippocampal neurons were cultured in tripartite microfluidic devices. The polymethyldimethylsiloxane (PDMS) devices were fabricated as previously described [4]. These devices have different channels separated by a physical barrier with embedded microgrooves (10 μm wide, 3 μm high). Unlike traditional microfluidic chambers with two channels [5], the tripartite PDMS chamber contains three channels (Figure 3). To access a high number of dendritic synapses, a ‘synaptic’ channel was placed 200 μm from the edge of one channel and 500 μm away from the other. Since the average length of dendrites extending into the microgrooves is less than 300 μm at 21 days in culture (DIV21), the closer “postsynaptic cell channel” allows a large number of dendrites to project from this channel to the new “synaptic” channel. On the other hand, the “presynaptic cell channel” that is located 500 μm away from the new channel ensures that only axons from this channel can extend into the synaptic channel. Fluidic isolation is established across the microgrooves, making it possible to isolate synaptic regions or to selectively treat the synaptic regions as well as post- or presynaptic cells. Reagents applied to synaptic channels were as follows: Cycloheximide (10 μM, Calbiochem), MG132 (10 μM, Sigma), and Leupeptin (10 μM, Sigma). siRNAs were applied to synaptic channels at different concentrations using 10% NeuroPORTER (Sigma) following manufacturer’s instructions.

For microfluidic cultures, we reversibly affixed the tripartite chambers to PDL-coated coverslips (Assistant; 50×24mm; two per coverslip) and then place them in 10 cm dishes. For culturing hippocampal neurons, we followed a previously described protocol with some modifications [3]. We typically isolated hippocampal neurons from embryos at embryonic day 16 and plate 1×10^5^ neurons per coverslip (diameter: 12mm) pre-coated with poly D-lysine (PDL) in Neurobasal medium containing 2% B27, 1 mM sodium pyruvate, 2 mM Glutamax, 30% D-glucose and penicillin (100 units/mL) /streptomycin (100 μg/mL). We plated 8×10^4^ neurons in cell-body channels and maintain these cultures in neuronal medium for up to 21 days. For immunostaining, we fixed neurons inside microfluidic chambers and continued with the staining procedure upon careful removal of chambers.

### Immunohistochemistry

Hippocampal neuronal cultures were fixed in 4% paraformaldehyde (PFA)/PBS for 15 min, which was followed by 3 × 10 min washes in PBS. Standard immunohistochemistry was performed as previously described [4]. Briefly, samples were incubated overnight in PBS containing 1% Triton X-100, 10% normal goat serum and the relevant dilution of primary antibodies. The following morning, samples were washed with PBS (3 × 10 min) to remove any residual primary antibody. Secondary antibodies were incubated in PBS containing 1% Triton X-100 and 10% normal goat serum for 2 hr. Samples were subsequently washed with PBS (3 × 20 min) and mounted (Vectashield, Vector Laboratories). Primary commercial antibodies included AMPK (rabbit; Thermofisher, PA5-13797, PRKAG3 subunit), Arc (rabbit; Synaptic Systems, 156002), MAP2 (mouse; Abcam, ab11276), synapsin (rabbit; Abcam, ab64581) and UPF3B (rabbit; Sigma, HPA001882). Two commercial GluR1 antibodies were used: mouse GluR1 antibody (Abcam, ab174785) for stainings with synapsin and rabbit GluR1 antibody (Calbiochem, pc246) for the internalization assay. Anti-UPF1 and anti-UPF2 antibodies (both rabbit) were a gift from Jens Lykke-Andersen. Secondary antibodies were Alexa 488, 546, and 647 conjugated (Molecular Probes, 1:2000).

### qRT-PCR

RNA was isolated from the synaptic channels of microfluidic devices by perfusion with TRIzol reagent (Invitrogen), with RNA being subsequently extracted from lysates following the manufacturer’s instructions. Purified RNA was inbcubated with DNase I (RNase-Free DNase Set, Qiagen) over a mini kit column (RNAeasy, Qiagen) to ensure DNA digestion. SuperScript III First-Strand Synthesis SuperMix (Invitrogen) was used to amplify cDNAs in a 20 μl reaction using 1 μg of RNA. Quantitative Real-Time PCR (qRT-PCR) reactions were performed as previously described [4] using the iQ SYBR Green Supermix (Bio-Rad) and an Eppendorf Mastercycler EP Realplex Thermocycler. For each reaction 10 ng cDNA was used. Relative expression levels were either normalized to *GAPDH* or *Beta-Actin*.

Primers used for qRT-PCR:

*Arc* Primers:

(FW) 5’CTGAGATGCTGGAGCACGTA3’

(REV) 5’GCCTTGATGGACTTCTTCCA3’

*AMPK* primers:

(FW) 5’ CAGGTCTACATGCACTTCATGC 3’

(REV) 5’ AAAGCTCTGCTTCTTGCTGTCC 3’

*GluR1* primers:

(FW) 5’AGAGAAGAGGAGGAGAGCAG 3’

(REV) 5’ CTATCTGGATATTGTTGGGGA 3’

*GAPDH* primers:

(FW) 5’ ACATGGTCTACATGTTCC 3’

(REV) 5’ CAGATCCACAACGGAATAC 3’

*Beta-Actin* primers:

(FW) 5’ AGTGTGACGTTGACATCCGT 3’

(REV) 5’ TGCTAGGAGCCAGAGCAGTA 3’

Dicer-substrate siRNAs were acquired from Integrated DNA Technologies, and were predesigned using their proprietary algorithm which integrates design principles from 21mer siRNA theory with updated 27mer criteria. This design process ensures siRNA target sites do not match known single nucleotide polymorphisms or exons that are the product of alternative splicing, with all sequences having been pre-screened using Smith-Waterman analysis to reduce off-target binding.

*Arc* siRNA Cocktail

Duplex 1:

5’ CUGAUGGCUAUGACUAUACCGUUAG 3’

3’ UCGACUACCGAUACUGAUAUGGCAAUC 5’

Duplex 2:

5’ CUACAUGGACUGAACAUCAAGAAGC 3’

3’ ACGAUGUACCUGACUUGUAGUUCUUCG5’

*AMPK* siRNA Cocktail

Duplex 1:

5’ GACCUUGUUUCAAAUUGAAACAATA 3’

3’ UUCUGGAACAAAGUUUAACUUUGUUAU 5’

Duplex 2:

5’ CGAGAAAUUCAAAAUCUUAAACUCT 3’

3’ UUGCUCUUUAAGUUUUAGAAUUUGAGA 5’

*UP3B* siRNA Cocktail

Duplex 1:

5’ AAAUUGAAGCCAAAAAUCGAGAATT 3’

3’ CCUUUAACUUCGGUUUUUAGCUCUUAA 5’

Duplex 2:

5’ GGGUCAAGAAUAUCAUGCUAUAGTA 3’

3’ UUCCCAGUUCUUAUAGUACGAUAUCAU 5’

### Fluorescent *in-situ* hybridization (FISH) for *UPF3B* mRNA

For the detection of *UP3B* mRNA, a mix of five non-overlapping antisense probes were used. Each oligonucleotide was 49 base pairs in length. FISH was visualized using the Tyramide In Situ System (Perkin Elmer) as previously described [4]. Briefly, samples were pretreated with hybridization buffer (50% formaldehyde, 5 × SSC, 0.1% tween, 100 ug/ml tRNA [Roche], 50 ug/ml heparin [Sigma]) for 2 hr at 65°C. Riboprobes were (100 ng/μl) were next heated to 97°C for 7 min and left to cool on ice. Hybridization was performed overnight at 65°C, and was followed by a series of washes at room temperature in hybridization buffer (no tRNA or heparin, 2 × 30 min), hybridization buffer/PBT (0.1% tween in PBS, 2 × 30 min), and lastly just PBT (4 × 30 min). Samples were next blocked for 1 hr (Roche Blocking Solution) and incubated with anti-DIG-POD (Roche) 1:800 in PBT overnight at 4°C. The following morning, samples were washed in PBT (3 × 20 min). Detection was visualized with tyramide-488 1:50 in supplied amplification buffer from the TSA-Plus *in situ* kit (Perkin Elmer).

Oligonucleotides used for antisense *UPF3B* probes

1-GGAAGAAAAAGATCATAGGCCTAAGGAGAAACGAGTGACCCTGTTTACG
2-GTCAAGAATATCATGCTATAGTAGAATTTGCACCATTTCAAAAAGCTGC
3-AAGAAAAGAGAGAAGAAAGGAGGAGACGGGAAATAGAGAGGAAAAGGCA
4-AAGAAAGAGCCAGTGGGCACAGTTATACTCTGCCCAGGCGTTCTGATGT
5-GGCACTTCGAGATAAAGGAAAGAAGAGTGAGAATACAGAATCAATATGC

### GluR1 internalization assay

GluR1 internalization experiments were carried-out and quantified as previously described [6, 7]. Briefly, we incubated live DIV21 hippocampal cultures infected with control or *UPF2* shRNAs at 37°C for 5 min with GluR1 antibody (Calbiochem, PC246) to allow labeling of surface GluR1. After washing, we incubated neurons at 37°C for 10 min to allow for basal internalization. We then fixed neurons with 4% PFA in PBS and blocked in a detergent-free blocking solution for 1 hr. Following blocking, we incubated neurons with AlexaFluor546 secondary antibody for 1 hr to label the surface population. To label the pre-labeled internalized fraction, we post-fixed neurons with 100% methanol at −20°C for 1 min and stained with Alexa-647 secondary antibody. We calculated the percent internalization by dividing the integrated far-red intensity (internalized GluR1) by the total (red + far red) intensity between control and NMD-deficient neurons.

### Click chemistry and SDS-page

To measure the dendritic translation of *GluR1*, we used a click chemistry based approach and labeled locally synthesized GluR1 at synaptic regions. At DIV21, we exchanged the growth media in the synaptic channels to methionine-free medium containing the methionine-analog azidohomoalanine (AHA; 25 μM) for 6 hr. AHA is incorporated into all newly synthesized proteins. After harvesting the material from the synaptic channels in RIPA, we incubated the lysate with DBCO-biotin [8] (30 μM; Click Chemistry Tools LLC) for 1 hr at ambient temperature. DBCO reacts with AHA [9], functionalizing newly synthesized proteins with biotin. We purified biotin-containing proteins with Dynabeads (150 μl per reaction; Dynabeads M-280 Streptavidin-Thermo Fisher Scientific). Dynabeads were separated over a magnetic rack and subsequently washed 2 times in PBS containing 0.1% Bovine Serum Albumin (BSA). Proteins were dissociated and denatured from beads by boiling Dynabeads for 5 min in 0.1% SDS. Lysates were subsequently analyzed for protein concentration via BCA Protein Assay (Thermofisher). Proteins were separated via SDS-PAGE, as previously described [10]. Briefly, protein aliquots were mixed with an equal volume of loading buffer, denatured at 95°C and loaded onto 8% gels before being subject to SDS-PAGE. Proteins were subsequently transferred to nitrocellulose membrane, incubated in 5% BSA diluted in TBST for 1hr and then incubated with primary anti-C-terminus GluR1 antibody overnight (Abcam, ab31232). Membranes were washed in TBST (3 × 5 min), incubated for 1 hr in IRDye 800CW goat anti-rabbit secondary (Licor) diluted in 5% low-fat milk diluted in TBST, and bands resolved on a Licor Odyssey CLx imager.

### Electrophysiology

The electrophysiology experiments were performed as previously described (114). All experiments were performed on transvers hippocampal slices (400 μm) from 3-month old (LTP studies) or 6-week-old (LTD studies) *UPF2*^fl/fl^;α*CaMKII*:*CreER^T2^* mice and their control littermates (*UPF2*^wt/wt^;α*CaMKII*:*CreER^T2^*) following induction of Cre expression by tamoxifen (Sigma) at 2-months (for LTP) and 1-month (for LTD) of age. Given that Tamoxifen is fat soluble, we prepared our tamoxifen solution by dissolving the relevant quantity of drug into sunflower oil with 1:10 ETOH added. Tamoxifen solutions were incubated at 37°C with rocking for 6-8 hr. Tamoxifen was administered via oral gavage at 200 mg tamoxifen/kg body weight every other day for five total doses. This tamoxifen regimen led to successful ablation of UPF2 protein in the adult brain (Figure S1a). Slice preparation and aCSF composition were performed as described previously [11, 12]. Briefly, slices were isolated and transferred to recording chambers (preheated to 32°C), where they were superfused with oxygenated aCSF. We recorded field excitatory postsynaptic potentials (fEPSPs) from the stratum radiatum of CA1 using microelectrodes filled with artificial cerebrospinal fluid (ACSF) (resistance 1-4 MΩ). A bipolar Teflon-coated platinum electrode was placed in the stratum radiatum of CA3 to activate Schaffer collateral/commissural afferents at 0.05 Hz. Slices recovered in the recording chamber at least 1 hr before recordings began. In all experiments, basal field excitatory postsynaptic potentials (fEPSPs) were stable for at least 20 min before the start of each experiment. LTP was induced with either one (E-LTP) or three 1 sec 100-Hz high-frequency stimulation (HFS) trains, with an intertrain interval of 60 sec. To induce mGluR-LTD, slices were incubated with DHPG (100 μM) for 10 min. After induction of either E-LTP or mGluR-LTD, we collected fEPSPs for an additional 60 min (180 min for L-LTP). Slope values were compared from the conditional NMD mutant mice and their control littermates.

### Stereotaxic injection of virus and CA1 spine density analysis

To measure hippocampal spine density, the spine density of CA1 hippocampal neurons in control (CTRL: *UPF2*^wt/wt^;α*CaMKII*:*CreER^T2^*) and Conditional Knockout (CKO: *UPF2*^fl/fl^;α*CaMKII*:*CreER^T2^*) mice was examined post-labelling with GFP-expressing virus. To target the dorsal CA1 field, the stereotaxic coordinates AP: −2, ML: 1.6, DV: 1.5-.25 were used. 1 μl of AAV-*CaMKIIa*-*eGFP* (Addgene, #50469-AAV5) virus was injected bilaterally using a 10 μl nanofil syringe (World Precision Instruments) fitted with a 33-gauge beveled needle. Post-surgery, wounds were closed with tissue adhesive (Vetbond, 3M). Mice were transcardially perfused 7 days later with 4% PFA (without PBS flush) brains removed, sectioned at 200 μm and apical CA1 dendrites imaged on an Olympus FluoView-FV1000 confocal microscope. All spine quantifications were completed in ImageJ.

### Behavioral assays

To study the role of NMD in cognitive function, we induced Cre in 2-month old *UPF2* CKO mice as well as in their control littermates by tamoxifen. This experimental design ensured that phenotypes are not caused by an acute effect of tamoxifen, or a non-specific effect of Cre at cryptic LoxP sites. A 2-week washout period followed tamoxifen dosing before behavior began, at approximately 3-months of age. The Y-maze [13, 14], fear conditioning [14, 15], Morris water maze [16, 17], marble burying [18], grooming (12), locomotor hyperactivity/open field [19], elevated-plus maze [19, 20] and three-chamber social interaction assay [21] were completed and analyzed as previously described, except where outlined below. Briefly, the Y-Maze comprised exposing mice to 2 of 3 arms of the maze for 10 min as a training trial. The third, blocked, arm of the maze was designated as the novel arm, and was pseudorandomized between mice. Following a 1 hr delay, mice were returned to the maze but were free to explore all 3 arms for 5 min. The time spent exploring the novel arm, relative to the other familiar arms, was quantified as our index of short-term spatial memory. To examine associative memory, we utilized a simple 2-day fear conditioned memory protocol. On day 1, mice were exposed to a 6 min conditioning trial that involved 3 tone-shock pairings. A 30-sec tone duration and 30 sec ISI was utilized, while all shock stimuli comprised a 0.7mA scrambled foot-shock that co-terminated with tone presentation. Learning was evaluated by plotting % freezing to each successive tone presentation as a time-course. On day 2, mice were returned to their conditioning context for 6 min, with % freezing over the test session being quantified as contextual fear memory. All experiments were completed in Colbourn fear chambers. The Morris water maze was used to test more sophisticated learning and longer-term spatial memory. Learning was measured by training mice to find a hidden platform (four trials per day, 75 sec/trial) over 4 consecutive days of testing. Trials on each day of testing were averaged, and plotted as a time-course, with gradually decreasing escape latencies being used as our index of spatial learning behavior. To assess spatial memory, we conducted a probe trial 24 hr after the last day of training on day 5. In this trial, the hidden escape platform was removed from the pool. Latency to enter the hidden platform zone, and time spent exploring the target quadrant, were quantified as measurements of spatial memory. Reversal learning was not evaluated as *UPF2* CKO mice did not display adequate learning behavior, thus making it impossible to distinguish a general learning deficit from a *reversal* learning deficit even with addition of extra training trials. Assays were performed in order of least stressful (locomotor hyperactivity/open field) to most stressful (Morris water maze). Mice were bred and tested across 4-5 small cohorts, ensuring that all phenotypes were independent of idiosyncratic effects or nuisance variability that may contaminate any single day of testing.

### Data analysis

Where total protein levels were measured via immunostaining, Image J was used to measure total fluorescence. In these analyses, a defined area containing dendrites were identified and the integrated fluorescent density was measured in each region. Background fluorescence was measured by selecting a region adjacent to dendrites, and was subtracted from positive fluorescent signal-measurements to derive corrected total fluorescence. Depending on the test, behavior was analyzed using a between-groups or mixed-model ANOVA with genotype and sex added as main effects. For tests that involved multiple observations, data were corrected using Tukey’s method. Consistent with prior research [10, 14, 22], as no significant interaction involving the main effect of sex was observed on any given behavioral assay - explicating no modulatory effect of sex on genotype-dependent effects - data from mice were pooled to increase power. No specific exclusion criteria were applied to the datasets, excepting the removal of outliers (defined as values falling outside of ± 2 S.D.). Significance was set at *p*=0.05 per Fisher’s tables. Data analysis was completed using the Graphpad Prism software package.

## Supplementary Figures

**Figure S1.**
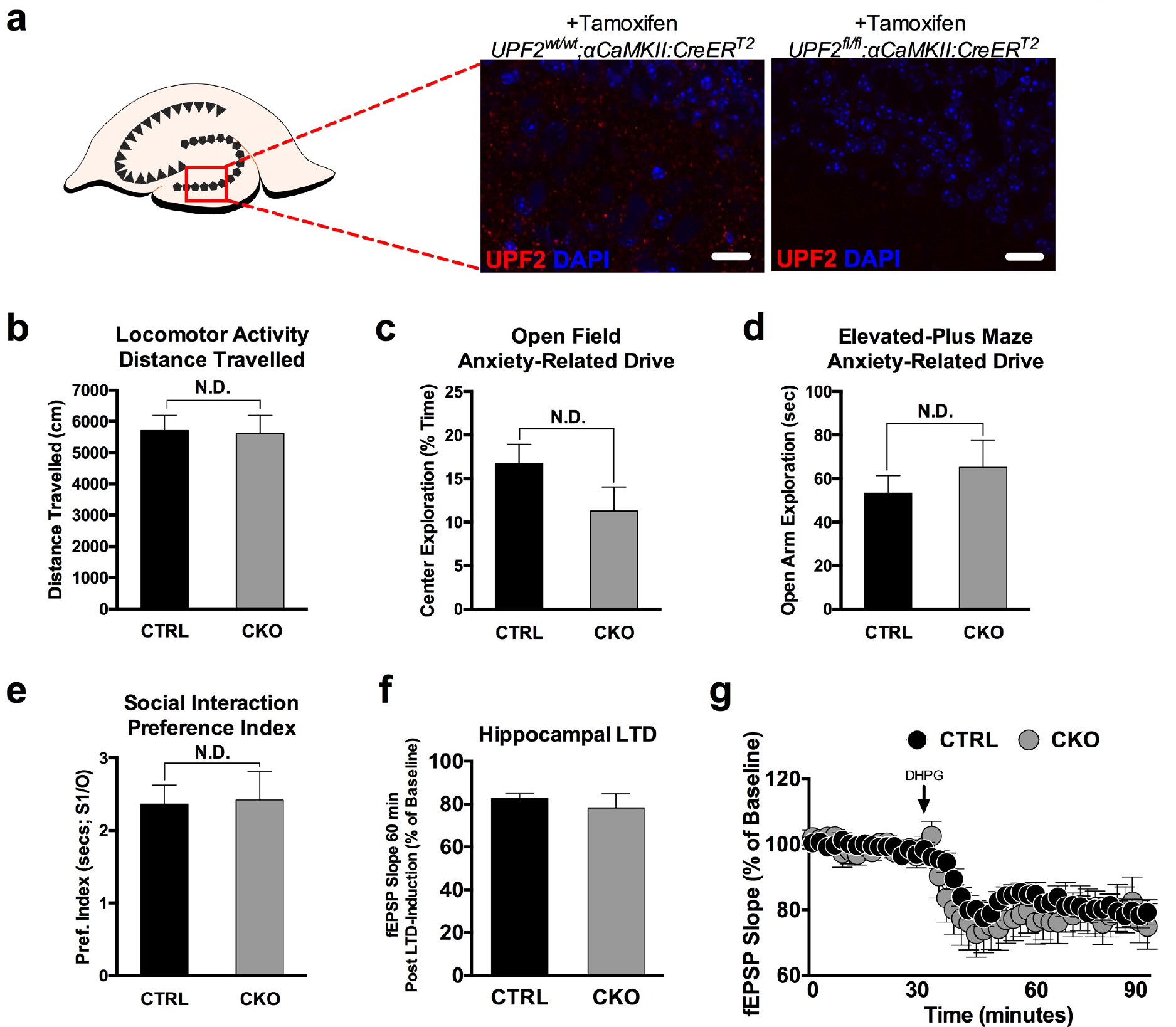
NMD does not influence baseline locomotor activity, anxiety-related behavior, sociability, and LTD in mice. **a**, To determine the potential role of NMD in hippocampus-dependent cognition in the adult brain, we disrupted NMD in postmitotic neurons using a conditional knockout mouse of *UPF2* (*UPF2*^fl/fl^) [1]. To temporally control *UPF2* disruption, we crossed *UPF2^fl/fl^* mice with a mouse line that expresses tamoxifen-inducible Cre under the α*CaMKII* promoter (α*CaMKII*::*CreER^T2^*) [2]. Before assessing synaptic plasticity and learning and memory (Figure 1) in *UPF2*^fl/fl^;α*CaMKII*:*CreER^T2^* (CKO) and *UPF2*^wt/wt^;α*CaMKII*:*CreER^T2^* (CTRL) mice, we confirmed that conditional deletion of *UPF2* gene (using 200 mg/kg Tamoxifen, for 5 doses) led to the successful loss of UPF2 protein in the hippocampus. **b-d**, To determine whether NMD induces hyperactivity, mice were subjected to a 2 hr locomotor hyperactivity assay in photocells. During this 2 hr trial, the distance that mice travelled in cm was analyzed. Greater distances travelled are used as an indicator of locomotor hyperactivity, which may confound behavioral testing on other maze tasks. Similarly, this assay can also assess hypoactivity induced by genetic alterations, which may also influence task performance. **b**, In our assessment of baseline locomotor activity, there were no genotype differences between CTRL (n=25) and CKO (n=22) mice. This indicates that loss of UPF2 does not result in acquired hyper- or hypoactivity. **c**, A 10 min open field assay was also conducted in locomotor photocells as an additional control assay, with less time spent exploring the center field of photocells being interpreted as an anxiety-like phenotype. Consistent with the outcome of the locomotor activity assay, no evidence of anxiety-like behavior between CTRL (n=25) and CKO (n=22) mice emerged on this test, indicating that NMD does not modify anxiety-related behavior. **d**, The elevated-plus maze was used as a control assay to ensure that NMD did not modify anxiety-related exploratory drive, which may compound measurements of learning and memory. Briefly, mice were subjected to a 10 min test session, with time spent exploring the open arms being used as an index of anxiety-like behavior. No genotype differences emerged between CTRL (n=25) and CKO (n=22) mice on this test. **e**, Alterations in both spine density and GluR1 levels are linked to deficits in sociability [3, 4]. In addition, defects in the NMD machinery are linked to numerous neurodevelopmental diseases associated with alterations in normal social behavior [5–10]. To determine whether NMD is required for sociability, we used the three-chamber social interaction task and measured sociability in CTRL and CKO mice. Mice were habituated to the empty three-chamber apparatus for 10 min before being exposed to a stranger mouse placed inside a cup in one chamber and an empty cup in the opposite chamber for an additional 5 min. A preference index was derived from time spent sniffing or interacting with the cup containing the stranger mouse divided by time spent interacting with the empty cup, with higher preference indexes representing more social behavior. No significant differences in preference index emerged between CTRL (n=24) and CKO (n=19) mice on the three-chamber social interaction assay. Data are represented as mean ± SEM; N.D. represents “No Difference”. Scale bar: 15 μm. **f-g**, NMD regulates spine density and L-LTP in hippocampus (Figure 1j-m). We also assayed LTD in *UPF2* CKO. Because of age-related decline in LTD, we induced Cre at 3 weeks of age and hippocampal slices were acquired from 6 week-old CTRL and CKO mice (see Methods). LTD was induced by acute application of 100 μM DHPG for 10 min to hippocampal slices. fEPSPs were collected for 60 min post-LTD induction. Compared to CTRL mice (n=14 slices from 7 mice), CKO mice (n=10 slices from 5 mice) did not show significantly different slope values post-DHPG application indicating that LTD is unaltered in the absence of NMD in hippocampus

**Figure S2.**
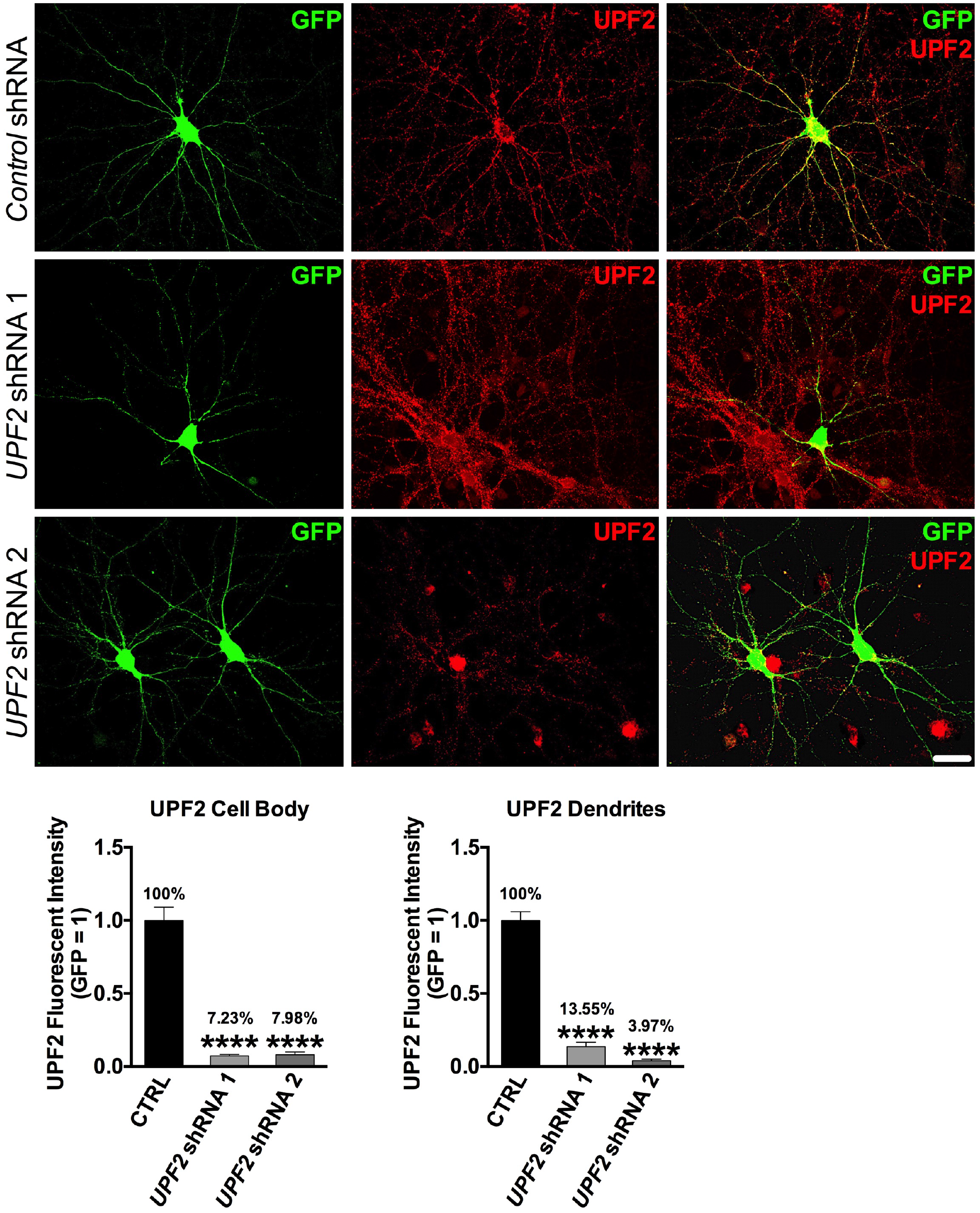
Knockdown of *UPF2* in mouse hippocampal neurons by *UPF2*-shRNA:GFP lentivirus. To disrupt NMD, we targeted the *UPF2* mRNA using *UPF2*-shRNA:GFP lentivirus. Canonical NMD involves the interaction of UPF1, UPF2 and UPF3 proteins. While UPF1 function is not restricted to the NMD pathway [11], UPF2 has been successfully used to disrupt NMD in several studies [1, 12, 13]. We infected E16 hippocampal neurons at DIV7 with *control*- or two independent *UPF2*-shRNA viruses. To assess the knockdown of UPF2, we fixed the cells at DIV14 and performed immunohistochemistry for UPF2. Application of both *UPF2*-shRNA viruses resulted in a robust depletion of UPF2 within infected neurons and their dendrites, while UPF2 expression remained intact in scrambled *control*-shRNA cultures and non-infected neurons in *UPF2*-shRNA cultures (UPF2+, GFP-cells in *UPF2*-shRNA 2 panel). This control experiment with two separate viruses targeting non-overlapping UPF2 sequences exemplifies specific knockdown of *UPF2* across our studies, and ensures that the phenotypes that arise from this knockdown are unlikely off-target effects. Scale bar: 30 μm.

**Figure S3.**
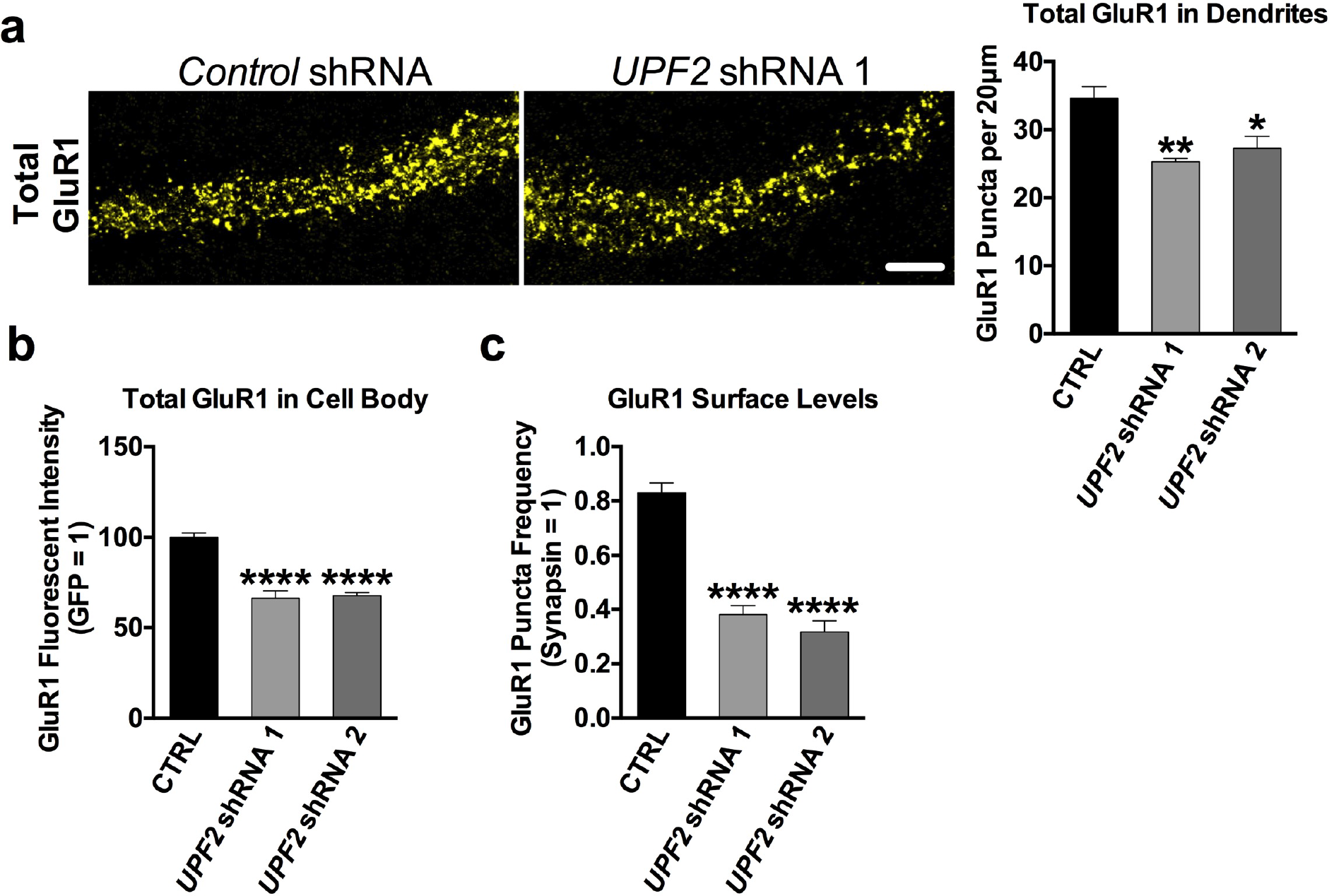
Total levels and surface levels of GluR1 protein are decreased upon knockdown of *UPF2*. Loss of *UPF2* led to a decrease in GluR1 levels *in vivo* (Figure 1m). To explore this phenotype *in vitro*, we infected hippocampal neurons with the 2 independent *UPF2*-shRNA:GFP lentivirus vectors validated in Figure S2 to interrupt endogenous NMD activity. Hippocampal neurons were infected at 7 days *in vitro* (DIV7) and GluR1 expression was examined on DIV21. We quantified total GluR1 signal using an anti-C-terminus-GluR1 antibody in fixed and permeabilized cells. **a**, UPF2-deficient dendrites had significantly lower GluR1 density compared to control dendrites (n=3 biological replicates per *control*-shRNA [10 neurons, 23 dendrites] and per *UPF2*-shRNA [11 neurons, 25 dendrites]; representative image reflects *UPF2* shRNA 1, and *UPF2* shRNA refers to the UPF2 shRNA 1 in the following text). This suggests that while NMD does not alter *GluR1* transcription or mRNA degradation (Figure 2d), there is either an increase in GluR1 receptor degradation or a decrease in GluR1 synthesis. **b**, Total GluR1 levels were also significantly decreased in the cell bodies of UPF2-deficient neurons. We measured total fluorescence signal by measuring the integrated density in a defined region using ImageJ. Knockdown of *UPF2* led to a decrease in total GluR1 levels in cell bodies (n=3 biological replicates per *control*-shRNA [15 neurons] and per *UPF2*-shRNA [13 neurons]). **c,** Surface GluR1 expression was also quantified, as described both in-text and methods, using both validated *UPF2*-shRNA:GFP lentivirus vectors. Similar to total GluR1 levels in dendrites, surface levels of GluR1 were also decreased in UPF2-deficient dendrites (n=3 biological replicates per *control*-shRNA [15 neurons] and per *UPF2*-shRNA [13 neurons]). Increased local internalization and decreased local synthesis of GluR1 are likely to contribute to this phenotype (see Figure 3). Data are represented as mean ± SEM; **p < 0.01, and ***p < 0.001. Scale bar: 20 μm.

**Figure S4.**
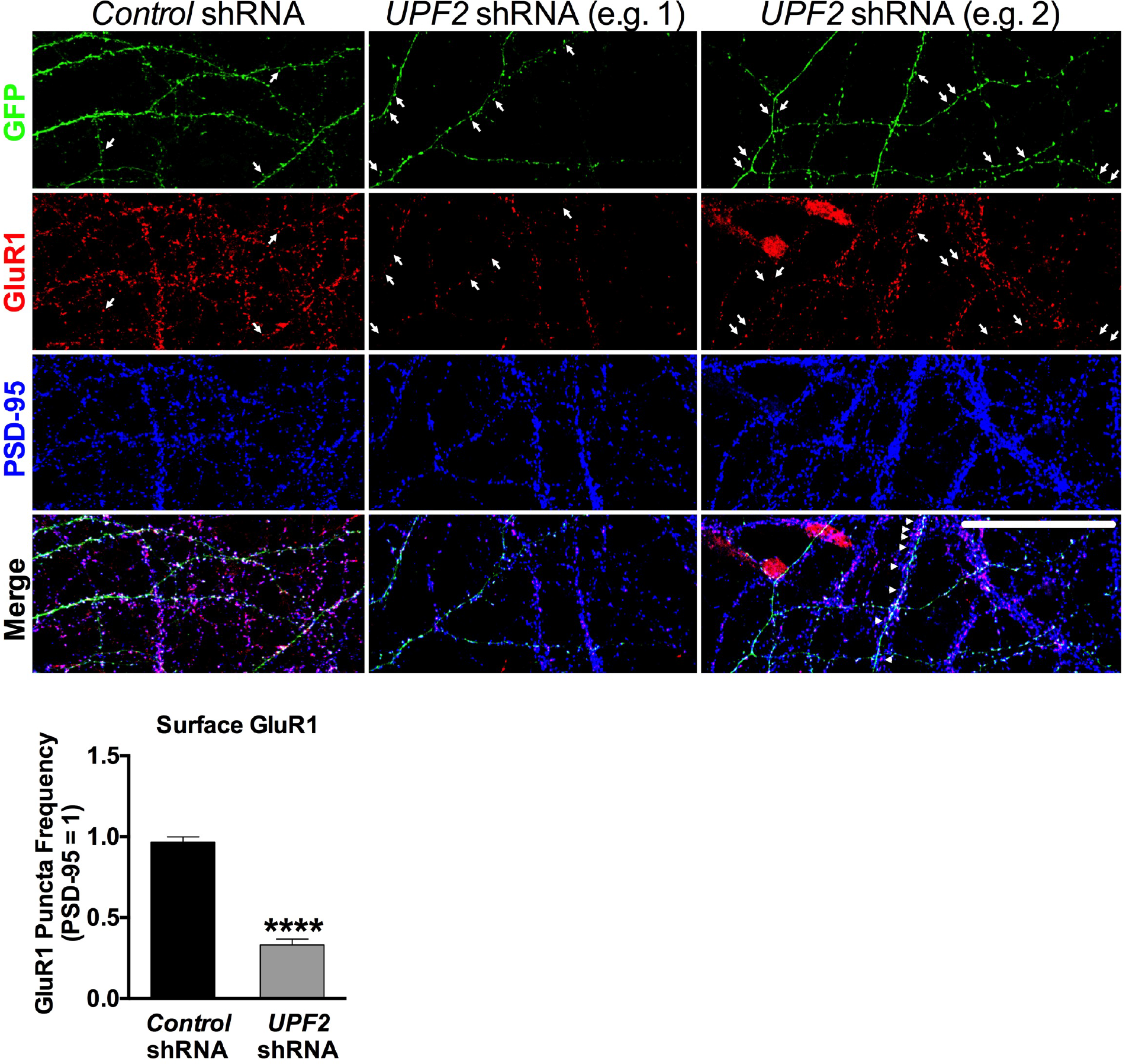
Decrease in GluR1 surface levels in UPF2-deficient dendrites was also recapitulated when the synaptic marker PSD-95, in addition to synapsin, was used for quantifications. In Figure 2, we show that knockdown of *UPF2* in hippocampal neurons led to a robust depletion of surface GluR1 levels without altering the mRNA transcription of this receptor. For these quantifications, we used synapsin as a normalization marker as this is consistent with prior work (see Methods). To address whether this GluR1 phenotype is recapitulated when another synaptic marker is used for normalization, we repeated this experiment using the postsynaptic marker PSD-95. Similar to the synapsin quantifications, the disruption of UPF2 in hippocampal neurons resulted in a significant depletion of surface GluR1 levels when synapses were identified and normalized against PSD-95 expression (n=3 biological replicates per group; 11 dendrites from 7 neurons for *control*-shRNA cultures, and 12 dendrites from 8 neurons for *UPF2*-shRNA cultures). Arrows show examples of GFP and GluR1 colocalization in spines. Two panels of *UPF2*-shRNA dendrites are presented to exemplify contrasting aspects of GluR1 expression. Arrows depict the loss of surface GluR1 in spines of *UPF2*-shRNA infected hippocampal neurons relative to control cultures. Arrowheads in the merged panel of *UPF2*-shRNA example 2 depict a non-infected dendrite segment with intact GluR1 expression that is overlapping an infected dendrite that exhibits reduced GluR1 colocalization in isolated GFP+ spines. These data replicate the depletion of GluR1 surface levels in UPF2-deficient dendrites presented in Figure 2, and provide confirmation that synapsin produces concordant data to PSD-95 when used to evaluate surface GluR1 levels. Data are represented as mean ± Standard Error of the Mean (SEM); ****p < 0.0001. Scale bar: 30 μm.

**Figure S5.**
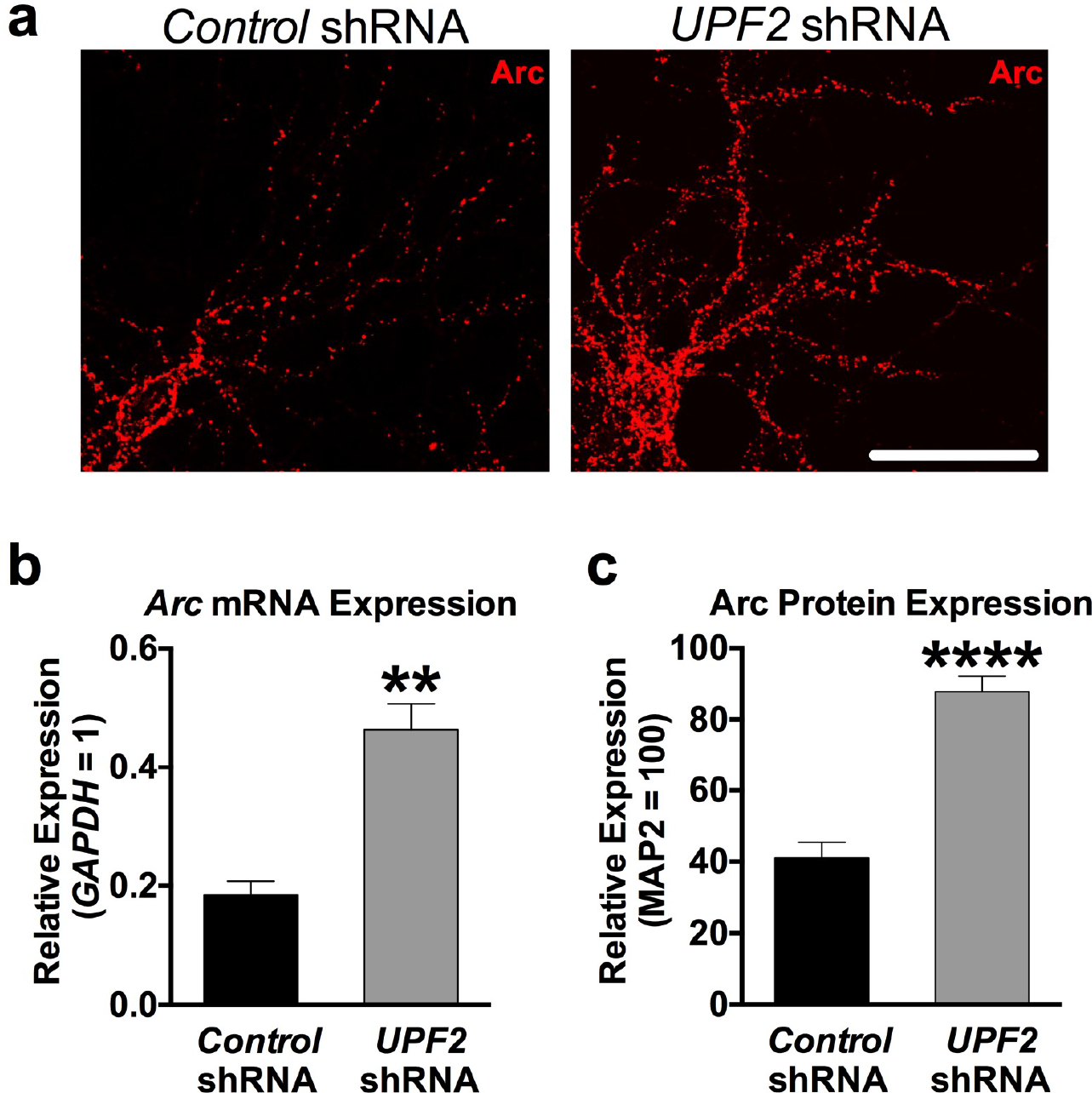
Arc levels are increased upon knockdown of *UPF2* in hippocampal neurons. **a**, Arc immunostaining in mouse hippocampal neurons upon knockdown of *UPF2*. Disruption of NMD caused a significant reduction in the total (Figures 1 and S3) and surface (Figure 2) levels of GluR1 without affecting the *GluR1* mRNA levels (Figure 2d). This suggests that NMD regulates GluR1 surface expression through other mechanisms than transcription or *GluR1* degradation. GluR1 signaling is modulated by dynamic insertion and removal of the GluR1 receptor to the synaptic surface [14]. The synaptic plasticity protein Arc drives the removal of GluR1 from the surface of synapses. *Arc* mRNA is a known target of NMD. Both *Arc* mRNA and Arc protein are increased upon disruption of NMD [12, 15] suggesting that elevated Arc levels may contribute to the reduced surface expression of GluR1. To confirm that knockdown of *UPF2* alters endogenous Arc levels, we cultured E16 mouse hippocampal neurons and infected with *control*- or *UPF2*-shRNA lentivirus at DIV7 to disrupt NMD. At DIV21, we performed immunostaining for Arc protein. **b**, Quantitative RT-PCR analysis of *Arc* mRNA in UPF2-deficient neurons. To quantify *Arc* mRNA, we performed qRT-PCR. *GAPDH* was used as a control transcript. Expression of *Arc* was presented relative to *GAPDH*. Neurons infected with *UPF2*-shRNA lentivirus displayed exaggerated *Arc* mRNA expression (n=3 biological replicates per group) compared to control neurons. **c**, Quantifications of Arc protein in UPF2-deficient neurons. Quantifications of fluorescent intensity, normalized to MAP2, showed that Arc protein levels are also increased in neurons with disrupted UPF2 (per group, n=10 dendrites for analysis of fluorescent intensity). Together, these data confirm that Arc levels are increased upon disruption of UPF2. Arc is known to induce internalization of GluR1 receptor [16]. However, the total levels of GluR1 are independent of its internalization rate. While Arc-mediated internalization of GluR1 influences surface levels, it does not influence total GluR1 levels [17]. This suggests that while loss of UPF2 does not alter *GluR1* transcription or mRNA degradation (Figure 2d), there is either an increase in GluR1 receptor degradation or a decrease in GluR1 synthesis. Data are represented as mean ± Standard Error of the Mean (SEM); ** p < 0.01, **** p < 0.0001. Scale bar: 30 μm.

**Figure S6.**
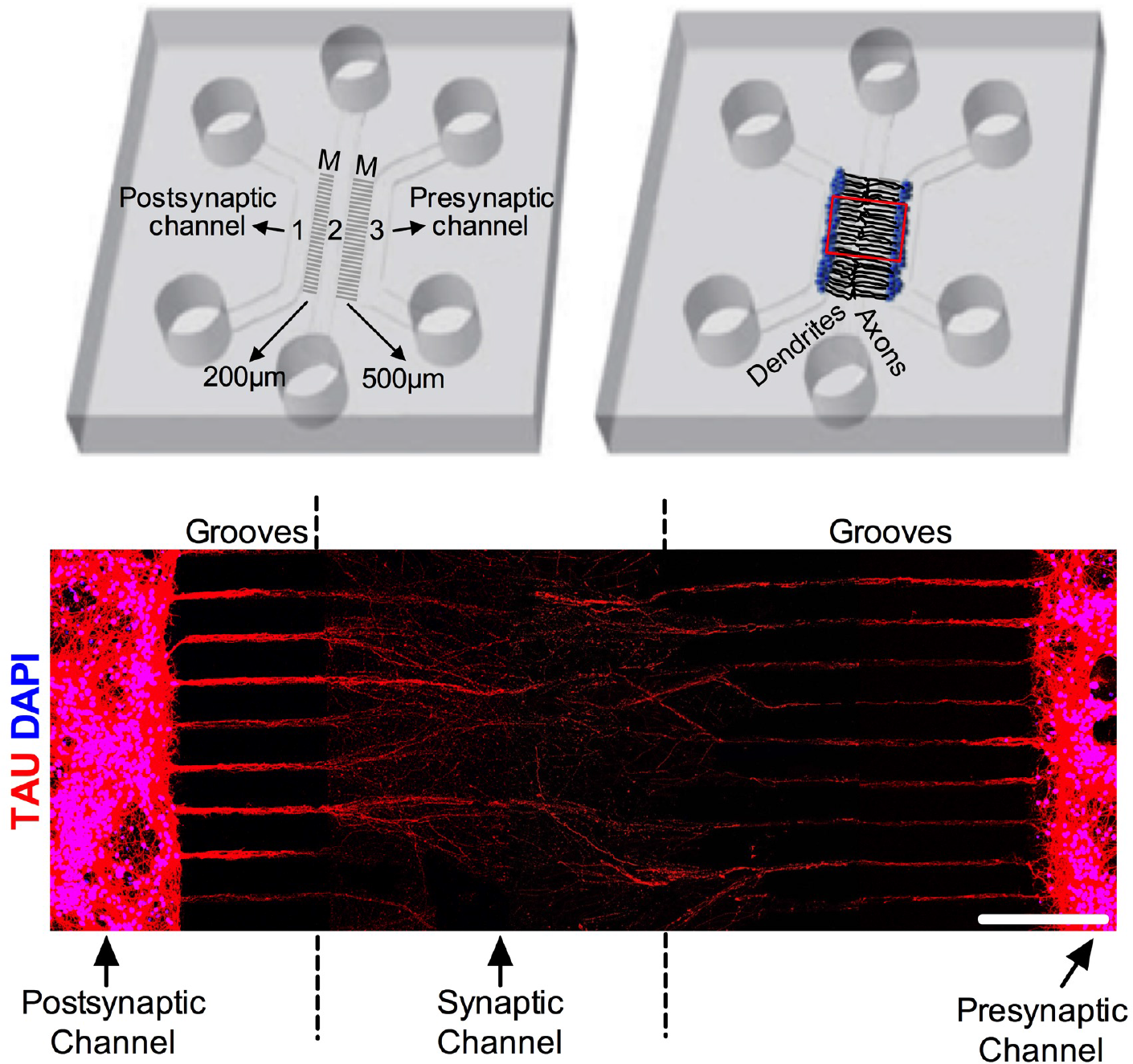
Axonal penetrance of microgrooves in tripartite microfluidic devices. As shown in the schematic presented here, and the MAP2 immunostaining presented in Figure 3, our custom-engineered tripartite microfluidic device only permits the dendrites of neurons from left channel (channel 1) to penetrate the middle channel, which is referred to as the “synaptic channel” (channel 2). Here, TAU staining shows that axons from both channels extend to the synaptic channel suggesting that axons of both channel 1 and channel 3 can form synapses with dendrites that originated from channel 1. However, due to their limited length, dendrites remain in very close proximity to channel 1 (Figure 3). This tends to restrict dendrite-axon synapses arising from neurons in channel 1 to the proximal-most area of the synaptic channel. Because axons are very long, and the short microgrooves that separate channels 1 and 2 are just 200 μm in length, most channel 1 axons span the entire synaptic channel and go-on to penetrate channel 3. Thus, the majority of the synapses formed in the synaptic channel consist of dendrites projecting from channel 1 and axons originating from channel 3, referred to as the “presynaptic channel”. UPF2 is not expressed in mature axons (Figure S10), and only dendrites from neurons plated in channel 1 can reach the synaptic channel (Figure 3). Therefore, regardless of where the presynaptic axons project from, or where synapses are formed within the synaptic channel, the manipulation of NMD exclusively in channel 1 is confined to postsynaptic dendrites. Scale bar: 200 μm.

**Figure S7.**
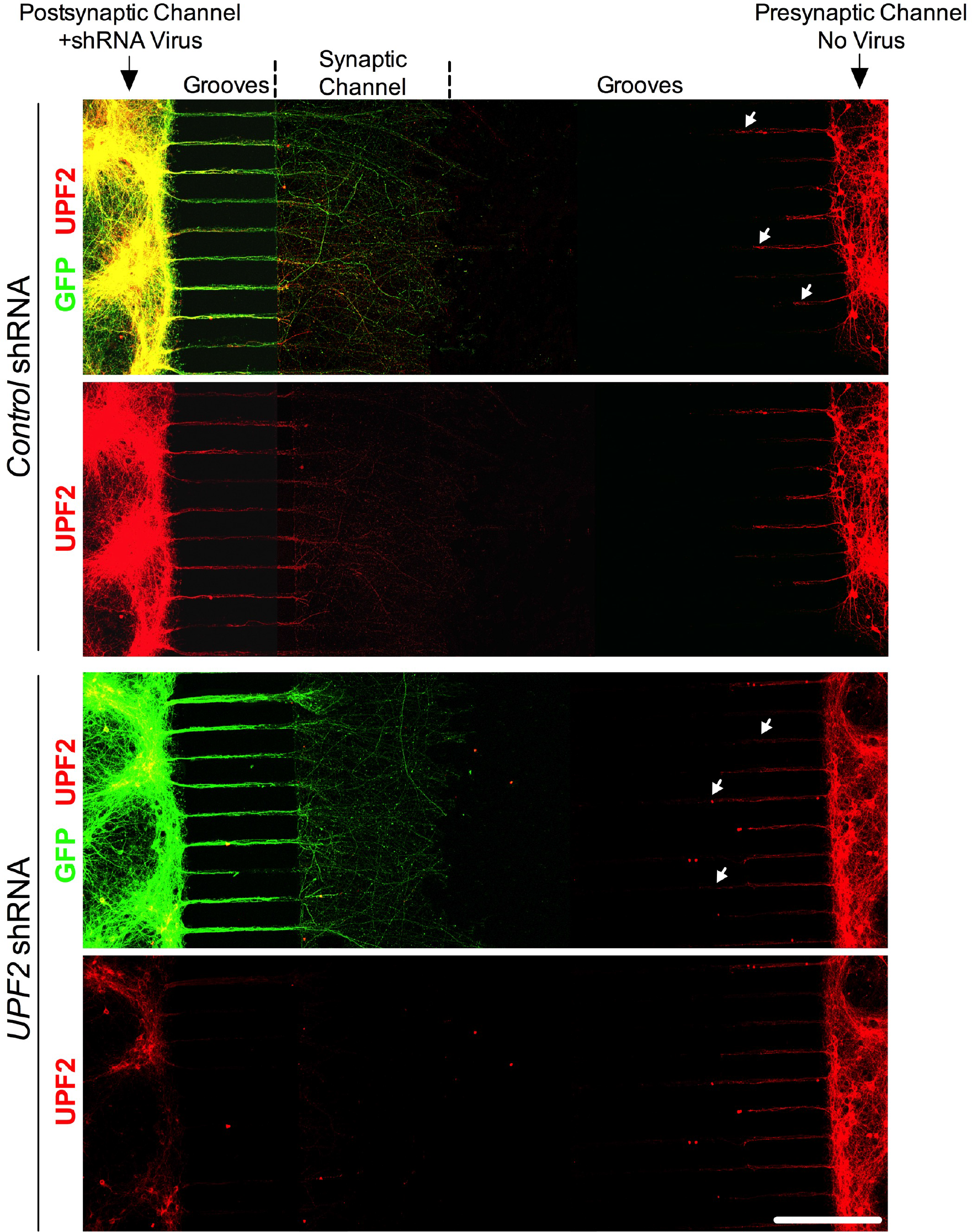
Application of *UPF2* shRNA virus exclusively to the postsynaptic channel results in loss of UPF2 in this channel, as well as in dendrites within the synaptic channel, but not in presynaptic channel neurons or their projections. The physical and fluidic isolation of channels is a fundamental feature and component of microfluidic devices, as it permits the selective manipulation/isolation of neuronal populations and compartments. To address the role of NMD in dendrites in synaptic function, we selectively treated postsynaptic channels with *UPF2*-shRNA virus at DIV7. To determine whether the virus application resulted in selective loss of UPF2, we performed immunostainings for UPF2 following fixation of cells at DIV13. Consistent with the fluidic isolation of channels, the GFP signal was specific to infected neurons restricted to the postsynaptic channels and their neurites in the synaptic channel. Selective application of the *UPF2*-shRNA virus in the postsynaptic channel resulted in nearly complete loss of UPF2 protein in this channel, as well as in dendrites in the synaptic channel, but not in the presynaptic channel. Because mature axons do not express UPF2 (Figure S10), the UPF2-positive signal in the synaptic channels in the control experiment solely reflects dendritic expression. Consistent with this, dendrites that emerged from the presynaptic channel, which only traveled half way through microgrooves (white arrows), also retained their UPF2 expression in both the *control*- and *UPF2*-shRNA experiments (see Figure 3 and Figure S6 for the comparison of dendrite and axon projection in these grooves). In all experiments that employed *control*- or *UPF2*-shRNAs in tripartite devices, viruses were only applied to the postsynaptic cell body channel of microfluidic devices. Scale bar: 200 μm.

**Figure S8.**
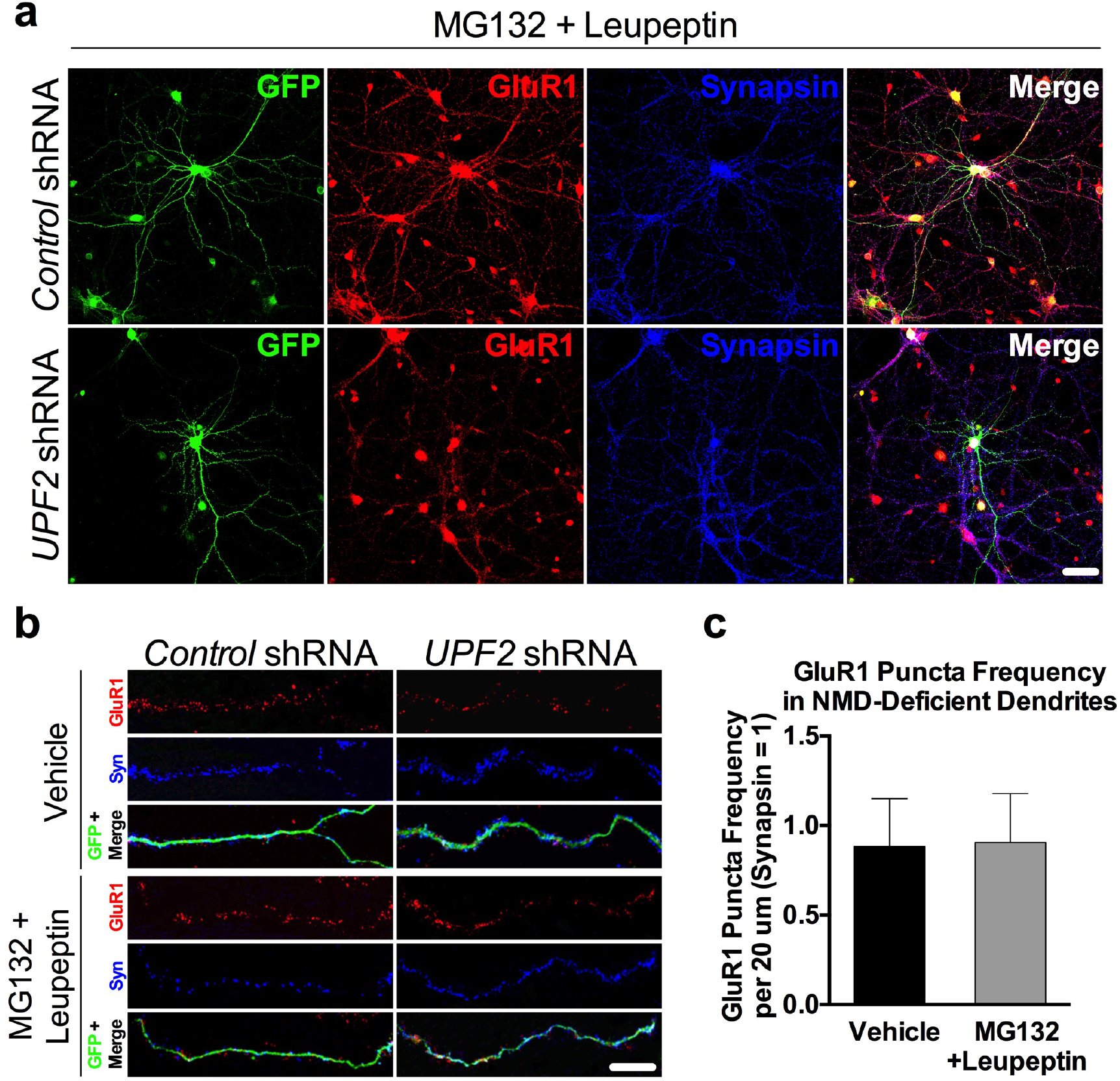
GluR1 degradation is not altered upon disruption of UPF2. Both the surface and total expression of GluR1 in dendrites is reduced upon knockdown of UPF2 (Figure S3 and 2). Although the increase in basal internalization rate of GluR1 (Figure 3a) contributes to the reduction in surface levels of this receptor in the absence of UPF2, it does not explain the decrease in total GluR1 levels [17]. Since the disruption of NMD does not alter *GluR1* transcription or degradation (Figure 2d), an alteration in the balance of local synthesis and degradation of GluR1 is likely to account for the overall reduction in the total levels of this receptor upon disruption of NMD. Therefore, we examined the local synthesis of nascent GluR1 protein (Figure 3) as well as the degradation of GluR1 upon loss of UPF2. Figure 3 shows that the local synthesis of GluR1 is repressed in UPF2-deficient dendrites. To study GluR1 protein degradation, we cultured E16 mouse hippocampal neurons in either 24 well plates or in the cell body compartments of tripartite chambers and infected with *UPF2*-shRNA lentivirus at DIV7. In the case of tripartite chambers, we only infected postsynaptic cells with the *UPF2*-shRNA lentivirus. Because endocytosed AMPA receptors can undergo either lysosome- or proteasome-mediated degradation [18–22], we targeted both lysosomal and proteasomal degradation simultaneously. At DIV21, proteasomal and lysosomal degradation were inhibited with 10 μM of the protease inhibitor MG132 and 20 μM of leupeptin, respectively, for 6 hr. For the tripartite experiment, proteasomal and lysosomal degradation were inhibited by treating the synaptic channels with 10 μM of the protease inhibitor MG132 and 20 μM of leupeptin, respectively, for 6 hr. To label surface GluR1, neurons were fixed and stained with an anti-Nterminus-GluR1 antibody without permeabilization (see also Methods). **a-b**, Representative low (regular cultures) and high magnification (in tripartite chambers) images of GluR1 surface expression in treated and non-treated UPF2-deficient dendrites. **c,** Quantification of surface GluR1 puncta shows that inhibition of proteasomal and lysosomal degradation did not significantly influence GluR1 puncta frequency in UPF2-deficient dendrites (n=3 biological replicates per group; 11 neurons per group). Data are represented as mean ± SEM. Scale bar: 20 μm.

**Figure S9.**
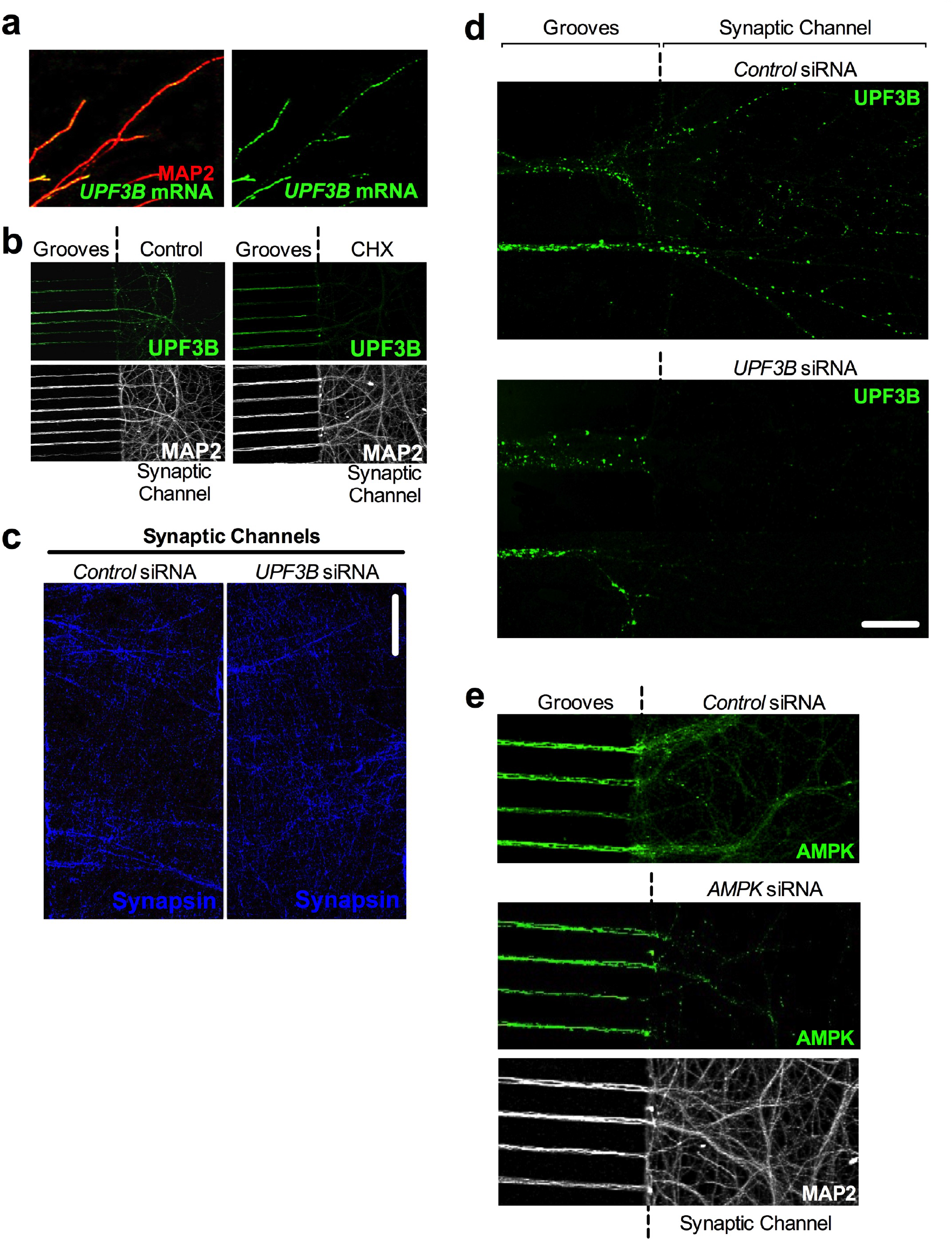
UPF3B is locally synthesized in dendrites. **a-b**, UPF3B is locally synthesized in hippocampal dendrites. Both dendritic internalization and local synthesis of GluR1 receptor are altered upon knockdown of UPF2. In addition, NMD targets *Arc* and *AMPK* are subjected to translation-dependent degradation in dendrites (Figure 4). Although these data suggest that NMD might regulate GluR1 locally in dendrites, there is no direct evidence for the local requirement for NMD in these compartments. To study the specific requirement of locally-occurring NMD in the regulation of GluR1 within dendrites, we devised a strategy for targeting UPF3B protein (which, like UPF2, is also a specific and major component of the NMD machinery) locally within the synaptic channel of tripartite devices (Figure 4d-e). Dendritic transcriptome analysis showed that the mRNA of UPF3B protein is localized to dendrites [23]. To confirm this, we performed Fluorescent In Situ Hybridization (FISH) on hippocampal neurons with antisense riboprobes against *UP3B* mRNA at DIV21. *UPF3B* FISH resulted in punctate labeling (green) along dendrites confirming the localization of *UPF3B* mRNA to these compartments (**a**). Next, we sought to determine if UPF3B is locally synthesized in dendrites. To do this, we inhibited *UPF3B* mRNA translation via dendritic application of Cycloheximide (CHX, 10 μM) and evaluated the UPF3B protein in synaptic channels. We cultured E16 mouse hippocampal neurons in tripartite chambers and, at DIV21, we treated the synaptic channels with CHX for 6 hr. This resulted in an almost complete loss of UPF3B protein in dendrites (**b**). MAP2 was used to visualize dendrites in both (**a**) and (**b**). **c-d**, To determine whether inhibition of local synthesis of UPF3B in synaptic channels alters GluR1 locally, we selectively treated synaptic channels with siRNAs against the *UPF3B* mRNA and evaluated GluR1 levels (Figure 4d-e). Here, we confirmed that while the treatment of synaptic channels with *UPF3B* siRNA-cocktail (10 nM) for 7 days did not cause an alteration in synaptic potential in these channels (**c**), it led to a complete loss of UPF3B protein (**d**). Note that the expression of UPF3B remained intact within dendrites adjacently located within fluidically-isolated microgrooves in panel **d**. **e**, Activated AMPK is known to interrupt translation by either inhibiting the mammalian target of rapamycin (mTOR) kinase or the translation elongation factor 2. These data suggest that NMD might regulate total levels of GluR1 through degradation of *AMPK* mRNA in dendrites. To determine whether local AMPK in dendrites modulates GluR1 levels, we inhibited AMPK synthesis in the synaptic channels of tripartite microfluidic devices and examined total GluR1 levels (Figure 4**f**). To do this, we targeted *AMPK* mRNA by systematic application of siRNAs against the *AMPK* mRNA to synaptic channels. Application of 10 nM of two non-overlapping *AMPK*-siRNAs to synaptic channels for 7 days, starting at DIV14, led to selective depletion of AMPK in dendrites in synaptic channels but not in dendrites in microgrooves (see middle panel). Data are represented as mean ± SEM. Scale bar: **c** 20 μm, **d** 75 μm.

**Figure S10.**
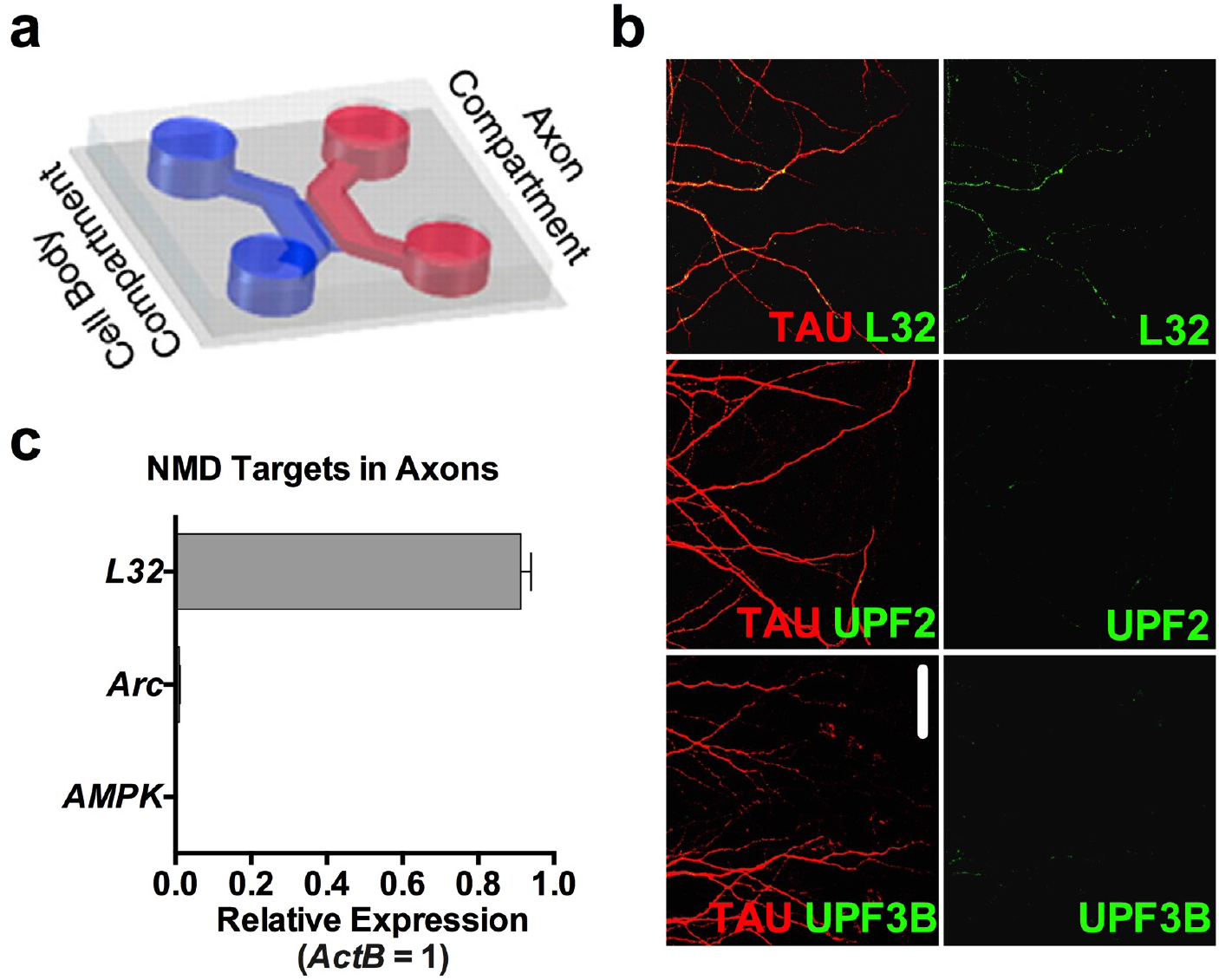
The NMD machinery and NMD targets are not expressed in mature axons. **a-c**, The synaptic channel of tripartite chambers contains both dendrites and axons, raising the possibility that some of the phenotypes observed upon loss of local UPF3B or AMPK in synaptic channels might arise from affected axons. To clarify this, we addressed whether the NMD machinery, as well as *Arc* and *AMPK*, are also present in mature axons in addition to dendrites. We cultured E16 neurons in two-partite microfluidic chambers (**a**). Unlike tripartite chambers used in other experiments, two-partite chambers allow physical isolation of axons [12]. At DIV21, we collected the axonal material and measured the levels of *Arc* and *AMPK* mRNAs by qRT-PCR (**c**). While the axonal mRNA *L32* was readily detected, mature hippocampal axons lacked *Arc* and *AMPK* mRNAs (n=3 biological replicates). This is consistent with previous reports that the axonal transcriptome significantly shrinks as axons mature [24] and does not include any known NMD targets [25, 26]. Similarly, unlike navigating axons [12], mature axons did not contain the major proteins of the NMD machinery (**b**). These data suggest that the GluR1 phenotypes observed upon application of siRNAs against both *UPF3B* and *AMPK* in the synaptic channels of tripartite chambers solely originate from dendrites and not axons. Data are represented as mean ± SEM; Scale bar: 30 μm.

**Figure S11.**
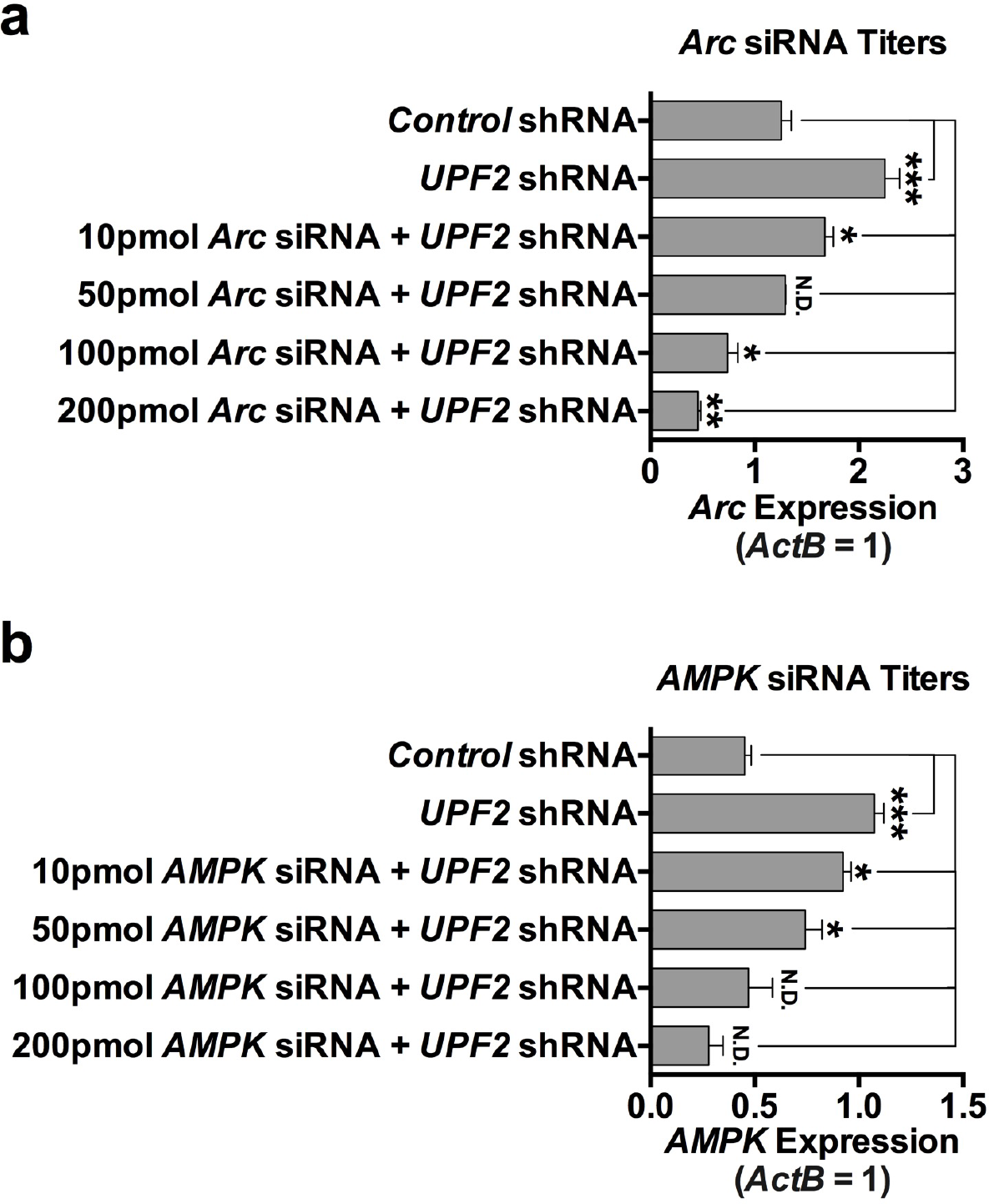
*Arc* and *AMPK* titers for siRNA normalization of exaggerated expression in NMD-deficient dendrites. **a-b**, We have identified *Arc* and *AMPK* as candidate targets in the NMD-mediated regulation of GluR1 levels within the dendrites and synaptic compartments of hippocampal neurons. We next sought to functionally determine which of these molecules is responsible for depletion of surface GluR1 in UPF2-deficient dendrites. To do this, we systematically modulated the elevated levels of *Arc* and *AMPK* in dendrites by siRNA transfection of synaptic channels that contain UPF2-deficient dendrites (Figure 5). As siRNA treatment affects mRNA expression in a concentration-dependent manner [27–30], we first determined the amount of siRNA required to correct the levels of each target in UPF2-deficient dendrites presented in Figure 5. First, we applied the *UPF2*-shRNA virus to the postsynaptic-cell channels of tripartite microfluidic devices at DIV7. Starting at DIV14, we treated the synaptic channels of UPF2-deficient dendrites with non-overlapping siRNAs (2 independent siRNAs per mRNA; see Methods) at different concentrations for either *Arc* (**a**) or *AMPK* (**b**). To deliver siRNAs, we used 10% NeuroPORTER (Sigma). At the end of the treatment, synaptic channels were perfused with TRIzol and lysates used for quantitation of *Arc* and *AMPK* mRNA by qRT-PCR. A panel of *Arc* or *AMPK* siRNA-cocktail titers was selectively applied to the synaptic channels of tripartite chambers (n=3 replicates per mRNA and group). As expected, infection of neurons with *UPF2*-shRNA virus significantly increased the expression of *Arc* (**a**) and *AMPK* (**b**). **a**, Subsequent treatment with *Arc* siRNA cocktail at 10 pmol led to a significant decrease in the expression of *Arc* mRNA in the dendrites of UPF2-deficient postsynaptic neurons, but was still elevated relative to levels observed in the dendrites of postsynaptic neurons in control cultures. Similarly, 100 pmol and 200 pmol titers of *Arc* siRNA cocktail led to a significant downregulation of *Arc* mRNA in the dendrites of UPF2-deficient postsynaptic neurons, but produced expression levels lower than that required to recapitulate the physiological profile of dendritic *Arc* mRNA expression observed in control cultures. However, we observed that 50 pmol of *Arc* siRNA cocktail normalized the expression of *Arc* mRNA in the dendrites of UPF2-deficient postsynaptic neurons, and that these cultures were indistinguishable from control cultures, leading us to adapt this titer in experiments Figure 5. **b**, Treatment of the synaptic channels of tripartite chambers with 10 pmol and 50 pmol titers of *AMPK* siRNA cocktail significantly reduced the overexpression of *AMPK* in the dendrites of UPF2-deficient postsynaptic neurons, but did not restore expression levels to those observed in control cultures. 100 pmol *AMPK* siRNA cocktail titer was chosen for rescue experiments in Figure 5, as raw values closely resembled and statistically did not differ from those observed in the control group (n=3 replicates). Data are represented as mean ± SEM; * p < 0.05, ** p < 0.01, *** p < 0.001, N.D. indicates “No Difference”.

**Figure S12.**
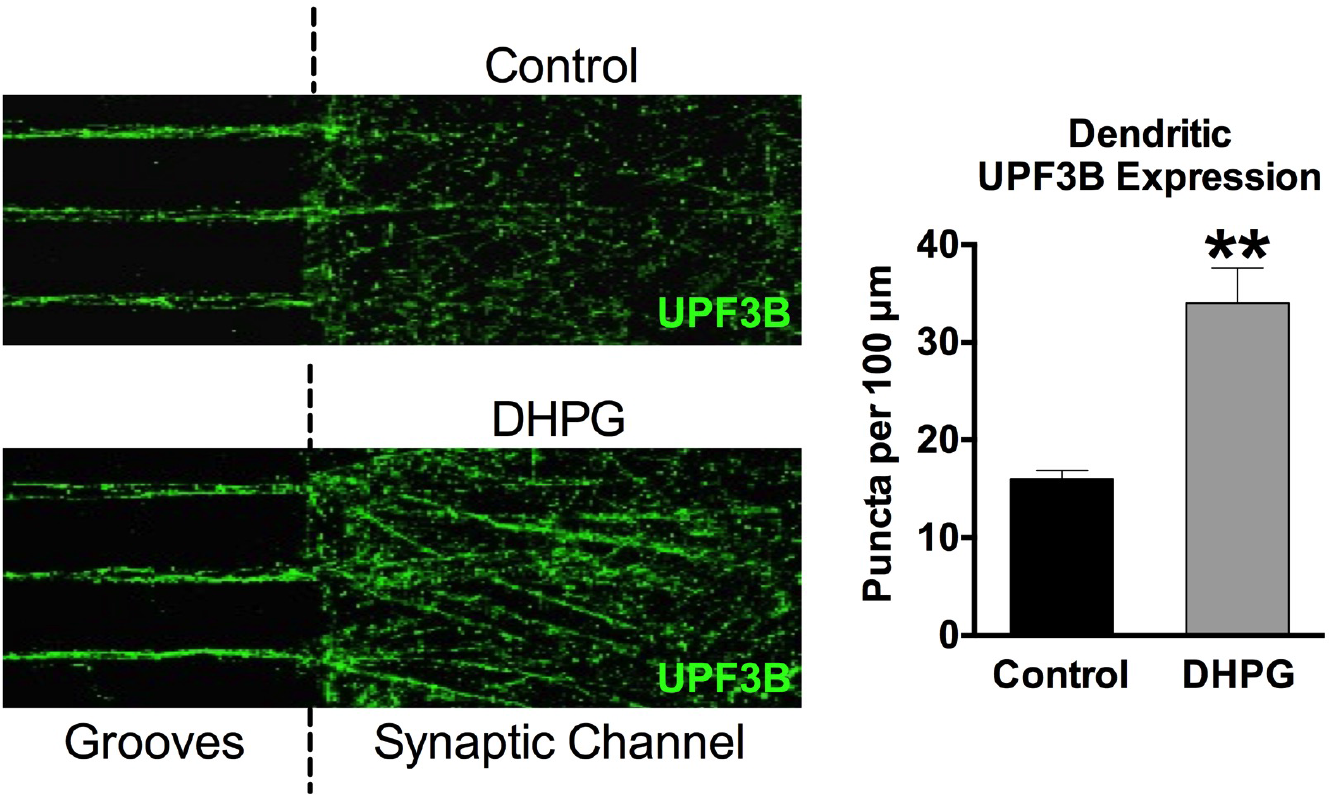
Neuronal activity upregulates local UPF3B expression in dendrites. The mRNA of UPF3 protein is trafficked to [23] and locally translated in dendrites (see Figure S9). Local translation of selected mRNAs is induced by activity in dendrites [31], suggesting that NMD might be regulated at the level of translation in these compartments. To test this, we stimulated synapses with the mGluR agonist DHPG and measured the levels of UF3B in dendrites. E16 mouse hippocampal neurons were cultured in tripartite microfluidic devices and, at DIV21, the mGluR agonist DHPG (100 μM) was selectively applied to synaptic channels for 5 min to induce synaptic activity. Following fixation of cells, immunostaining for UPF3B was performed in control (vehicle) and DHPG-treated dendrites. Quantification of UPF3B puncta within 100 μm dendrite segments revealed a significant increase in UPF3B expression following treatment with DHPG relative to vehicle-treated cultures (n=3 biological replicates per group). This suggests that NMD is induced by neuronal activity in dendrites. Data are represented as mean ± SEM; ** p < 0.01.

**Figure S13.**
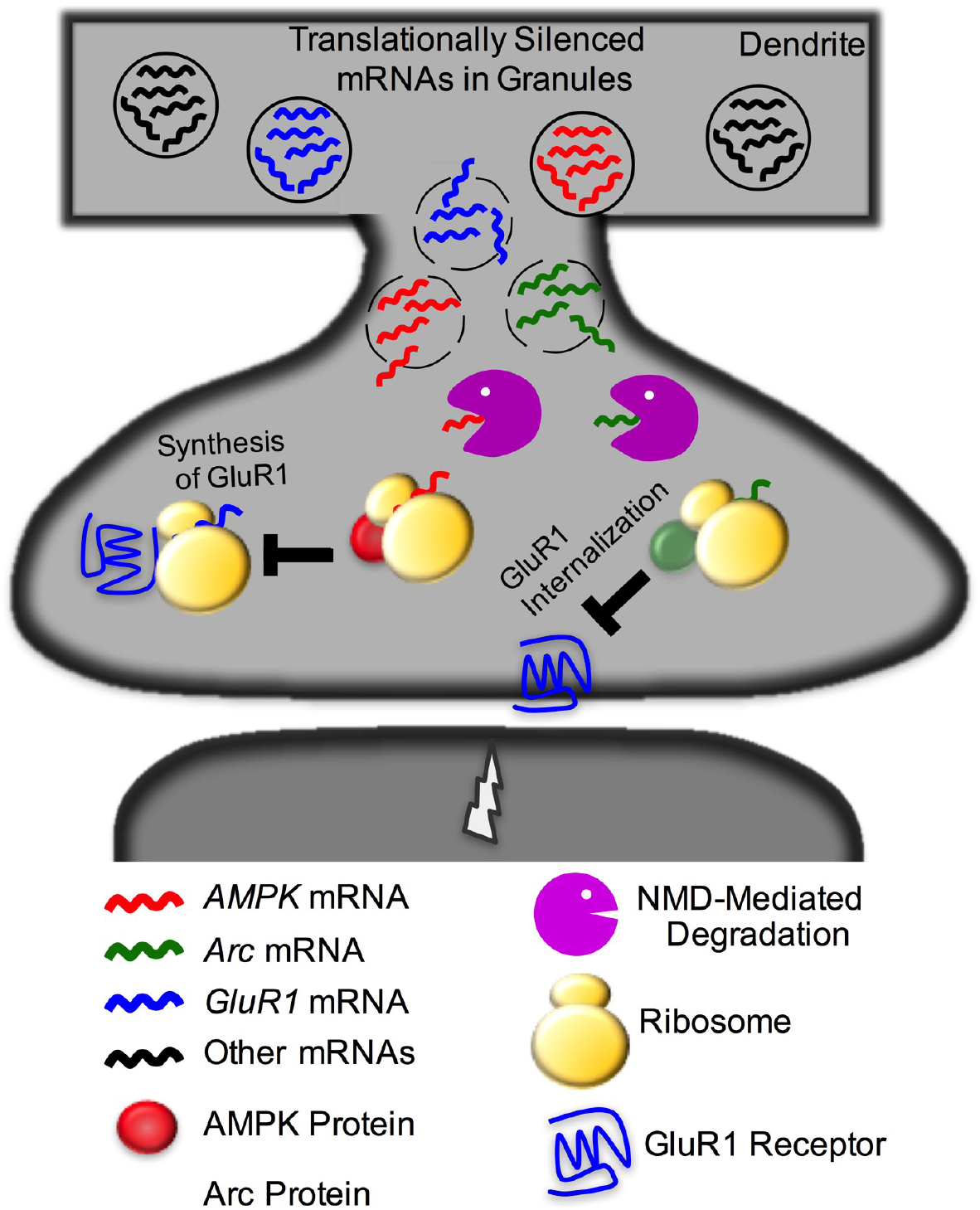
Schematic model for compartmentalized NMD in dendrites. We establish that NMD functions in dendrites to modulate local GluR1 levels, via increased internalization and decreased local synthesis of GluR1 in dendrites. We mechanistically report that NMD degrades both *Arc* and *AMPK* mRNAs within dendrites. When NMD is disrupted, AMPK over-accumulates and suppresses local translation of nascent GluR1. Similarly, Arc also over-accumulates and increases GluR1 internalization rates. Co-normalization of both *Arc* and *AMPK* mRNA levels resulted in the complete recovery of dendritic surface GluR1 levels in UPF2-deficient dendrites. Additionally, our data also suggest that NMD activity is induced by synaptic stimulation. Together, we mechanistically outlay a role for NMD in plasticity, hippocampus-dependent learning and memory, and the local regulation of GluR1 specifically in dendrites, establishing that the NMD pathway is not just a passive surveillance pathway but is a critical, activity-dependent, regulator of synaptic function.

