## Supplemental Figures and legends for "The Nonsense-Mediated mRNA Decay pathway degrades dendritically-targeted mRNAs to regulate long-term potentiation and cognitive function"

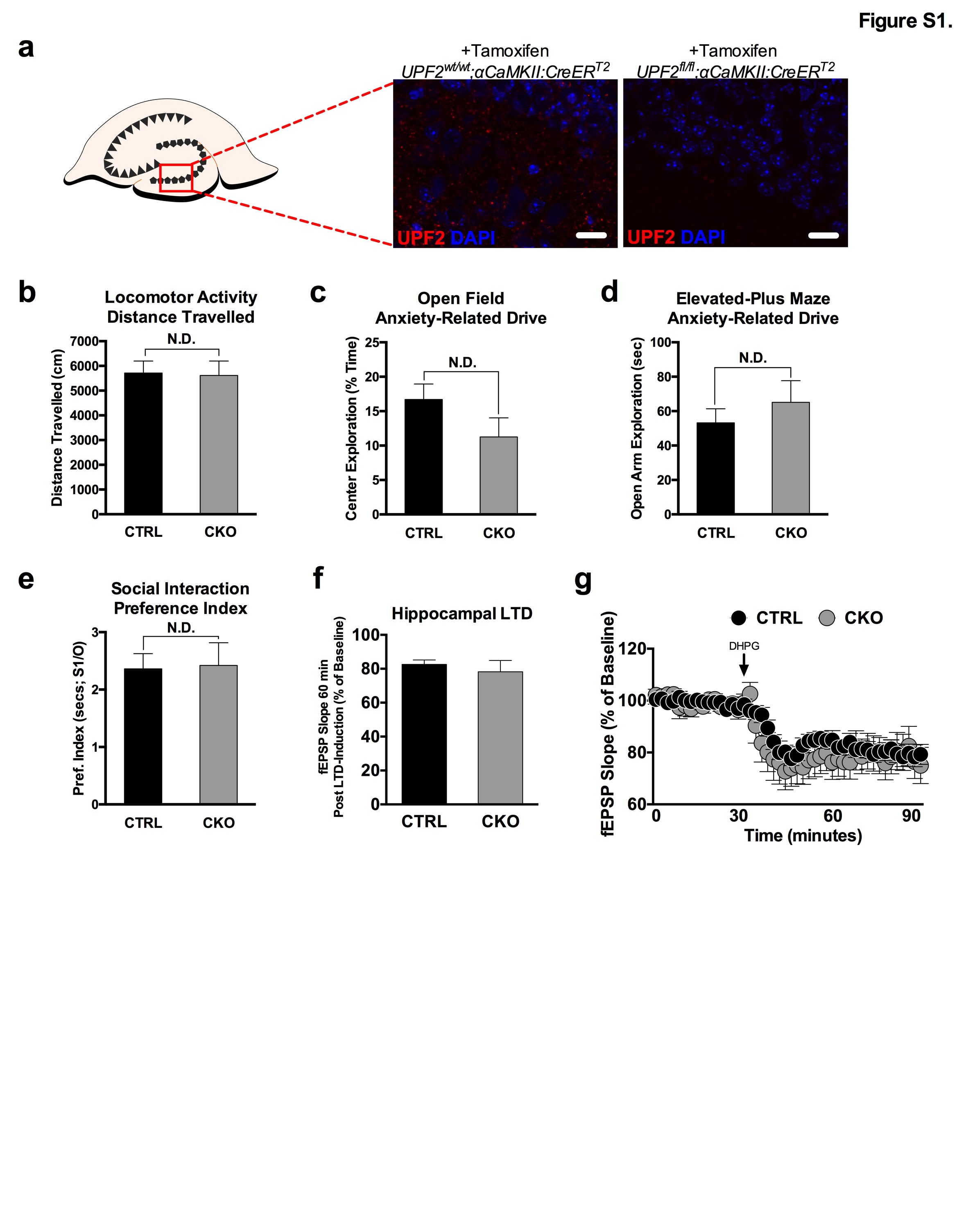

**Figure S1. NMD does not influence baseline locomotor activity, anxiety-related behavior, sociability, and LTD in mice.**

**a**, To determine the potential role of NMD in hippocampus-dependent cognition in the adult brain, we disrupted NMD in postmitotic neurons using a conditional knockout mouse of *UPF2* (*UPF2*^fl/fl^) [1]. To temporally control *UPF2* disruption, we crossed *UPF2^fl/fl^* mice with a mouse line that expresses tamoxifen-inducible Cre under the α*CaMKII* promoter (α*CaMKII*::*CreER^T2^*) [2]. Before assessing synaptic plasticity and learning and memory (Figure 1) in *UPF2*^fl/fl^;α*CaMKII*:*CreER^T2^* (CKO) and *UPF2*^wt/wt^;α*CaMKII*:*CreER^T2^* (CTRL) mice, we confirmed that conditional deletion of *UPF2* gene (using 200 mg/kg Tamoxifen, for 5 doses) led to the successful loss of UPF2 protein in the hippocampus.

**b-d**, To determine whether NMD induces hyperactivity, mice were subjected to a 2 hr locomotor hyperactivity assay in photocells. During this 2 hr trial, the distance that mice travelled in cm was analyzed. Greater distances travelled are used as an indicator of locomotor hyperactivity, which may confound behavioral testing on other maze tasks. Similarly, this assay can also assess hypoactivity induced by genetic alterations, which may also influence task performance.
**b**, In our assessment of baseline locomotor activity, there were no genotype differences between CTRL (n*=*25) and CKO (n=22) mice. This indicates that loss of UPF2 does not result in acquired hyper- or hypoactivity.
**c**, A 10 min open field assay was also conducted in locomotor photocells as an additional control assay, with less time spent exploring the center field of photocells being interpreted as an anxiety-like phenotype. Consistent with the outcome of the locomotor activity assay, no evidence of anxiety-like behavior between CTRL (n=25) and CKO (n=22) mice emerged on this test, indicating that NMD does not modify anxiety-related behavior.
**d**, The elevated-plus maze was used as a control assay to ensure that NMD did not modify anxiety-related exploratory drive, which may compound measurements of learning and memory. Briefly, mice were subjected to a 10 min test session, with time spent exploring the open arms being used as an index of anxiety-like behavior. No genotype differences emerged between CTRL (n*=*25) and CKO (n=22) mice on this test.

**e**, Alterations in both spine density and GluR1 levels are linked to deficits in sociability [3, 4]. In addition, defects in the NMD machinery are linked to numerous neurodevelopmental diseases associated with alterations in normal social behavior [5-10]. To determine whether NMD is required for sociability, we used the three-chamber social interaction task and measured sociability in CTRL and CKO mice. Mice were habituated to the empty three-chamber apparatus for 10 min before being exposed to a stranger mouse placed inside a cup in one chamber and an empty cup in the opposite chamber for an additional 5 min. A preference index was derived from time spent sniffing or interacting with the cup containing the stranger mouse divided by time spent interacting with the empty cup, with higher preference indexes representing more social behavior. No significant differences in preference index emerged between CTRL (n=24) and CKO (n=19) mice on the three-chamber social interaction assay.
Data are represented as mean ± SEM; N.D. represents “No Difference”. Scale bar: 15 μm.

**f-g**, NMD regulates spine density and L-LTP in hippocampus (Figure 1j-m). We also assayed LTD in *UPF2* CKO. Because of age-related decline in LTD, we induced Cre at 3 weeks of age and hippocampal slices were acquired from 6 week-old CTRL and CKO mice (see Methods). LTD was induced by acute application of 100 µM DHPG for 10 min to hippocampal slices. fEPSPs were collected for 60 min post-LTD induction. Compared to CTRL mice (n=14 slices from 7 mice), CKO mice (n=10 slices from 5 mice) did not show significantly different slope values post-DHPG application indicating that LTD is unaltered in the absence of NMD in hippocampus

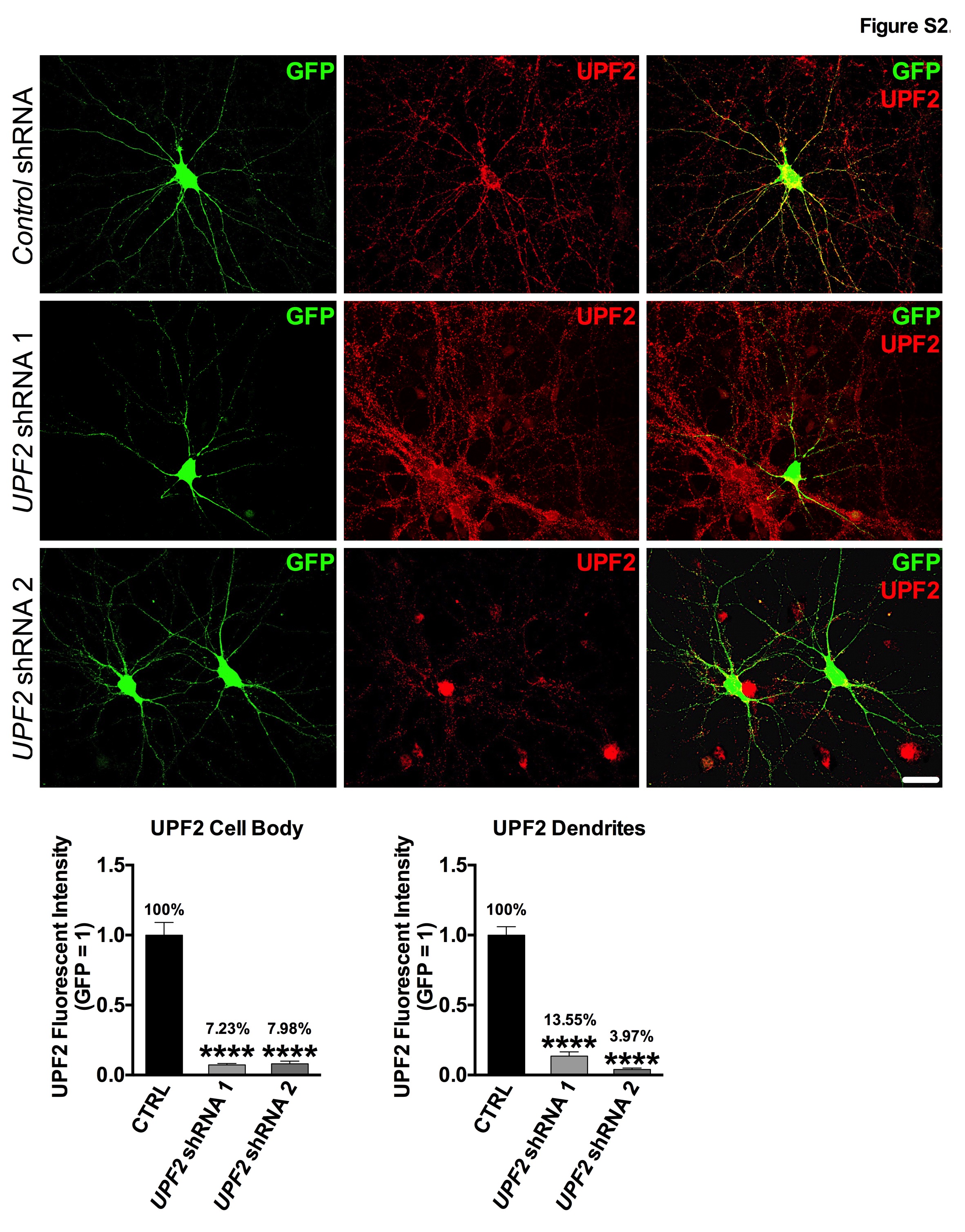

**Figure S2. Knockdown of *UPF2* in mouse hippocampal neurons by *UPF2-*shRNA:GFP lentivirus.**

To disrupt NMD, we targeted the *UPF2* mRNA using *UPF2-*shRNA:GFP lentivirus. Canonical NMD involves the interaction of UPF1, UPF2 and UPF3 proteins. While UPF1 function is not restricted to the NMD pathway [11], UPF2 has been successfully used to disrupt NMD in several studies [1, 12, 13]. We infected E16 hippocampal neurons at DIV7 with *control*- or two independent *UPF2-*shRNA viruses. To assess the knockdown of UPF2, we fixed the cells at DIV14 and performed immunohistochemistry for UPF2. Application of both *UPF2-*shRNA viruses resulted in a robust depletion of UPF2 within infected neurons and their dendrites, while UPF2 expression remained intact in scrambled *control­*-shRNA cultures and non-infected neurons in *UPF2*-shRNA cultures (UPF2+, GFP- cells in *UPF2*-shRNA 2 panel). This control experiment with two separate viruses targeting non-overlapping UPF2 sequences exemplifies specific knockdown of *UPF2* across our studies, and ensures that the phenotypes that arise from this knockdown are unlikely off-target effects. Scale bar: 30 μm.

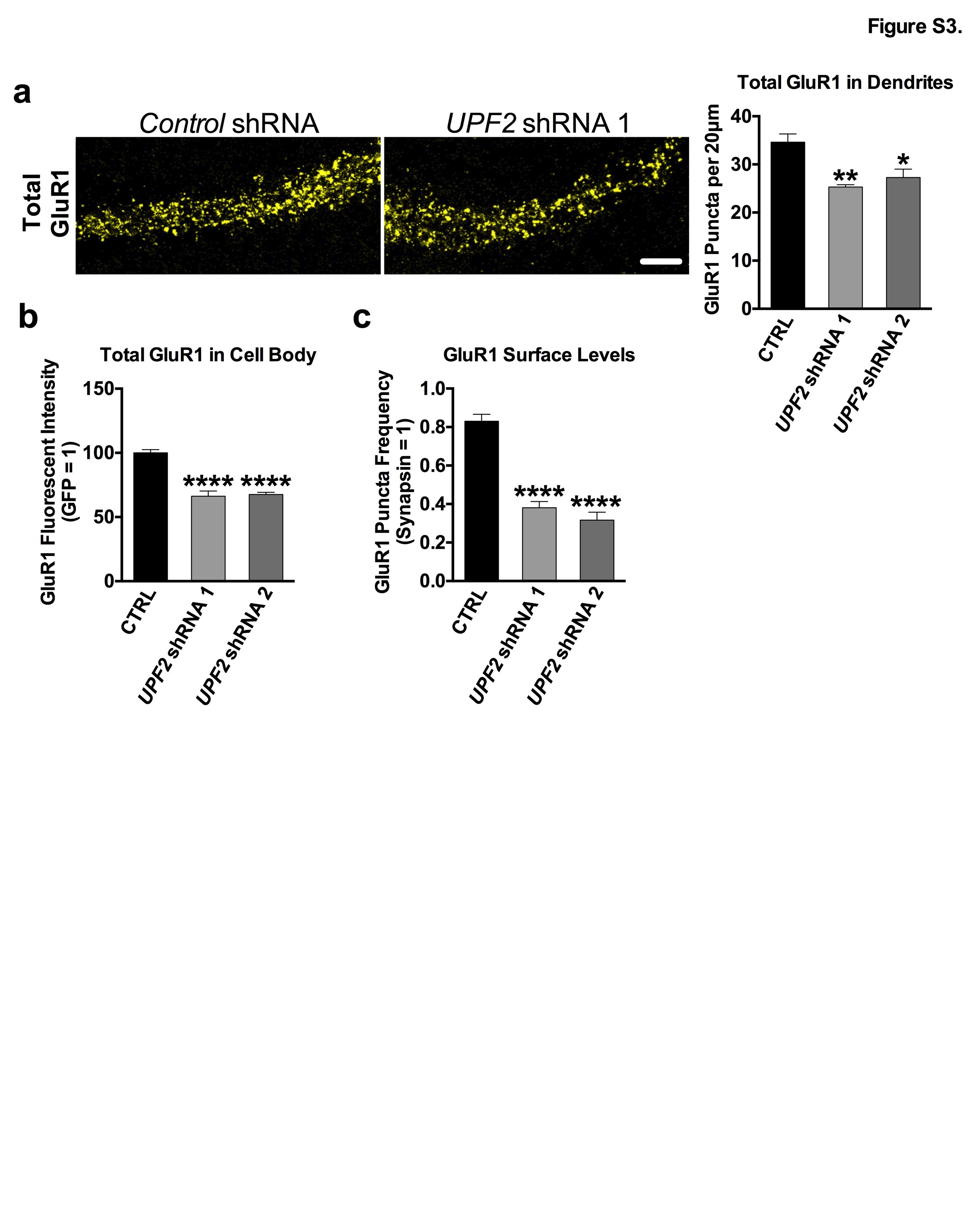

**Figure S3. Total levels and surface levels of GluR1 protein are decreased upon knockdown of *UPF2*.**

Loss of *UPF2* led to a decrease in GluR1 levels *in vivo* (Figure 1m). To explore this phenotype *in vitro,* we infected hippocampal neurons with the 2 independent *UPF2*-shRNA:GFP lentivirus vectors validated in Figure S2 to interrupt endogenous NMD activity. Hippocampal neurons were infected at 7 days *in vitro* (DIV7) and GluR1 expression was examined on DIV21. We quantified total GluR1 signal using an anti-C-terminus-GluR1 antibody in fixed and permeabilized cells.

**a**, UPF2-deficient dendrites had significantly lower GluR1 density compared to control dendrites (n=3 biological replicates per *control*-shRNA [10 neurons, 23 dendrites] and per *UPF2*-shRNA [11 neurons, 25 dendrites]; representative image reflects *UPF2* shRNA 1, and *UPF2* shRNA refers to the UPF2 shRNA 1 in the following text). This suggests that while NMD does not alter *GluR1* transcription or mRNA degradation (Figure 2d), there is either an increase in GluR1 receptor degradation or a decrease in GluR1 synthesis.

**b,** Total GluR1 levels were also significantly decreased in the cell bodies of UPF2-deficient neurons. We measured total fluorescence signal by measuring the integrated density in a defined region using ImageJ. Knockdown of *UPF2* led to a decrease in total GluR1 levels in cell bodies (n=3 biological replicates per *control*-shRNA [15 neurons] and per *UPF2*-shRNA [13 neurons]).

**c,** Surface GluR1 expression was also quantified, as described both in-text and methods, using both validated *UPF2*-shRNA:GFP lentivirus vectors. Similar to total GluR1 levels in dendrites, surface levels of GluR1 were also decreased in UPF2-deficient dendrites (n=3 biological replicates per *control*-shRNA [15 neurons] and per *UPF2*-shRNA [13 neurons]). Increased local internalization and decreased local synthesis of GluR1 are likely to contribute to this phenotype (see Figure 3).

Data are represented as mean ± SEM; **p < 0.01, and ***p < 0.001. Scale bar: 20 μm.

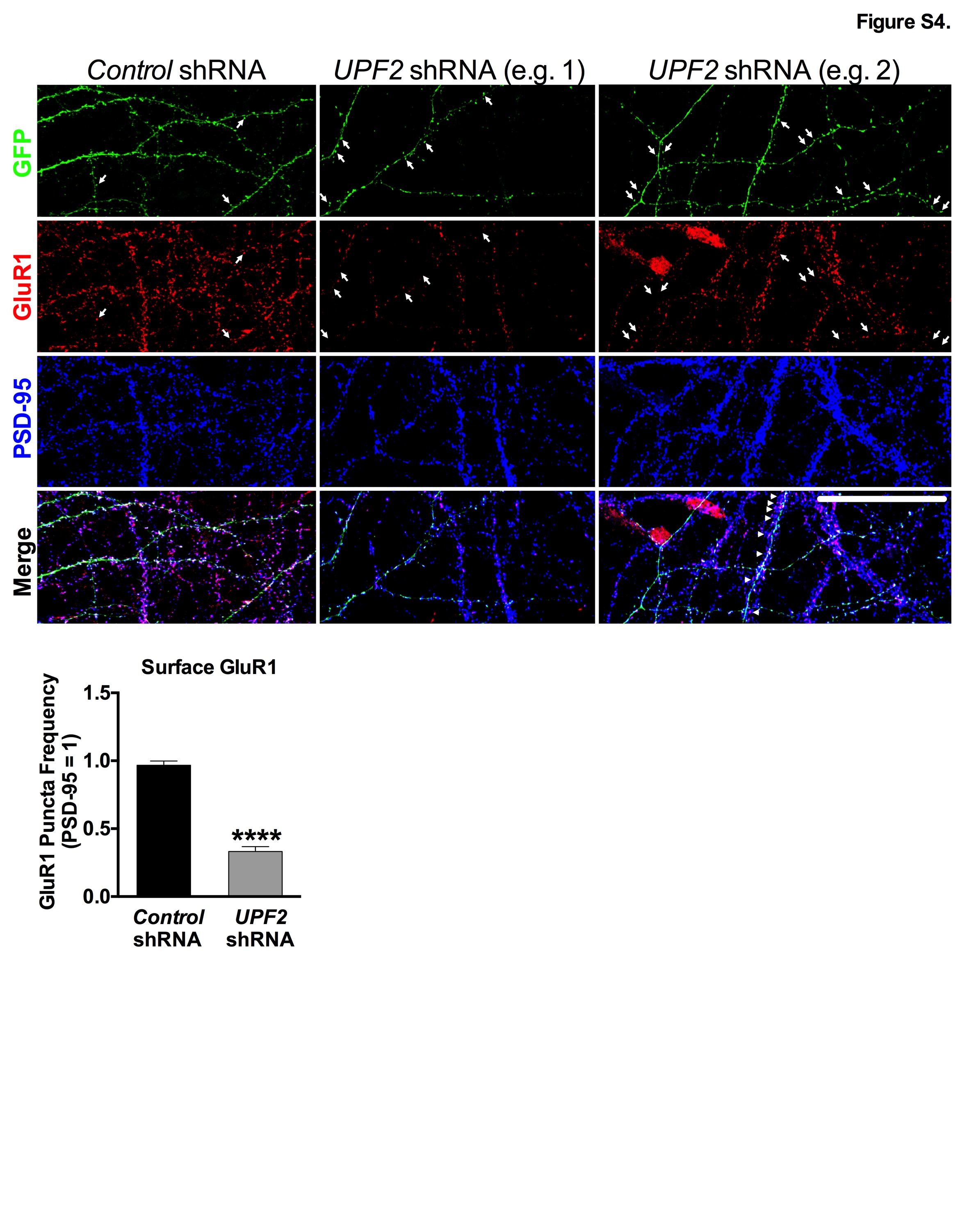

**Figure S4. Decrease in GluR1 surface levels in UPF2-deficient dendrites was also recapitulated when the synaptic marker PSD-95, in addition to synapsin, was used for quantifications.**

In Figure 2, we show that knockdown of *UPF2* in hippocampal neurons led to a robust depletion of surface GluR1 levels without altering the mRNA transcription of this receptor. For these quantifications, we used synapsin as a normalization marker as this is consistent with prior work (see Methods). To address whether this GluR1 phenotype is recapitulated when another synaptic marker is used for normalization, we repeated this experiment using the postsynaptic marker PSD-95. Similar to the synapsin quantifications, the disruption of UPF2 in hippocampal neurons resulted in a significant depletion of surface GluR1 levels when synapses were identified and normalized against PSD-95 expression (n=3 biological replicates per group; 11 dendrites from 7 neurons for *control*-shRNA cultures, and 12 dendrites from 8 neurons for *UPF2-*shRNA cultures). Arrows show examples of GFP and GluR1 colocalization in spines. Two panels of *UPF2-*shRNA dendrites are presented to exemplify contrasting aspects of GluR1 expression. Arrows depict the loss of surface GluR1 in spines of *UPF2*-shRNA infected hippocampal neurons relative to control cultures. Arrowheads in the merged panel of *UPF2-*shRNA example 2 depict a non-infected dendrite segment with intact GluR1 expression that is overlapping an infected dendrite that exhibits reduced GluR1 colocalization in isolated GFP+ spines. These data replicate the depletion of GluR1 surface levels in UPF2-deficient dendrites presented in Figure 2, and provide confirmation that synapsin produces concordant data to PSD-95 when used to evaluate surface GluR1 levels.

Data are represented as mean ± Standard Error of the Mean (SEM); ****p < 0.0001. Scale bar: 30 μm.

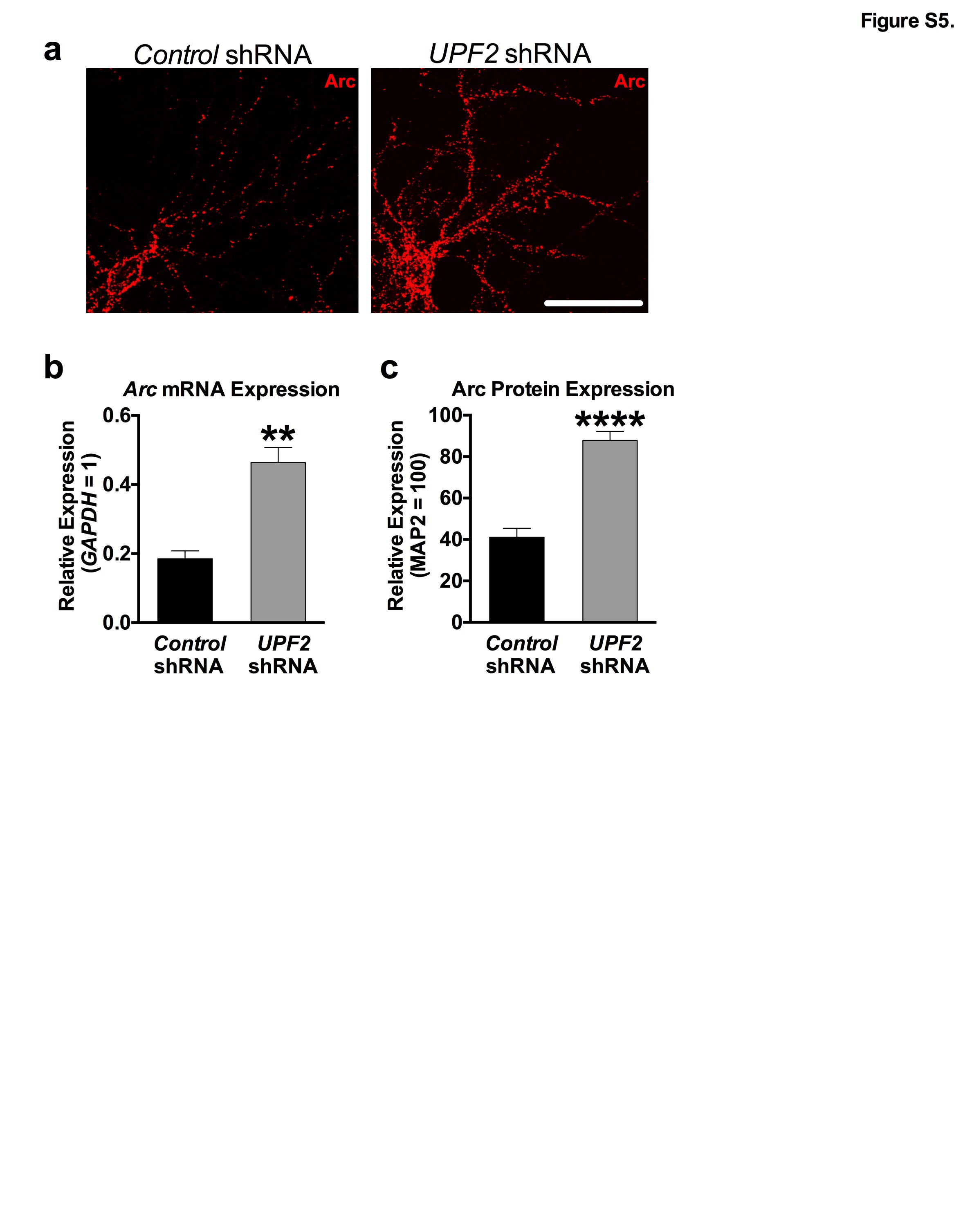

**Figure S5. Arc levels are increased upon knockdown of *UPF2* in hippocampal neurons.**

**a**, Arc immunostaining in mouse hippocampal neurons upon knockdown of *UPF2*. Disruption of NMD caused a significant reduction in the total (Figures 1 and S3) and surface (Figure 2) levels of GluR1 without affecting the *GluR1* mRNA levels (Figure 2d). This suggests that NMD regulates GluR1 surface expression through other mechanisms than transcription or *GluR1* degradation. GluR1 signaling is modulated by dynamic insertion and removal of the GluR1 receptor to the synaptic surface [14]. The synaptic plasticity protein Arc drives the removal of GluR1 from the surface of synapses. *Arc* mRNA is a known target of NMD. Both *Arc* mRNA and Arc protein are increased upon disruption of NMD [12, 15] suggesting that elevated Arc levels may contribute to the reduced surface expression of GluR1. To confirm that knockdown of *UPF2* alters endogenous Arc levels, we cultured E16 mouse hippocampal neurons and infected with *control*- or *UPF2-*shRNA lentivirus at DIV7 to disrupt NMD. At DIV21, we performed immunostaining for Arc protein.

**b**, Quantitative RT-PCR analysis of *Arc* mRNA in UPF2-deficient neurons. To quantify *Arc* mRNA, we performed qRT-PCR. *GAPDH* was used as a control transcript. Expression of *Arc* was presented relative to *GAPDH*. Neurons infected with *UPF2*-shRNA lentivirus displayed exaggerated *Arc* mRNA expression (n=3 biological replicates per group) compared to control neurons.

**c**, Quantifications of Arc protein in UPF2-deficient neurons. Quantifications of fluorescent intensity, normalized to MAP2, showed that Arc protein levels are also increased in neurons with disrupted UPF2 (per group, n=10 dendrites for analysis of fluorescent intensity). Together, these data confirm that Arc levels are increased upon disruption of UPF2. Arc is known to induce internalization of GluR1 receptor [16]. However, the total levels of GluR1 are independent of its internalization rate. While Arc-mediated internalization of GluR1 influences surface levels, it does not influence total GluR1 levels [17]. This suggests that while loss of UPF2 does not alter *GluR1* transcription or mRNA degradation (Figure 2d), there is either an increase in GluR1 receptor degradation or a decrease in GluR1 synthesis.

Data are represented as mean ± Standard Error of the Mean (SEM); ** p < 0.01, **** p < 0.0001. Scale bar: 30 μm.

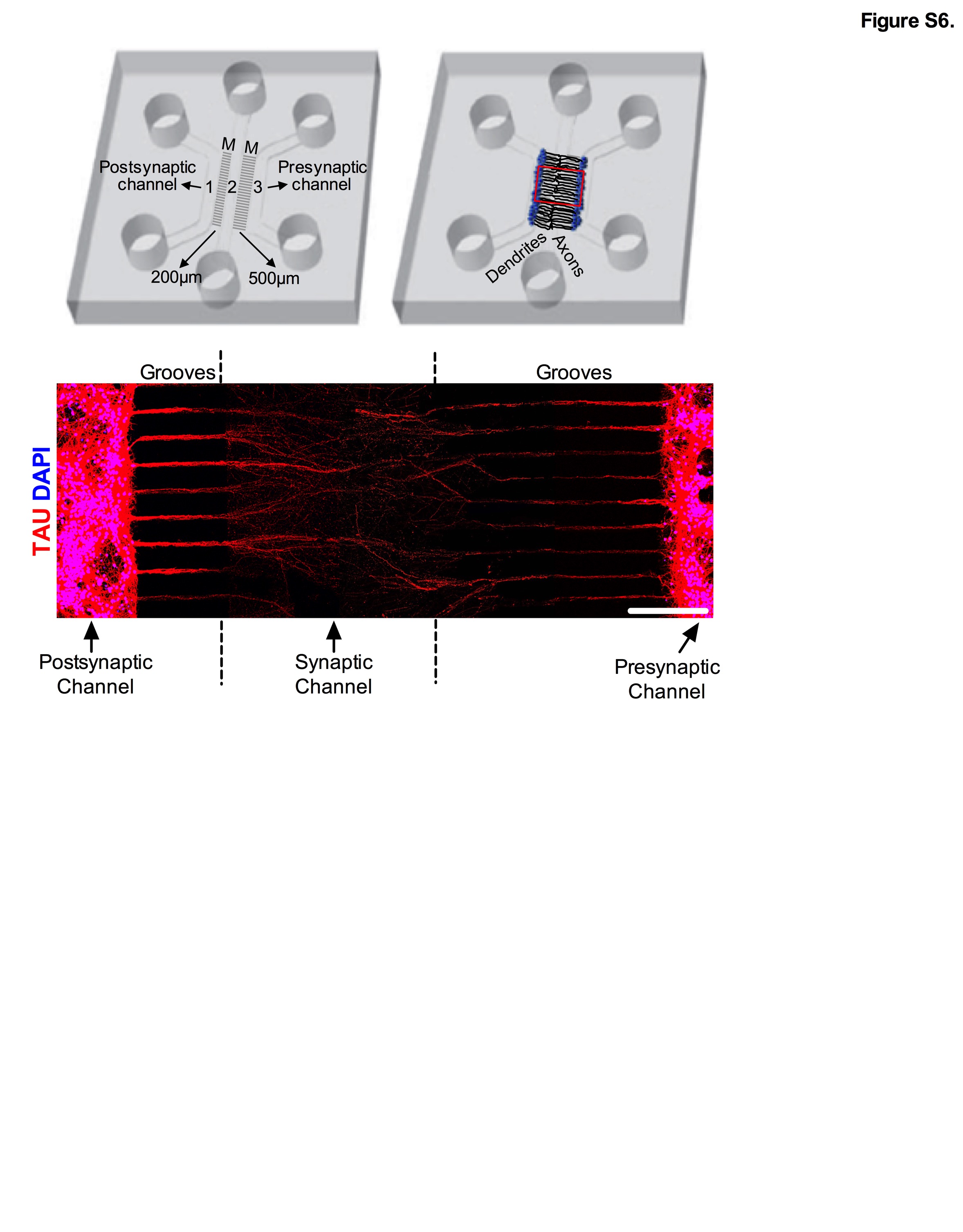

**Figure S6. Axonal penetrance of microgrooves in tripartite microfluidic devices.**

As shown in the schematic presented here, and the MAP2 immunostaining presented in Figure 3, our custom-engineered tripartite microfluidic device only permits the dendrites of neurons from left channel (channel 1) to penetrate the middle channel, which is referred to as the “synaptic channel” (channel 2). Here, TAU staining shows that axons from both channels extend to the synaptic channel suggesting that axons of both channel 1 and channel 3 can form synapses with dendrites that originated from channel 1. However, due to their limited length, dendrites remain in very close proximity to channel 1 (Figure 3). This tends to restrict dendrite-axon synapses arising from neurons in channel 1 to the proximal-most area of the synaptic channel. Because axons are very long, and the short microgrooves that separate channels 1 and 2 are just 200 μm in length, most channel 1 axons span the entire synaptic channel and go-on to penetrate channel 3. Thus, the majority of the synapses formed in the synaptic channel consist of dendrites projecting from channel 1 and axons originating from channel 3, referred to as the “presynaptic channel”. UPF2 is not expressed in mature axons (Figure S10), and only dendrites from neurons plated in channel 1 can reach the synaptic channel (Figure 3). Therefore, regardless of where the presynaptic axons project from, or where synapses are formed within the synaptic channel, the manipulation of NMD exclusively in channel 1 is confined to postsynaptic dendrites. Scale bar: 200 μm.

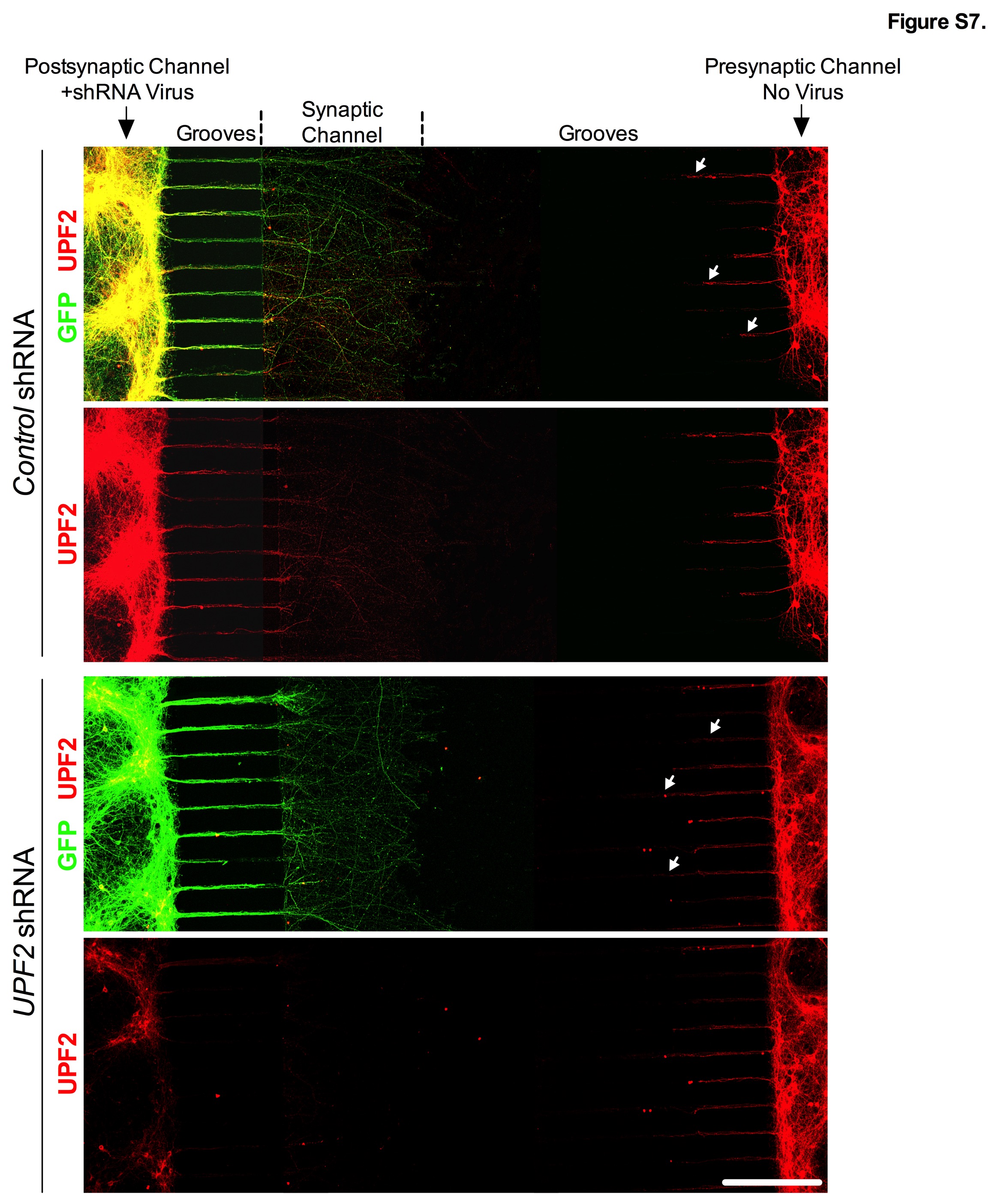

**Figure S7. Application of *UPF2* shRNA virus exclusively to the postsynaptic channel results in loss of UPF2 in this channel, as well as in dendrites within the synaptic channel, but not in presynaptic channel neurons or their projections.**

The physical and fluidic isolation of channels is a fundamental feature and component of microfluidic devices, as it permits the selective manipulation/isolation of neuronal populations and compartments. To address the role of NMD in dendrites in synaptic function, we selectively treated postsynaptic channels with *UPF2*-shRNA virus at DIV7. To determine whether the virus application resulted in selective loss of UPF2, we performed immunostainings for UPF2 following fixation of cells at DIV13. Consistent with the fluidic isolation of channels, the GFP signal was specific to infected neurons restricted to the postsynaptic channels and their neurites in the synaptic channel. Selective application of the *UPF2*-shRNA virus in the postsynaptic channel resulted in nearly complete loss of UPF2 protein in this channel, as well as in dendrites in the synaptic channel, but not in the presynaptic channel. Because mature axons do not express UPF2 (Figure S10), the UPF2-positive signal in the synaptic channels in the control experiment solely reflects dendritic expression. Consistent with this, dendrites that emerged from the presynaptic channel, which only traveled half way through microgrooves (white arrows), also retained their UPF2 expression in both the *control*- and *UPF2*-shRNA experiments (see Figure 3 and Figure S6 for the comparison of dendrite and axon projection in these grooves). In all experiments that employed *control-* or *UPF2*-shRNAs in tripartite devices, viruses were only applied to the postsynaptic cell body channel of microfluidic devices. Scale bar: 200 μm.

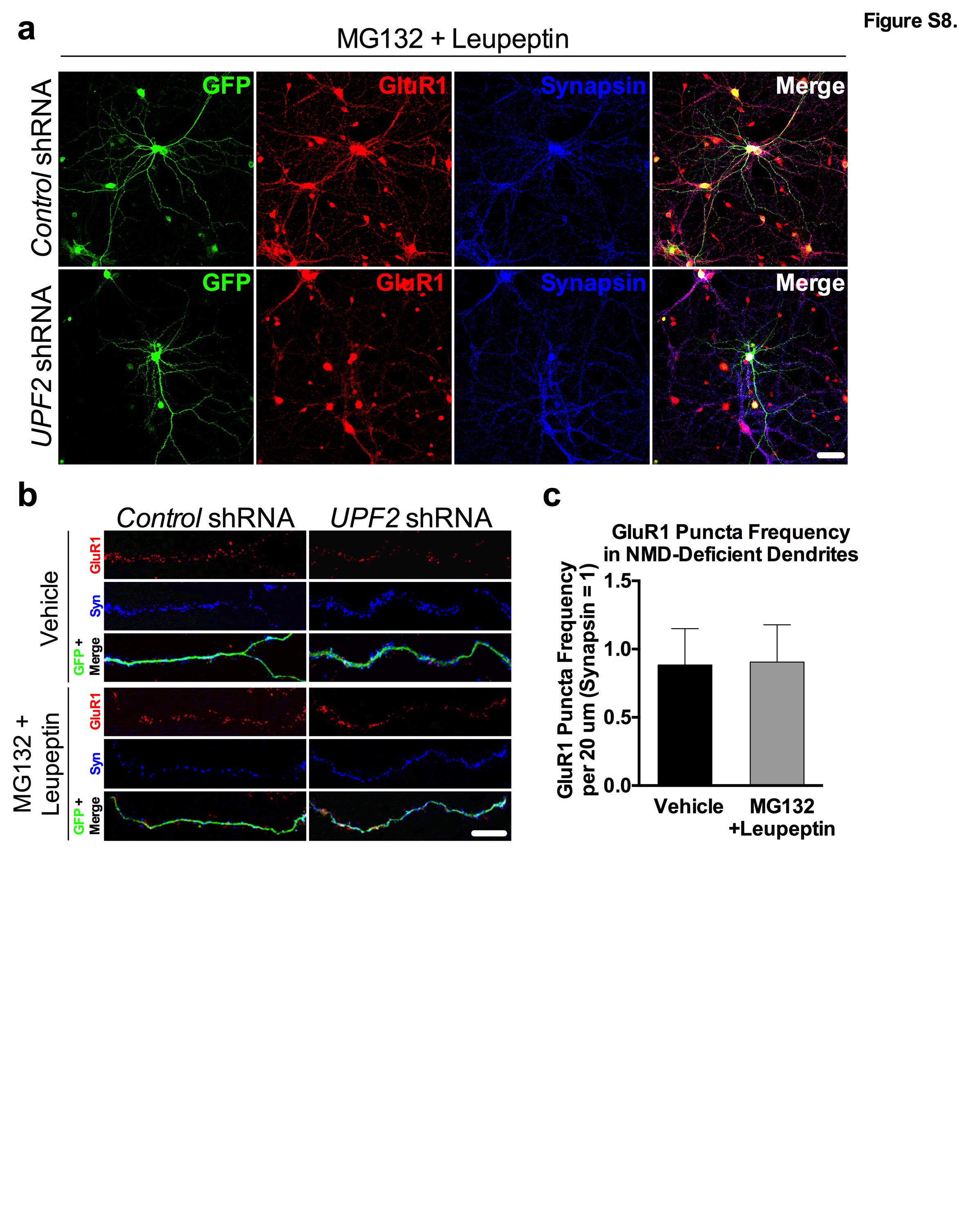

**Figure S8. GluR1 degradation is not altered upon disruption of UPF2.**

Both the surface and total expression of GluR1 in dendrites is reduced upon knockdown of UPF2 (Figure S3 and 2). Although the increase in basal internalization rate of GluR1 (Figure 3a) contributes to the reduction in surface levels of this receptor in the absence of UPF2, it does not explain the decrease in total GluR1 levels [17]. Since the disruption of NMD does not alter *GluR1* transcription or degradation (Figure 2d), an alteration in the balance of local synthesis and degradation of GluR1 is likely to account for the overall reduction in the total levels of this receptor upon disruption of NMD. Therefore, we examined the local synthesis of nascent GluR1 protein (Figure 3) as well as the degradation of GluR1 upon loss of UPF2. Figure 3 shows that the local synthesis of GluR1 is repressed in UPF2-deficient dendrites. To study GluR1 protein degradation, we cultured E16 mouse hippocampal neurons in either 24 well plates or in the cell body compartments of tripartite chambers and infected with *UPF2-*shRNA lentivirus at DIV7. In the case of tripartite chambers, we only infected postsynaptic cells with the *UPF2-*shRNA lentivirus. Because endocytosed AMPA receptors can undergo either lysosome- or proteasome-mediated degradation [18-22], we targeted both lysosomal and proteasomal degradation simultaneously. At DIV21, proteasomal and lysosomal degradation were inhibited with 10 μM of the protease inhibitor MG132 and 20 μM of leupeptin, respectively, for 6 hr. For the tripartite experiment, proteasomal and lysosomal degradation were inhibited by treating the synaptic channels with 10 μM of the protease inhibitor MG132 and 20 μM of leupeptin, respectively, for 6 hr. To label surface GluR1, neurons were fixed and stained with an anti-Nterminus-GluR1 antibody without permeabilization (see also Methods).
**a-b**, Representative low (regular cultures) and high magnification (in tripartite chambers) images of GluR1 surface expression in treated and non-treated UPF2-deficient dendrites.
**c,** Quantification of surface GluR1 puncta shows that inhibition of proteasomal and lysosomal degradation did not significantly influence GluR1 puncta frequency in UPF2-deficient dendrites (n=3 biological replicates per group; 11 neurons per group).

Data are represented as mean ± SEM. Scale bar: 20 μm.

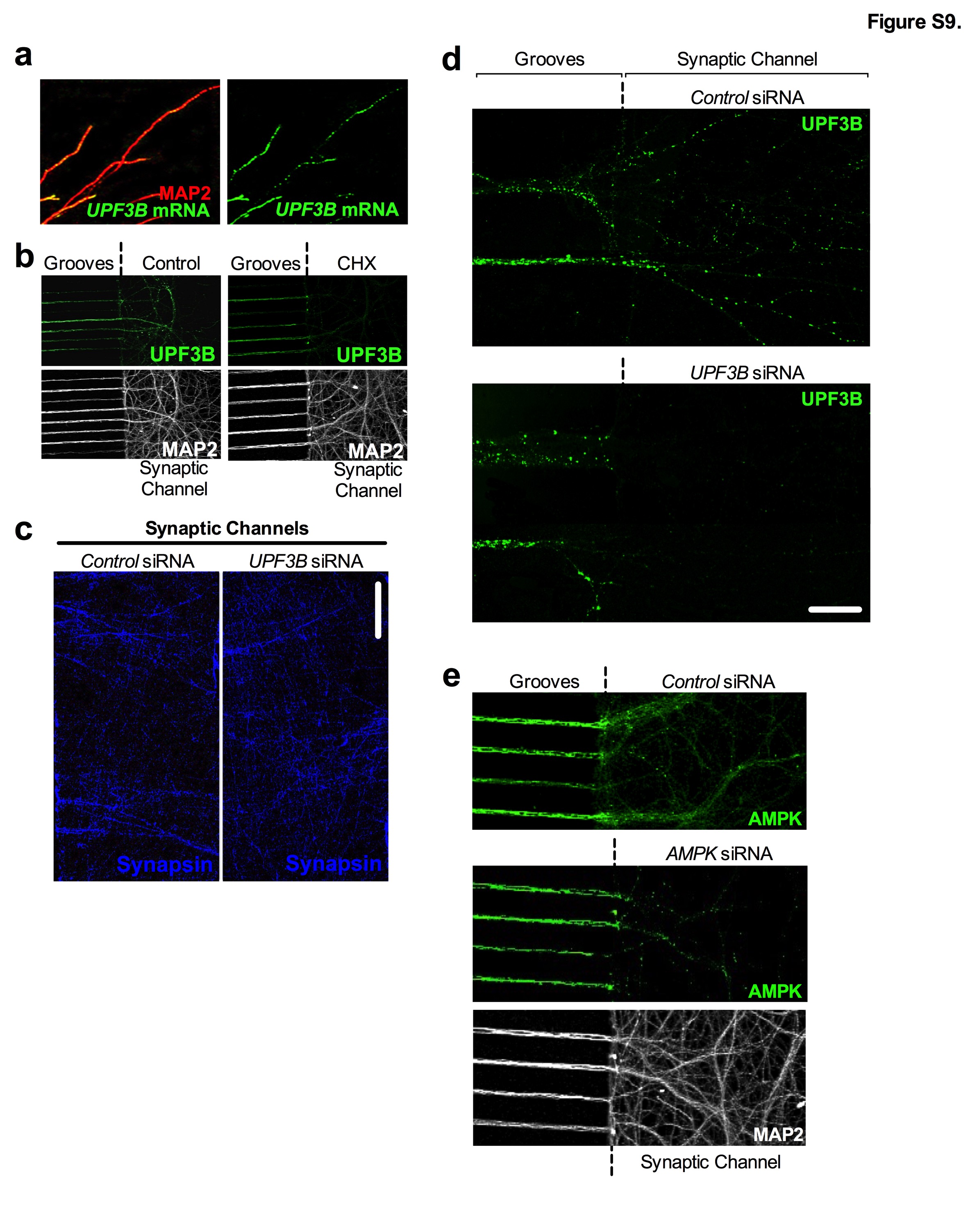

**Figure S9. UPF3B is locally synthesized in dendrites.**

**a-b**, UPF3B is locally synthesized in hippocampal dendrites. Both dendritic internalization and local synthesis of GluR1 receptor are altered upon knockdown of UPF2. In addition, NMD targets *Arc* and *AMPK* are subjected to translation-dependent degradation in dendrites (Figure 4). Although these data suggest that NMD might regulate GluR1 locally in dendrites, there is no direct evidence for the local requirement for NMD in these compartments. To study the specific requirement of locally-occurring NMD in the regulation of GluR1 within dendrites, we devised a strategy for targeting UPF3B protein (which, like UPF2, is also a specific and major component of the NMD machinery) locally within the synaptic channel of tripartite devices (Figure 4d-e). Dendritic transcriptome analysis showed that the mRNA of UPF3B protein is localized to dendrites [23]. To confirm this, we performed Fluorescent In Situ Hybridization (FISH) on hippocampal neurons with antisense riboprobes against *UP3B* mRNA at DIV21. *UPF3B* FISH resulted in punctate labeling (green) along dendrites confirming the localization of *UPF3B* mRNA to these compartments (**a**). Next, we sought to determine if UPF3B is locally synthesized in dendrites. To do this, we inhibited *UPF3B* mRNA translation via dendritic application of Cycloheximide (CHX, 10 μM) and evaluated the UPF3B protein in synaptic channels. We cultured E16 mouse hippocampal neurons in tripartite chambers and, at DIV21, we treated the synaptic channels with CHX for 6 hr. This resulted in an almost complete loss of UPF3B protein in dendrites (**b**). MAP2 was used to visualize dendrites in both (**a**) and (**b**).

**c-d**, To determine whether inhibition of local synthesis of UPF3B in synaptic channels alters GluR1 locally, we selectively treated synaptic channels with siRNAs against the *UPF3B* mRNA and evaluated GluR1 levels (Figure 4d-e). Here, we confirmed that while the treatment of synaptic channels with *UPF3B* siRNA-cocktail (10 nM) for 7 days did not cause an alteration in synaptic potential in these channels (**c**), it led to a complete loss of UPF3B protein (**d**). Note that the expression of UPF3B remained intact within dendrites adjacently located within fluidically-isolated microgrooves in panel **d**.

**e**, Activated AMPK is known to interrupt translation by either inhibiting the mammalian target of rapamycin (mTOR) kinase or the translation elongation factor 2. These data suggest that NMD might regulate total levels of GluR1 through degradation of *AMPK* mRNA in dendrites. To determine whether local AMPK in dendrites modulates GluR1 levels, we inhibited AMPK synthesis in the synaptic channels of tripartite microfluidic devices and examined total GluR1 levels (Figure 4**f**). To do this, we targeted *AMPK* mRNA by systematic application of siRNAs against the *AMPK* mRNA to synaptic channels. Application of 10 nM of two non-overlapping *AMPK-*siRNAs to synaptic channels for 7 days, starting at DIV14, led to selective depletion of AMPK in dendrites in synaptic channels but not in dendrites in microgrooves (see middle panel).

Data are represented as mean ± SEM. Scale bar: **c** 20 μm, **d** 75 μm.

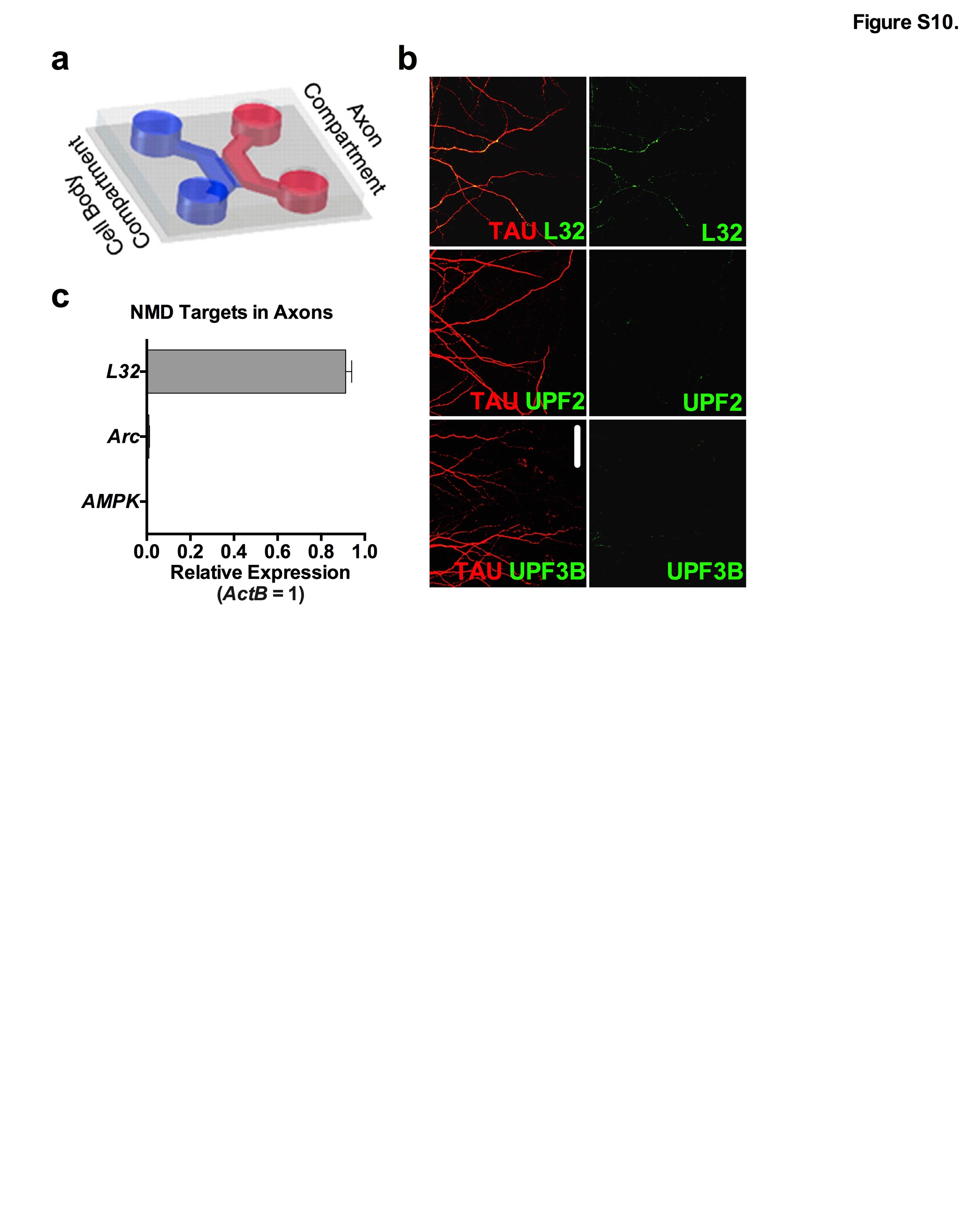

**Figure S10. The NMD machinery and NMD targets are not expressed in mature axons.**

**a-c**, The synaptic channel of tripartite chambers contains both dendrites and axons, raising the possibility that some of the phenotypes observed upon loss of local UPF3B or AMPK in synaptic channels might arise from affected axons. To clarify this, we addressed whether the NMD machinery, as well as *Arc* and *AMPK,* are also present in mature axons in addition to dendrites. We cultured E16 neurons in two-partite microfluidic chambers (**a**). Unlike tripartite chambers used in other experiments, two-partite chambers allow physical isolation of axons [12]. At DIV21, we collected the axonal material and measured the levels of *Arc* and *AMPK* mRNAs by qRT-PCR (**c**). While the axonal mRNA *L32* was readily detected, mature hippocampal axons lacked *Arc* and *AMPK* mRNAs (n=3 biological replicates). This is consistent with previous reports that the axonal transcriptome significantly shrinks as axons mature [24] and does not include any known NMD targets [25, 26]. Similarly, unlike navigating axons [12], mature axons did not contain the major proteins of the NMD machinery (**b**). These data suggest that the GluR1 phenotypes observed upon application of siRNAs against both *UPF3B* and *AMPK* in the synaptic channels of tripartite chambers solely originate from dendrites and not axons. Data are represented as mean ± SEM; Scale bar: 30 μm.

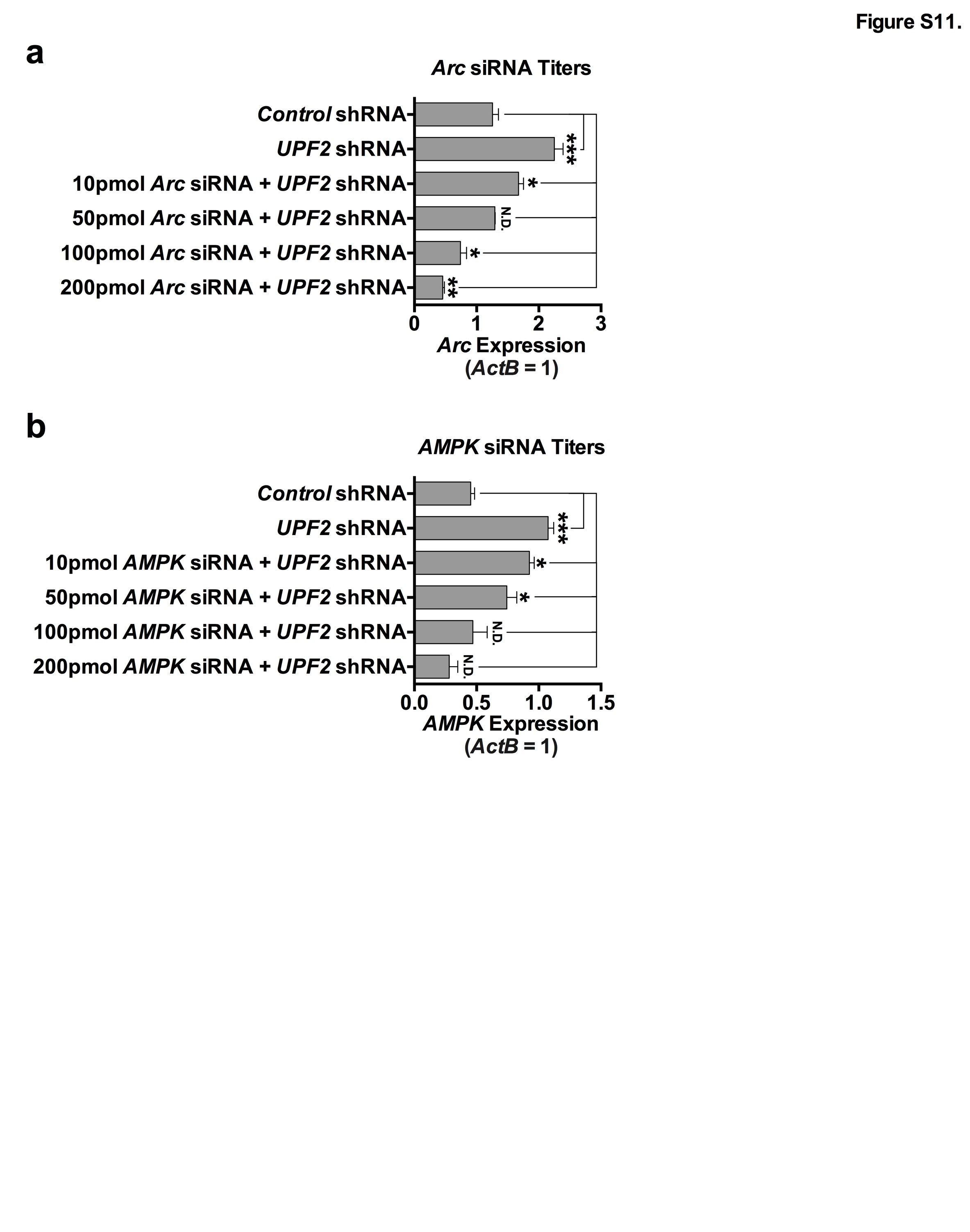

**Figure S11. *Arc* and *AMPK* titers for siRNA normalization of exaggerated expression in NMD-deficient dendrites.**

**a-b**, We have identified *Arc* and *AMPK* as candidate targets in the NMD-mediated regulation of GluR1 levels within the dendrites and synaptic compartments of hippocampal neurons. We next sought to functionally determine which of these molecules is responsible for depletion of surface GluR1 in UPF2-deficient dendrites. To do this, we systematically modulated the elevated levels of *Arc* and *AMPK* in dendrites by siRNA transfection of synaptic channels that contain UPF2-deficient dendrites (Figure 5). As siRNA treatment affects mRNA expression in a concentration-dependent manner [27-30], we first determined the amount of siRNA required to correct the levels of each target in UPF2-deficient dendrites presented in Figure 5. First, we applied the *UPF2*-shRNA virus to the postsynaptic-cell channels of tripartite microfluidic devices at DIV7. Starting at DIV14, we treated the synaptic channels of UPF2-deficient dendrites with non-overlapping siRNAs (2 independent siRNAs per mRNA; see Methods) at different concentrations for either *Arc* (**a**) or *AMPK* (**b**). To deliver siRNAs, we used 10% NeuroPORTER (Sigma). At the end of the treatment, synaptic channels were perfused with TRIzol and lysates used for quantitation of *Arc* and *AMPK* mRNA by qRT-PCR. A panel of *Arc* or *AMPK* siRNA-cocktail titers was selectively applied to the synaptic channels of tripartite chambers (n=3 replicates per mRNA and group). As expected, infection of neurons with *UPF2*-shRNA virus significantly increased the expression of *Arc* (**a**) and *AMPK* (**b**).
**a**, Subsequent treatment with *Arc* siRNA cocktail at 10 pmol led to a significant decrease in the expression of *Arc* mRNA in the dendrites of UPF2-deficient postsynaptic neurons, but was still elevated relative to levels observed in the dendrites of postsynaptic neurons in control cultures. Similarly, 100 pmol and 200 pmol titers of *Arc* siRNA cocktail led to a significant downregulation of *Arc* mRNA in the dendrites of UPF2-deficient postsynaptic neurons, but produced expression levels lower than that required to recapitulate the physiological profile of dendritic *Arc* mRNA expression observed in control cultures. However, we observed that 50 pmol of *Arc* siRNA cocktail normalized the expression of *Arc* mRNA in the dendrites of UPF2-deficient postsynaptic neurons, and that these cultures were indistinguishable from control cultures, leading us to adapt this titer in experiments Figure 5.

**b**, Treatment of the synaptic channels of tripartite chambers with 10 pmol and 50 pmol titers of *AMPK* siRNA cocktail significantly reduced the overexpression of *AMPK* in the dendrites of UPF2-deficient postsynaptic neurons, but did not restore expression levels to those observed in control cultures. 100 pmol *AMPK* siRNA cocktail titer was chosen for rescue experiments in Figure 5, as raw values closely resembled and statistically did not differ from those observed in the control group (n=3 replicates).

Data are represented as mean ± SEM; * p < 0.05, ** p < 0.01, *** p < 0.001, N.D. indicates “No Difference”.

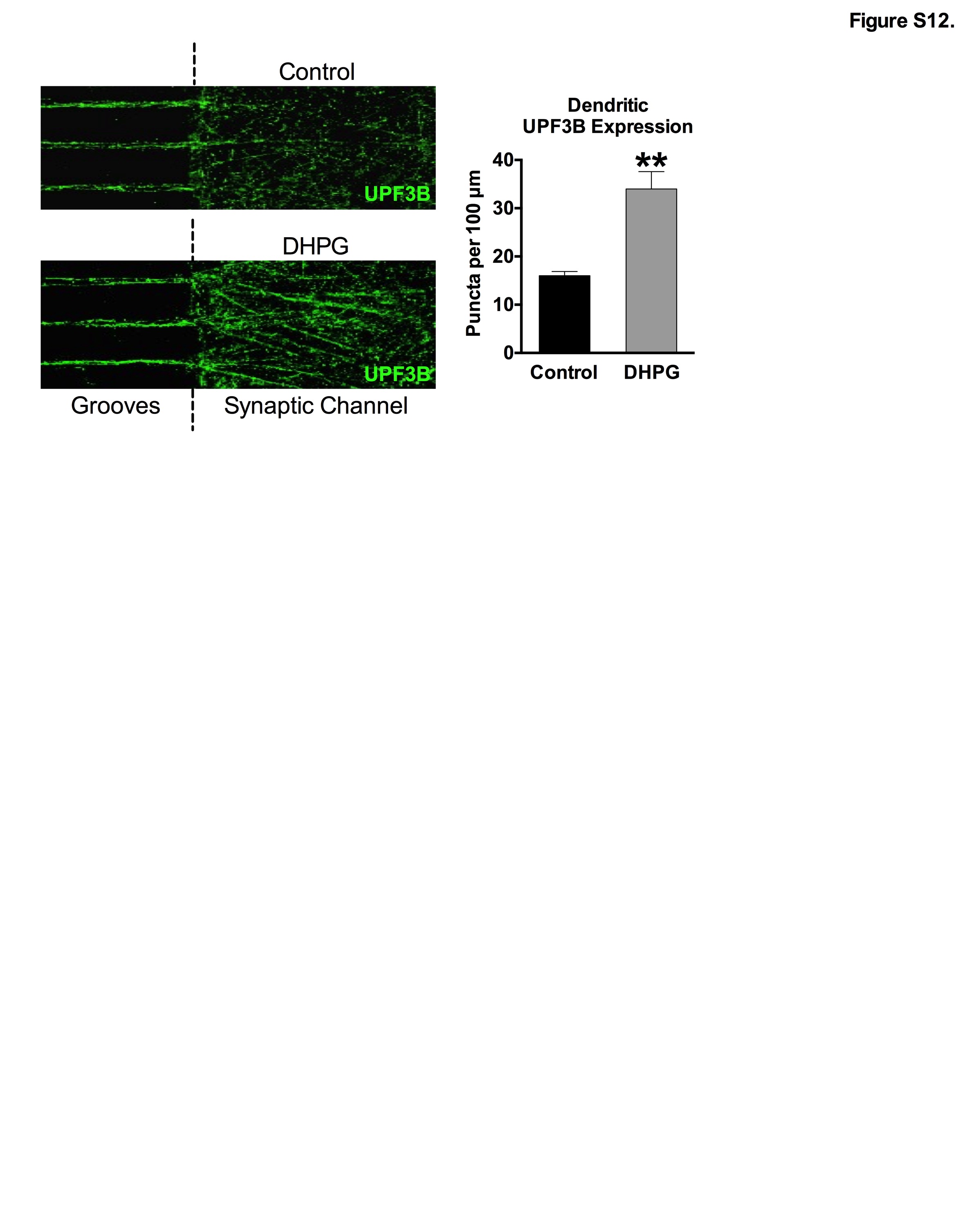

**Figure S12. Neuronal activity upregulates local UPF3B expression in dendrites.**

The mRNA of UPF3 protein is trafficked to [23] and locally translated in dendrites (see Figure S9). Local translation of selected mRNAs is induced by activity in dendrites [31], suggesting that NMD might be regulated at the level of translation in these compartments. To test this, we stimulated synapses with the mGluR agonist DHPG and measured the levels of UF3B in dendrites. E16 mouse hippocampal neurons were cultured in tripartite microfluidic devices and, at DIV21, the mGluR agonist DHPG (100 μM) was selectively applied to synaptic channels for 5 min to induce synaptic activity. Following fixation of cells, immunostaining for UPF3B was performed in control (vehicle) and DHPG-treated dendrites. Quantification of UPF3B puncta within 100 μm dendrite segments revealed a significant increase in UPF3B expression following treatment with DHPG relative to vehicle-treated cultures (n=3 biological replicates per group). This suggests that NMD is induced by neuronal activity in dendrites.
Data are represented as mean ± SEM; ** p < 0.01.

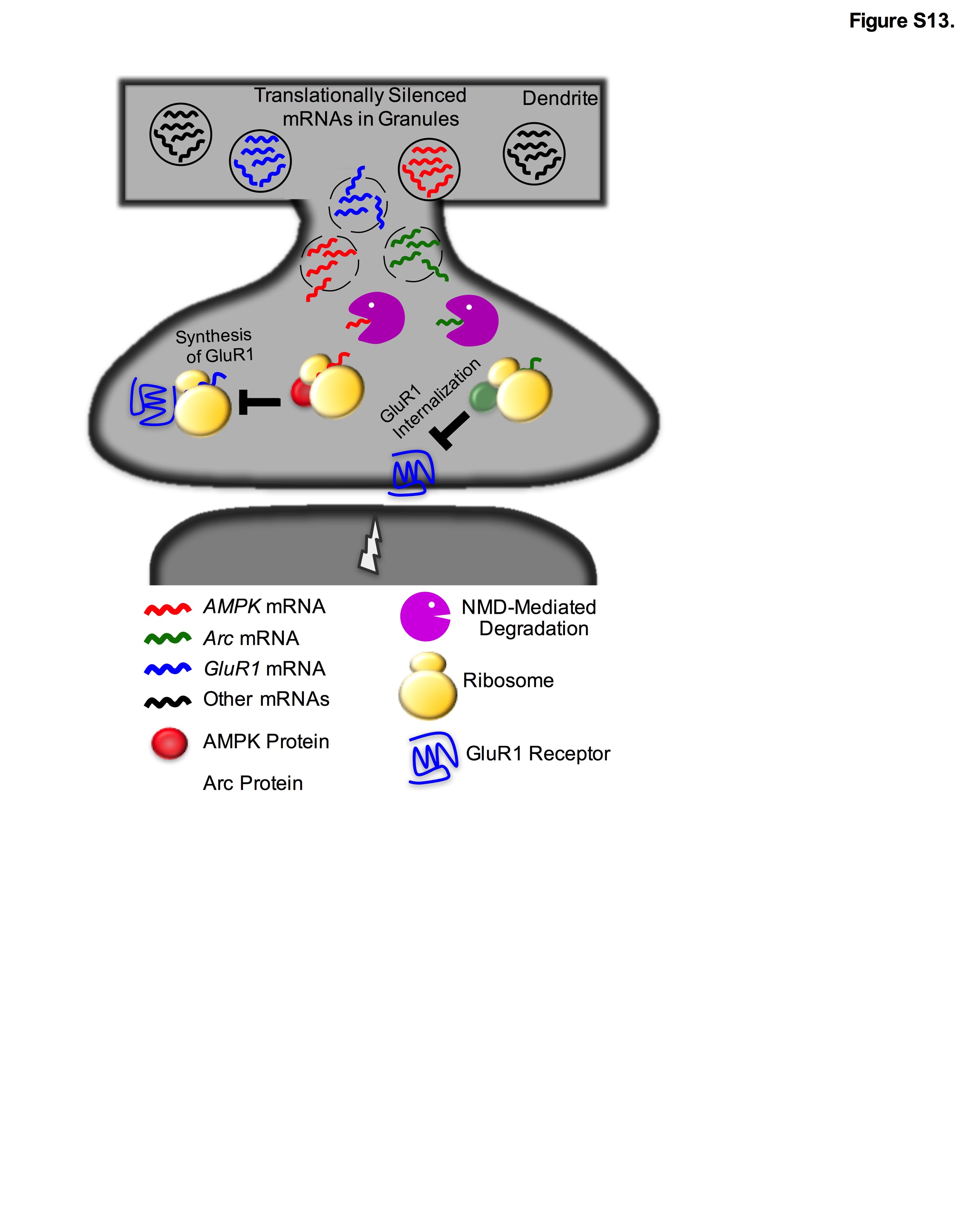

**Figure S13. Schematic model for compartmentalized NMD in dendrites.**

We establish that NMD functions in dendrites to modulate local GluR1 levels, via increased internalization and decreased local synthesis of GluR1 in dendrites. We mechanistically report that NMD degrades both *Arc* and *AMPK* mRNAs within dendrites. When NMD is disrupted*,* AMPK over-accumulates and suppresses local translation of nascent GluR1. Similarly, Arc also over-accumulates and increases GluR1 internalization rates. Co-normalization of both *Arc* and *AMPK* mRNA levels resulted in the complete recovery of dendritic surface GluR1 levels in UPF2-deficient dendrites. Additionally, our data also suggest that NMD activity is induced by synaptic stimulation. Together, we mechanistically outlay a role for NMD in plasticity, hippocampus-dependent learning and memory, and the local regulation of GluR1 specifically in dendrites, establishing that the NMD pathway is not just a passive surveillance pathway but is a critical, activity-dependent, regulator of synaptic function.
